# Deep phenotyping and multi-omics analyses reveal systems-wide metabolic dysregulation in a refined trisomy mouse model of Down syndrome

**DOI:** 10.64898/2026.08.26.747201

**Authors:** Muzna Saqib, Fangluo Chen, Diya K. Mistri, Laura Tan, Noelle Wright, Dylan C. Sarver, Robert A. Anders, Susan Aja, G. William Wong

## Abstract

Trisomy 21 or Down syndrome (DS) affects multi-organ systems across the lifespan. The presence of an extra chromosome, along with genome dosage imbalance due to triplicated genes, contributes to the DS phenotypes. Of the DS mouse models, few are aneuploid with a freely segregating extra chromosome. We previously showed that the aneuploid Ts65Dn mice exhibit metabolic deficits consistent with the metabolic profile of DS. However, the genotype-phenotype relationships in Ts65Dn mice are complicated by the presence of triplicated genes unrelated to human chromosome 21 (Hsa21). To address this issue, we leveraged a refined model, Ts66Yah, where the extra triplicated genes in Ts65Dn have been removed. Deep phenotyping and multi-omics analyses showed that Ts66Yah mice develop pronounced and widespread metabolic disturbances. Despite sexual dimorphism in weight gain, body temperature, lipid and lipoprotein profiles, hepatic injury and adipose fibrosis, both male and female Ts66Yah mice share a common phenotype of pronounced glucose intolerance and insulin resistance, reduced mitochondrial respiratory capacity in visceral fat, altered serum inflammatory cytokine profile, and dysregulated serum and liver metabolomes. Pan-tissue transcriptomes also reveal signatures of immune activation, disrupted metabolic processes and cellular respiration, altered cytokine signaling, enhanced oxidative stress, and extracellular matrix remodeling. These combined changes across tissues disrupt metabolic homeostasis more severely in Ts66Yah than in Ts65Dn mice. Several phenotypes, including glucose intolerance, insulin resistance, tissue fibrosis, and oxidative stress were further exacerbated by an obesogenic diet.

This foundational data establishes Ts66Yah as a valuable reference model for the mechanistic and comparative study of metabolic dysfunction in DS.

## INTRODUCTION

Trisomy 21 or Down syndrome (DS) is the most common survivable aneuploidy, affecting ∼1/800 live births (1). The presence of an extra human chromosome 21 (Hsa21) increases the expression dosage of ∼235 triplicated protein-coding genes, with cascading effects that impact many organ systems across the lifespan (2–6). Consequently, individuals with DS are presented with cognitive deficits, craniofacial dysmorphology, hypotonia, and the development of Alzheimer’s disease (AD)-like pathology in mid-life (7, 8). A significant number of individuals with DS also have additional clinical issues that include heart abnormalities, auditory and visual impairment, reduced bone mass, gastrointestinal diseases, and leukemia (7, 8).

Of the myriad deficits, DS-associated metabolic dysfunction has historically been understudied but is increasingly being appreciated as a major co-occurring condition across the lifespan in individuals with DS. It is known that adolescents and adults with DS have a much higher incidence of obesity, insulin resistance, type 2 diabetes, and dyslipidemia (9–18). One of the largest retrospective studies found that the median age at diabetes diagnosis was 15 years earlier in individuals with DS, and diabetes was more than four times more common in children and young adults with DS than in individuals without DS (12). There was also an increased incidence of obesity in children and young adults with DS, with rates increasing over time (12). Despite extensive documentation (9–11, 19–21), the underlying cause of DS-associated metabolic deficits is only beginning to be unraveled.

Human clinical studies have shown that trisomy 21 affects food intake, fat mass, physical activity, and energy expenditure in adolescents or adults with DS (22–30), as well as mitochondrial function in skeletal muscle (31). How much of this is due to changes in lifestyle rather than genetics remains an open question. Mechanistic studies have also highlighted cellular changes in mitochondrial morphology, dynamics, and function in cultured cells derived from DS (32–42); to what extent this is recapitulated *in vivo* remains largely unknown. These major knowledge gaps, along with the inherent limitations of human studies, have motivated us to explore the molecular and physiological underpinnings of metabolic dysfunction in DS mouse models. Although valuable, no single model recapitulates the full spectrum of DS phenotype, each has advantages and limitations (43). Of the two dozen DS mouse models that have been generated (44, 45)—each carrying a variable number of triplicated Hsa21 gene orthologs—most are segmental duplication models, and only four are considered aneuploid models with a freely segregating extra chromosome: Tc1, TcMAC21, Ts65Dn, and Ts66Yah (46–49).

The Tc1 model carries part of the Hsa21 but harbor numerous deletions, mutations, duplications, and structural rearrangements, and is missing >50 protein-coding genes (46, 50). This model has been largely abandoned due to its high degree of mosaicism, namely that the Hsa21 is present in zygotes but is lost randomly from cells during development, resulting in mice that are genetically different from each other. To overcome this issue, a new trans-chromosomic model, TcMAC21, was generated that carries a non-mosaic and near complete Hsa21q (48). Contrary to our expectation, we discovered that TcMAC21 mice are hypermetabolic, showing enhanced thermogenesis, with greatly elevated mitochondrial respiration and energy expenditure; they are markedly leaner despite hyperphagia, along with heightened insulin sensitivity (51). These phenotypes are fundamentally inconsistent with the metabolic profile of DS, making TcMAC21 an inappropriate model to study DS-associated metabolic dysfunction. It is likely that the abnormal interactions between human and mouse proteins, combined with the presence of over 400 non-coding human genes (miRNA and lncRNA)—many of which are not well conserved between human and mouse— have a profound and unintended impact on the mouse genome, pan-tissue transcriptomes, and physiology (51).

The Hsa21 gene orthologs in mice are spread across three syntenic regions on mouse chromosomes 16 (Mmu16), 10 (Mmu10), and 17 (Mmu17), with the majority located on Mmu16. A reciprocal translocation event between mouse chromosome 16 (Mmu16) and Mmu17 gives rise to a freely segregating marker chromosome, Ts(17^16^), in the Ts65Dn trisomy mice; this widely used model is triplicated for ∼103 Hsa21 gene orthologs found on Hsa21 (47, 52, 53); they lack ∼70 Hsa21 gene orthologs (54). We previously showed that Ts65Dn mice fed a standard chow exhibit only mild phenotypes with glucose intolerance (55). Metabolic homeostasis only becomes markedly disrupted by an obesogenic high-fat diet, as evidenced by dyslipidemia, impaired systemic insulin sensitivity, reduced mitochondrial activity, and elevated fibrotic and inflammatory gene signatures in the liver and adipose tissue. Disrupted gene connectivity and pathways in liver and adipose tissues contribute to impaired glucose and lipid metabolism in Ts65Dn mice (55). Our findings in the Ts65Dn mice are consistent with the general metabolic profile of DS. One important caveat and limitation of the Ts65Dn model is that it also carries an extra 41 triplicated protein-coding genes and 5 lncRNA genes (from the centromeric region of Mmu17) unrelated to Hsa21 (56, 57). This complicates the genotype-phenotype relationships in Ts65Dn mice (49, 58).

To circumvent the limitations associated with Ts65Dn and TcMAC21 models, we recently performed an in-depth metabolic analysis of Dp(16)1Yey/+ (abbreviated Dp16) mice, another widely used DS model (59). The Dp16 mice do not carry an extra chromosome but instead carry a duplicated segment of Mmu16 syntenic to Hsa21, with 115 triplicated Hsa21 gene orthologs (59). This model was generated by precise Cre/LoxP-mediated recombineering method and therefore carries no extra triplicated genes unrelated to Hsa21. Despite numerous sex differences, Dp16 male and female mice share a core phenotype of pronounced glucose intolerance, insulin resistance, impaired lipid clearance, and dyslipidemia; these phenotypes are further exacerbated by a high-fat diet (60). Metabolic dysfunction in Dp16 mice is underpinned by biochemical, metabolomic, and transcriptomic signatures of tissue fibrosis, oxidative stress, a proinflammatory state, and disrupted metabolism and mitochondrial function. This collective phenotype aligns well with the metabolic profile of DS. One limitation, however, is that Dp16 mice do not carry a freely segregating extra chromosome. Recent studies have suggested that the extra chromosome itself can contribute to DS phenotypes independent of the triplicated gene content (61).

In the present study, we aim to leverage a newly generated trisomy model, Ts66Yah, to simultaneously address the limitations found in Ts65Dn and Dp16 mouse models. The Ts66Yah is a refined model of Ts65Dn with improved construct and validity, in which all the extra triplicated genes unrelated to Hsa21 (spanning 6.4 Mb) have been removed by CRISPR/Cas9 technology (49). This model would help clarify the potential confounding effects of the extra triplicated non-Hsa21 gene orthologs on metabolic phenotypes. Dp16 and Ts66Yah mice carry 115 and 103 triplicated Hsa21 gene orthologs, respectively; both have approximately similar triplicated gene content (103 are identical) but only Ts66Yah mice carry an extra chromosome. The comparison of Ts66Yah with our published data on Dp16 mice would give insights into metabolic phenotypes that are potentially due to the presence of an extra chromosome.

Our comprehensive phenotyping and multi-omics analyses showed that Ts66Yah mice exhibit pronounced systems-wide metabolic perturbations that are more severe than in the Ts65Dn mice we previously studied (55). This suggests that the extra triplicated genes unrelated to Hsa21 likely mask some of the metabolic phenotypes in Ts65Dn mice; their removal reveals the more accurate impact of the triplicated Hsa21 gene orthologs on systemic metabolism. Additionally, comparison of the metabolic phenotypes of Ts66Yah mice with the published Dp16 data highlighted similarities but also differences that suggest the potential contribution of an extra chromosome to some metabolic phenotypes that warrant further investigation. Given the paucity of studies on this new trisomy model, our comprehensive metabolic and molecular data lay an important foundation and establish Ts66Yah mice as a valuable reference model for mechanistic and comparative studies of metabolic dysregulation in DS.

## RESULTS

### Increased expression dosage of triplicated genes across tissues

The Ts66Yah mice carry 93 triplicated protein-coding genes (**Fig. 1A**). We performed bulk RNA sequencing to determine which among the 93 triplicated Hsa21 gene orthologs on Mmu16 are expressed in six major metabolic tissues: gonadal white adipose tissue (gWAT; visceral fat), inguinal white adipose tissue (iWAT; subcutaneous fat), brown adipose tissue (BAT), skeletal muscle (gastrocnemius), and hypothalamus. We showed that the majority of the triplicated genes are expressed at the expected higher dosage (∼1.5-fold or higher) across tissues, and their expression is regulated in a tissue- and sex-specific manner (**Fig. 1B,C**). Only three triplicated Hsa21 gene orthologs (*Mrap*, *Chaf1b*, and *Sh3bgr*) are expressed at levels significantly below that of euploid controls (**Fig. 1B**), broadly consistent with recent findings indicating minimal dosage compensation in DS (62). Among the six tissues, skeletal muscle expresses the largest number of triplicated Hsa21 gene orthologs, whereas the hypothalamus has the largest number of shared triplicated genes between males and females (**Fig. 1C**). Visceral (gWAT) and subcutaneous (iWAT) fat depots have the lowest number of shared differentially expressed triplicated genes between sexes (**Fig. 1C**). Unlike the Ts65Dn model, the 46 triplicated genes on Mmu17 centromeric region unrelated to Hsa21 have been removed in Ts66Yah (49). Consequently, the expression of these genes was normalized to that of euploid controls (**Fig. 1 – figure supplement 1 and source data 1**). Together, these data indicate sexually dimorphic expression of the triplicated Hsa21 gene orthologs across metabolic tissues in Ts66Yah mice.

**Figure 1.**
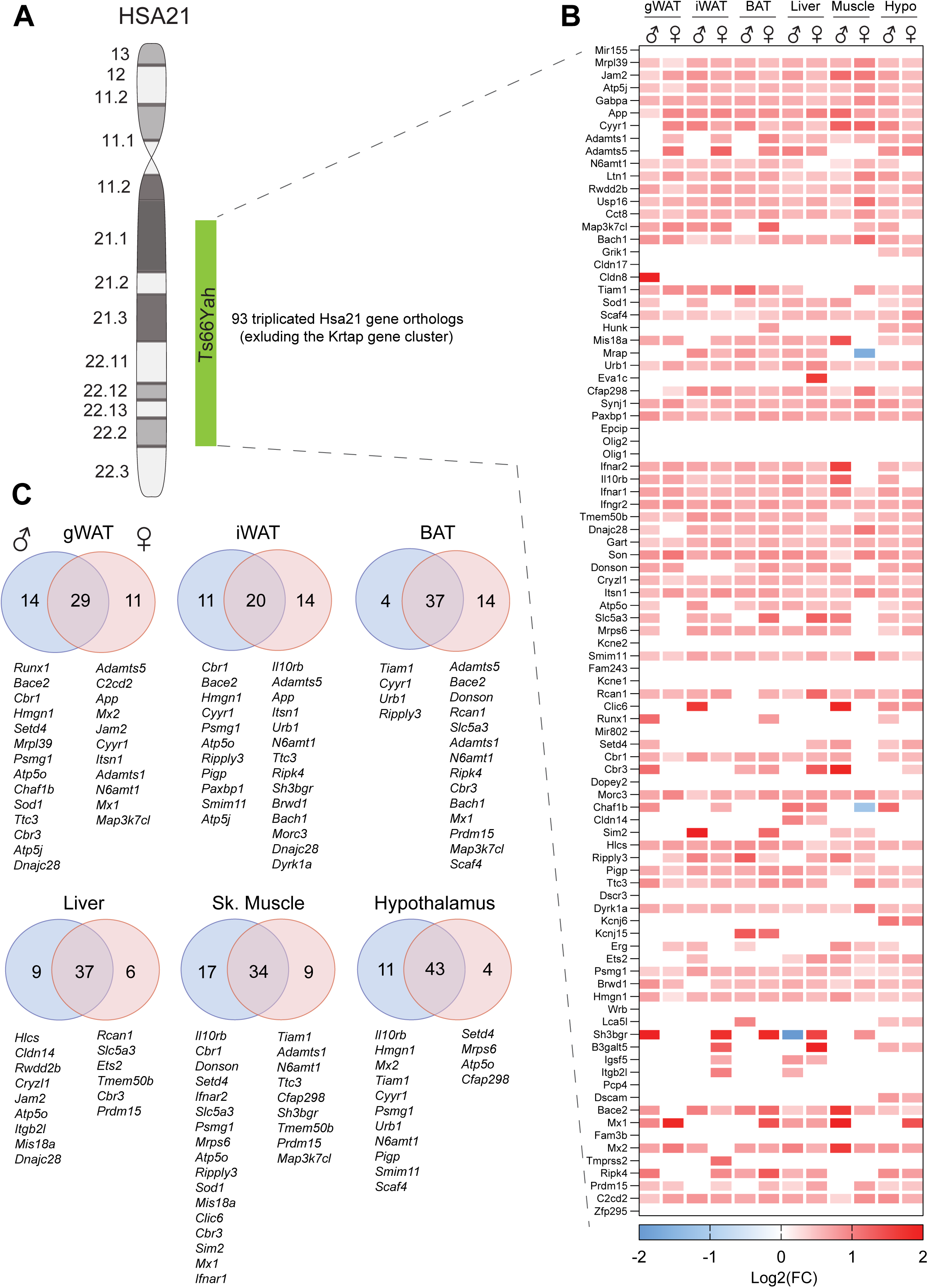
Increased gene expression dosage of the triplicated Hsa21 gene orthologs on mouse chromosome 16 (Mmu16) in Ts66Yah male and female mice. **(A)** Graphical representation of human chromosome 21 (Hsa21) and the syntenic Mmu16 segment that is trisomic in Ts66Yah mice. **(B)** Global view of the expression of 93 triplicated Hsa21 gene orthologs on Mmu16 in gonadal white adipose tissue (gWAT), inguinal white adipose tissue (iWAT), interscapular brown adipose tissue (BAT), skeletal muscle (gastrocnemius), and hypothalamus. Red denotes transcript that is expressed at >1.5-fold the WT level, whereas blue denotes transcript that is expressed at significantly lower level compared to euploid (Eu) control. The Krtap gene cluster (23 Krtap genes) located between Cldn8 and Tiam1is not shown. **(C)** Overlap analysis showing differentially expressed Hsa21 gene orthologs that are shared between males and females across six tissues. The criteria for differentially expressed genes (DEGs) is log2(FC) > 0 with padj (FDR) < 0.05. *n* = 6 RNA samples per genotype per sex per tissue-type.

### Sexual dimorphism in body weight, fecal output, and body temperature in Ts66Yah mice

We next examined the impact of triplicated gene expression on systemic metabolism under the basal state when mice were fed standard chow. We tracked the body weights of male and female mice over 10 weeks. The body weights of Ts66Yah male mice from 11 to 19 weeks of age were not different from WT controls (**Fig. 2A**). Despite similar body weight, body composition analysis at 11 weeks old revealed that Ts66Yah male mice have higher fat mass and reduced lean mass (**Fig. 2B**). At 20 weeks old, the body weights, as well as heart and kidney weights, were lower in Ts66Yah males relative to euploid controls (**Figure 2 – figure supplement 1**). In contrast to males, Ts66Yah females consistently have higher average body weights from 10 to 21 weeks of age (**Fig. 2C**). Increased body weight in Ts66Yah females was due to increased fat and lean mass (**Fig. 2D**). At 21 weeks old, the body weights and the absolute weights of gWAT and liver were higher in Ts66Yah females, whereas the relative weights of heart and kidney were lower compared to euploid controls (**Figure 2 – figure supplement 1**).

**Figure 2.**
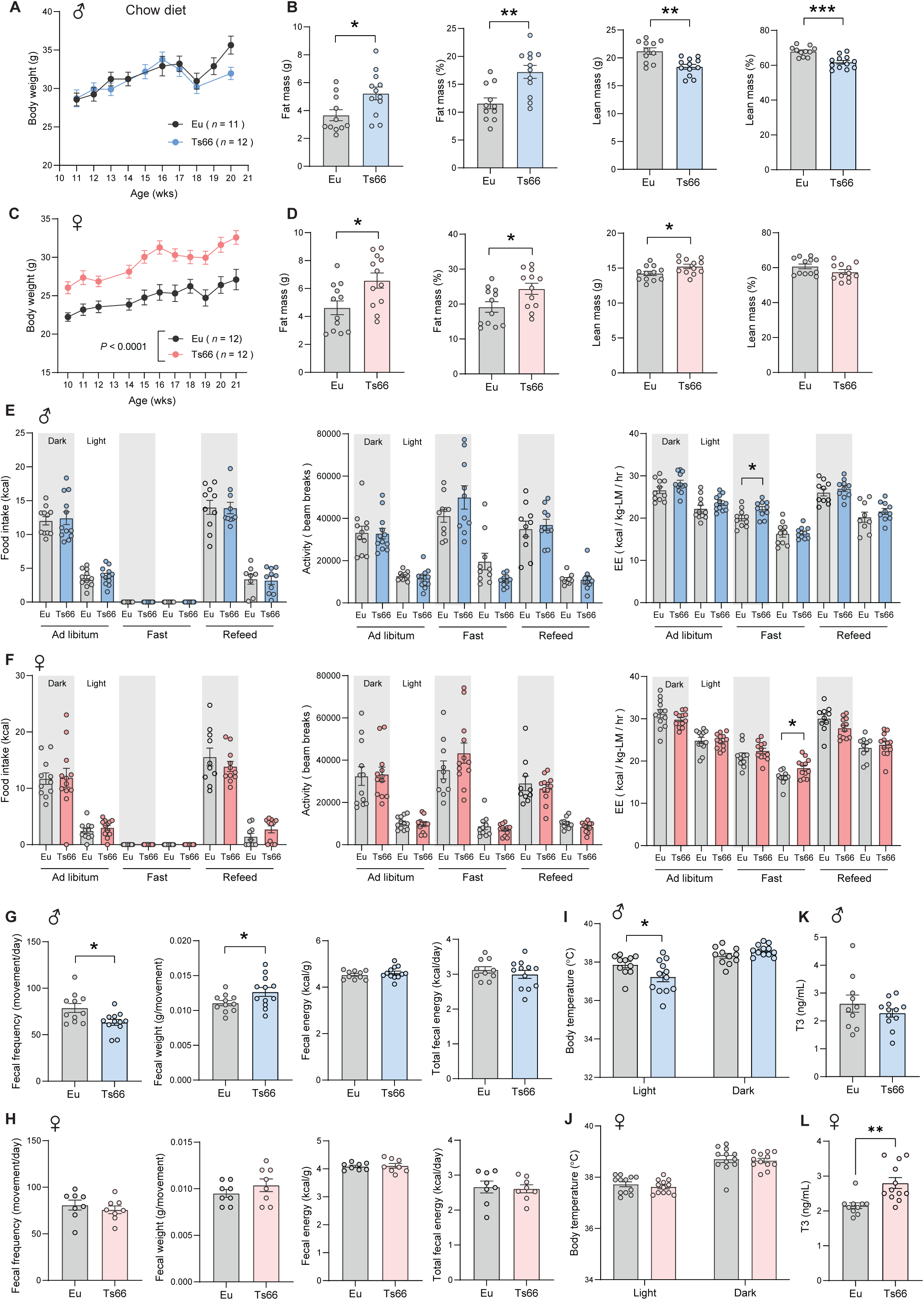
Sexually dimorphism in body weight, body temperature, fecal output, and T3 levels in Ts66Yah mice. **(A)** Body weight of chow-fed male Ts66Yah (abbreviated Ts66) mice and Euploid (Eu) controls over time. **(B)** Absolute and relative (% of body weight) fat and lean mass in male mice (Eu = 11; Ts66 = 12). **(C)** Body weight of chow-fed female Ts66 and euploid controls over time. **(D)** Absolute and relative (% of body weight) fat and lean mass in female mice (Eu = 12; Ts66 = 12). **(E-F)** Food intake, total physical activity level, and energy expenditure of male (E) and female (F) Ts66 and euploid controls across the circadian cycle (light and dark) and metabolic states (*ad libitum* fed, fast, refeed). Sample size for male (Eu = 11; Ts66 = 9-12) and female (Eu = 12; Ts66 = 10-12) mice. **(G-H)** Fecal frequency, average fecal weight, and fecal energy content (per gram and total) in male (G) and female (H) Ts66 and euploid controls. Sample size for male (Eu = 10-11; Ts66 = 12) and female (Eu = 8; Ts66 = 8) mice. **(I-J)** Body temperature in the light and dark cycle of male (I) and female (J) Ts66 and euploid controls. Sample size for male (Eu = 11; Ts66 = 12) and female (Eu = 12; Ts66 = 12) mice. **(K-L**) Serum triiodothyronine (T3) levels in male (K) and female (L) Ts66 and euploid controls. Sample size for male (Eu = 10; Ts66 = 12) and female (Eu = 10; Ts66 = 12) mice. All data are presented as mean ± SEM. \**P*<0.05; *** *P*<0.001. For body weight over time, data were analyzed by two-way ANOVA with Sidek post hoc tests; other data were analyzed by two-tailed student *t*-Test.

Indirect calorimetry analysis was used to determine food intake, physical activity, metabolic rate (VO_2_), and energy expenditure across the circadian cycle (light and dark phase) and metabolic states (*ad libitum* fed, fasted, and refed). Most of the parameters measured were not different between genotypes of either sex (**Fig. 2E,F**). However, both Ts66Yah males and females had higher energy expenditure during fasting, albeit at different phases of the circadian cycle. We performed ANCOVA analyses (using lean mass as a covariate of energy expenditure) to ensure that energy expenditure when normalized to lean mass does not lead to overestimation (63). Except during fasting, both types of analyses indicated no differences in energy expenditure between genotypes of either sex across the circadian cycles and metabolic states (**Figure 2 – figure supplement 2**).

Altered gastrointestinal function has been documented in individuals with DS (64). To rule out any potential differences in nutrient absorption in the intestine, we measured fecal output, frequency, and weight, as well as the fecal energy content by fecal bomb calorimetry. Interestingly, Ts66Yah males had reduced number of bowel movements (i.e., lower fecal frequency) but higher fecal output per bowel movement, resulting in no net changes in total fecal energy content (**Fig. 2G**). In contrast to males, none of the fecal parameters were different between genotypes in females (**Fig. 2H**).

We also measured the body temperature of male and female mice across the circadian cycle. We noted that Ts66Yah males, but not females, had lower body temperature during the light cycle when the animals are generally resting and less active (**Fig. 2I-J**). This prompted us to assess circulating triiodothyronine (T3), testosterone, and estradiol levels, as these hormones can affect body temperature. Contrary to expectation, serum T3 levels were not different between genotypes in male mice, but were higher in Ts66Yah females relative to euploid controls (**Fig. 2K,L**). Circulating sex hormones were not different between genotypes of either sex (**Figure 2 – figure supplement 3**). Together, these data reveal sex differences in body weight, fecal output, and body temperature in Ts66Yah mice. The higher body weights in Ts66Yah females are not due to altered caloric intake, physical activity, and energy expenditure, as these parameters are largely not different between genotypes.

### Glucose intolerance, insulin resistance, and dyslipidemia in Ts66Yah mice

To determine baseline insulin, glucose, and lipid profiles, we measured blood glucose, serum insulin, triglyceride (TG), cholesterol, non-esterified free fatty acids (NEFA), and β-hydroxybutyrate (BHB; ketone) in overnight fasted (16 h) mice. The Ts66Yah males fed standard chow had similar fasting serum insulin, blood glucose, and triglyceride levels as the euploid controls, but they had lower cholesterol and higher non-esterified free fatty acid (NEFA) and ketone (β-hydroxybutyrate) levels compared to euploid controls (**Fig. 3A**). Elevated NEFA and ketone levels in Ts66Yah male mice suggest enhanced fat mobilization from the adipose tissue and greater hepatic fat oxidation in the fasted states. In contrast, fasting insulin, glucose, triglyceride, cholesterol, NEFA, and ketone levels were not different between genotypes in female mice (**Fig. 3B**).

**Figure 3.**
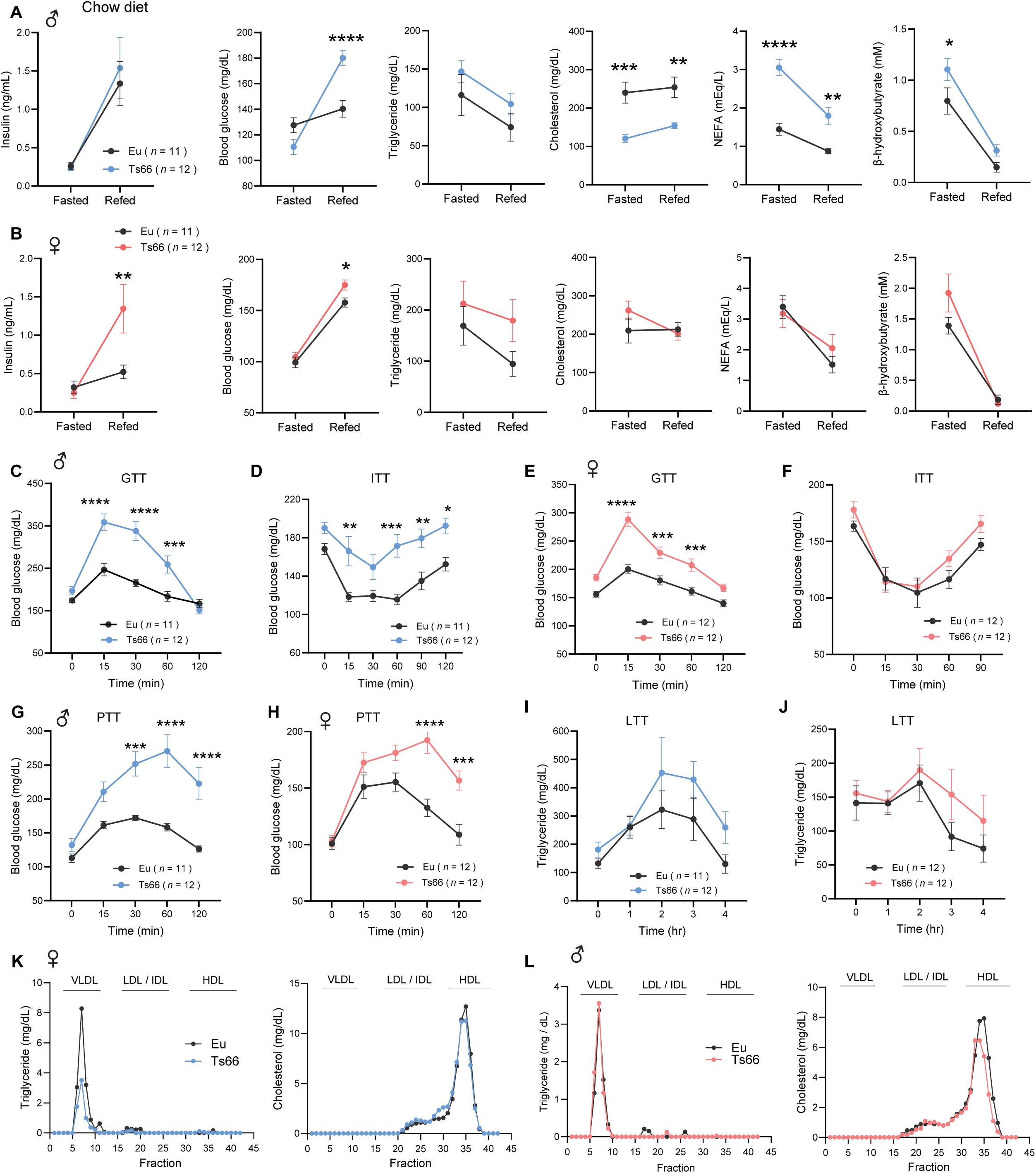
Glucose intolerance, insulin resistance, and dyslipidemia in Ts66Yah mice. **(A-B)** Overnight fasted and refed blood glucose, serum insulin, triglyceride, cholesterol, non-esterified free fatty acids (NEFA), and β-hydroxybutyrate (ketone) levels in male (A) and female (B) Ts66 mice and euploid (Eu) controls. Sample size for chow-fed male (Eu = 11; Ts66 = 12) and female (Eu = 11; Ts66 = 12) mice. (**C-F)** Impaired glucose tolerance as determined by the glucose tolerance test (GTT) in male (C) and female (E) Ts66 compared to euploid controls. Impaired insulin sensitivity as determined by the insulin tolerance test (ITT) in male (D) and female (F) Ts66 compared to euploid controls. Sample size for chow-fed male (Eu = 11; Ts66 = 12) and female (Eu = 12; Ts66 = 12) mice. **(G-H)** Impaired pyruvate tolerance as determined by the pyruvate tolerance test (PTT) in male (G) and female (H) Ts66 mice compared to euploid controls. Sample size for chow-fed male (Eu = 11; Ts66 = 12) and female (Eu = 12; Ts66 = 12) mice. **(I-J)** Triglyceride clearance in response to lipid gavage as determined by the lipid tolerance test (LTT) in male (I) and female (J) Ts66 compared to euploid controls. Sample size for chow- fed male mice (Eu = 11; Ts66 = 12) and female mice (Eu = 12; Ts66 = 12). **(K-L)** Pooled mouse sera from male (K) and female (L) Ts66 and euploid control were fractionated by fast protein liquid chromatography (FPLC), and the triglyceride and cholesterol content of each fraction was quantified. Fractions corresponding to very-low density lipoprotein (VLDL), low-density lipoprotein (LDL), intermediate-density lipoprotein (IDL), and high-density lipoprotein (HDL) are indicated. All data are presented as mean ± SEM. * *P*<0.05; ** *P*<0.01; *** *P*<0.001; **** *P*<0.0001. For all tolerance tests, data were analyzed by 2-way ANOVA with Sidek post hoc tests.

In response to refeeding, Ts66Yah males had much higher blood glucose despite similar insulin levels as the euploid controls (**Fig. 3A**), indicative of an insulin resistant state. In the fed state, insulin suppresses adipose lipolysis. In response to refeeding, Ts66Yah males were unable to fully suppress lipolysis, resulting in higher NEFA levels compared to euploid controls, again consistent with an insulin resistant state. Circulating ketones, produced by the liver, were fully suppressed by refeeding. Cholesterol levels remained low in Ts66Yah males regardless of metabolic states. In contrast to males, refeeding markedly elevated insulin levels in Ts66Yah females; despite this, blood glucose is still higher than the euploid controls, indicative of impaired insulin action (**Fig. 3B**). In the refed state, serum triglyceride, cholesterol, NEFA, and ketone levels were not different between genotypes in female mice.

Fasting and refeeding profiles suggest insulin resistance in Ts66Yah male and female mice. To further assess glucose metabolism in these mice, we performed glucose and insulin tolerance tests to determine the rate of glucose clearance in response to glucose or insulin injection. Both Ts66Yah males and females showed impaired glucose clearance after glucose loading (**Fig. 3C,E**). To confirm that Ts66Yah mice have impaired insulin action, we directly assessed insulin sensitivity via insulin tolerance test (ITT). The rate of glucose clearance in response to insulin injection was markedly impaired in Ts66Yah males but not females (**Fig. 3D,F**). To assess hepatic insulin sensitivity, we performed pyruvate tolerance tests (PTT). In the fasted state, pyruvate will be used as a substrate for hepatic gluconeogenesis, and insulin suppresses hepatic glucose production. Thus, the rise in blood glucose following pyruvate injection in fasted mice provides an indication of hepatic insulin action. We observed that Ts66Yah males and females have markedly higher blood glucose following pyruvate administration (**Fig. 3G,H**), indicative of hepatic insulin resistance. Together, these data strongly suggest an insulin resistance phenotype in Ts66Yah mice.

We next assessed whether Ts66Yah mice have altered lipid handling capacity by performing a lipid tolerance test (LTT). The rate of triglyceride clearance in response to an acute lipid load appeared to be blunted but was not significantly different between genotypes of either sex (**Fig. 3I,J**). To determine whether Ts66Yah mice have altered lipoprotein profiles, we subjected pooled sera to FPLC fractionation followed by the quantification of triglyceride and cholesterol levels in each fraction. Ts66Yah males but not females had lower triglyceride in the VLDL fraction (**Fig. 3I**). HDL-cholesterol level was not different in males, but modestly lower in Ts66Yah females (**Fig. 3K,L**). Altogether, these data indicate that both male and female Ts66Yah mice developed pronounced insulin resistance, glucose and pyruvate intolerance, and dyslipidemia.

### Altered tissue mitochondrial respiratory capacity in Ts66Yah mice

Impaired mitochondrial function has been well documented in cultured cells derived from DS (32–42), and this can contribute to metabolic deficits in Ts66Yah mice. To assess mitochondrial function, we performed high-resolution mitochondrial respirometry analysis on visceral fat, subcutaneous fat (iWAT), brown fat (BAT), liver, and skeletal muscle (gastrocnemius). Maximal mitochondrial respiration through complex I (CI), CII, and CIV was determined with the use of specific electron donors: NADH (CI substrate), succinate (CII substrate), and TMPD (CIV substrate). ATP synthase (CV) activity was determined by the acidification rate of ATP hydrolysis. In both male and female Ts66Yah mice, we observed significantly reduced mitochondrial respiratory capacity in gWAT, but not in iWAT, BAT, or liver (**Fig. 4**); ATP synthase activity in these tissues, however, was not different between genotypes of either sex. Interestingly, Ts66Yah females, but not males, had higher mitochondrial respiratory capacity and ATP synthase activity in skeletal muscle. Together, these data suggest impaired mitochondrial function in visceral fat depot and potential sex-dependent mitochondrial compensation and adaptation in skeletal muscle of Ts66Yah mice.

**Figure 4.**
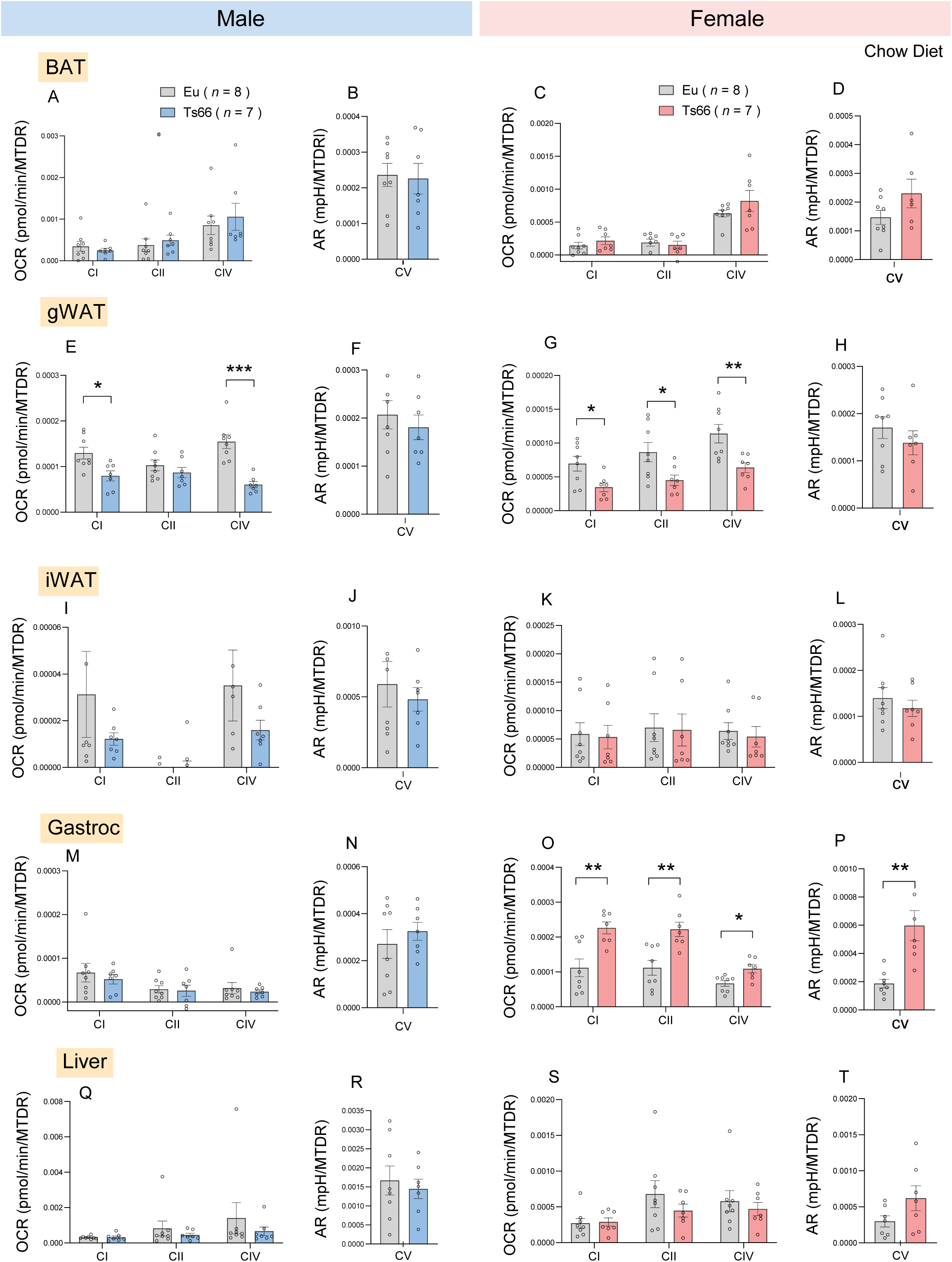
Altered mitochondrial respiratory capacity and ATP synthase activity in Ts66Yah mice. **(A-T)** Mitochondrial respiration through complex I (CI), CII, and CIV, as well as CV (ATP synthase) activity in BAT, gWAT, iWAT, gastroc, and liver of Tss66 male and female mice and euploid controls. All oxygen consumption rates (OCR) and Acidification Rates (AR) are normalized to mitochondrial content (based on MTDR). MTDR, MitoTracker Deep Red; BAT, brown adipose tissue; gWAT, gonadal white adipose tissue; iWAT, inguinal white adipose tissue; gastroc, gastrocnemius. Sample size for chow- fed male (Eu = 8; Ts66 = 7) and female (Eu = 8; Ts66 = 7) mice. All data are presented as mean ± SEM. * *P*<0.05; ** *P*<0.01; *** *P*<0.001 (two-tailed Student’s *t*-Test)

### Elevated liver injury marker and adipose tissue fibrosis in Ts66Yah mice

Tissue injury, fibrosis, and oxidative stress affect systemic metabolism. We therefore quantified serum ALT level, a liver enzyme whose circulating level reflects hepatic injury. Ts66Yah males appeared to have higher serum ALT levels though not significantly (**Fig. 5A**; *P* = 0.17). Ts66Yah females, in contrast, had elevated serum ALT levels indicative of hepatic injury (**Fig. 5D**). Despite higher circulating ALT levels, liver histology and pathological scoring for steatosis, cell injury, and inflammation revealed no significant differences in Ts66Yah female mice (**Fig. 5 – figure supplement 1**).

**Figure 5.**
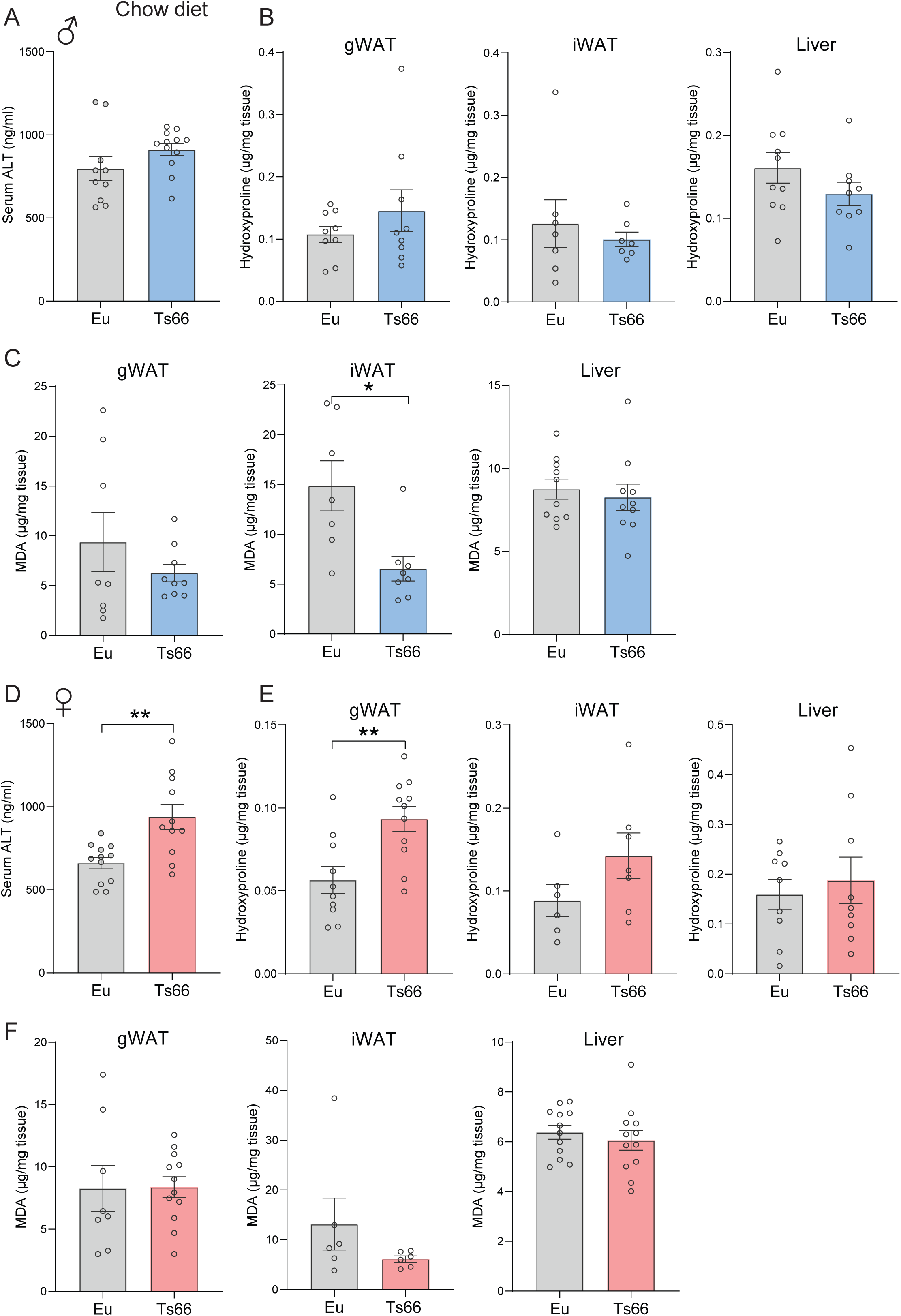
Sex-dependent vulnerability in liver injury, tissue fibrosis, and oxidative stress in Ts66Yah mice. **(A-F)** Quantification of serum alanine transaminase (ALT; marker of liver injury), as well as hydroxyproline content (marker of fibrosis) and malondialdehyde (MDA; marker of oxidative stress) in gWAT, iWAT, and liver of male (A-C) and female (D-F) Ts66 mice and euploid (Eu) controls. gWAT, gonadal white adipose tissue; iWAT, inguinal white adipose tissue. Sample size for chow-fed male (Eu = 7-10; Ts66 = 8-12) and female (Eu = 7-12; Ts66 = 8-11) mice. All data are presented as mean ± SEM. * *P*<0.05; ** *P*<0.01 (two-tailed Student’s *t*-Test)

Tissue hydroxyproline (derived from collagen) is a marker of fibrosis, and tissue malondialdehyde (MDA) derived from lipid peroxidation is a marker of oxidative stress. We measured hydroxyproline and MDA levels in gWAT, iWAT, and liver. While Ts66Yah males showed no differences in hydroxyproline across tissues (**Fig. 5B**), Ts66Yah females had elevated hydroxyproline levels in gWAT (**Fig. 5E**), indicative of a greater degree of fibrosis in the visceral fat depot. Interestingly, Ts66Yah males, but not females, had lower MDA levels in iWAT (**Fig. 5C,F**), suggesting lower oxidative stress in the subcutaneous fat depot. Together, these data suggest sex-dependent vulnerability to hepatic injury and adipose fibrosis in Ts66Yah mice.

### A systemic immune activation profile in Ts66Yah mice

Systemic low-grade inflammation is a frequent hallmark of metabolic dysfunction (65). We therefore quantified circulating proinflammatory cytokine levels in Ts66Yah mice. Relative to euploid controls, both male and female Ts66Yah mice had elevated serum IL-2, IL-5, and IL-10 levels (**Fig. 6A,B**). Serum IL-1β levels, however, were lower in Ts66Yah females. IL-2 and IL-5 are well established markers of immune activation, whereas the proinflammatory role of IL-10 is context-dependent (66). These data suggest a chronic immune activation profile and low-grade inflammation in Ts66Yah mice.

**Figure 6.**
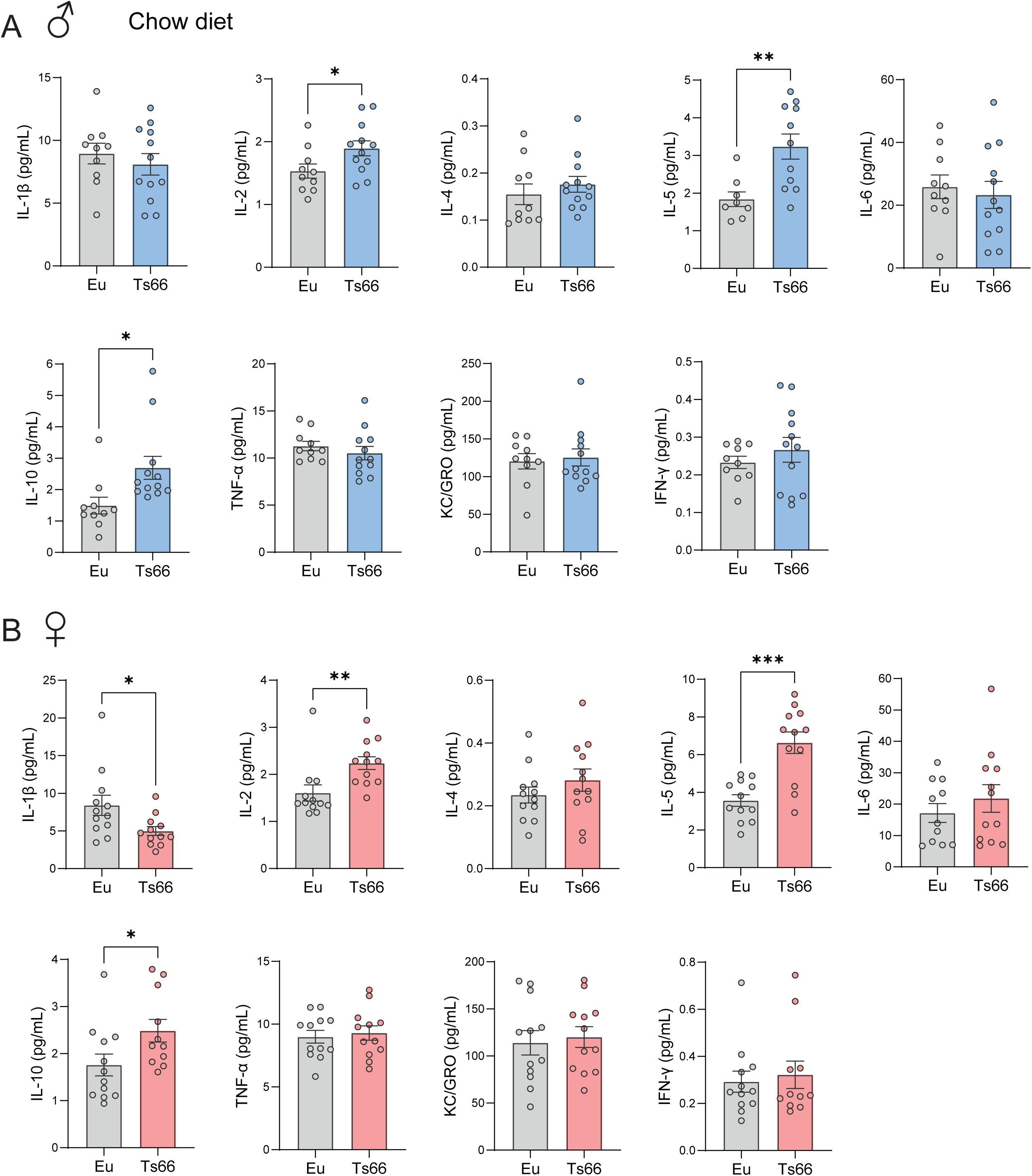
Elevated systemic immune activation in Ts66Yah mice. **(A-B)** Multiplex profiling of serum IL-1β, IL-2, IL-4, IL-5, IL-6, IL-10, TNF-α, KC/GRO (also known as CXCL1), and INF-γ in male (A) and female (B) Ts66 mice and euploid (Eu) controls. Sample size for chow-fed male (Eu = 10; Ts66 = 12) and female (Eu = 12; Ts66 = 12) mice.

### Remodeling of hepatic and serum metabolome in Ts66Yah mice

Given the systemic metabolic disturbances seen in Ts66Yah mice, we performed untargeted metabolomic analyses to further assess possible changes in liver and serum metabolomes. A total of 1873 differential metabolites were identified from the 48 serum and liver samples (**Fig. 7 – figure supplement 1; Fig. 7 – source data 1-4**). Partial Least Squares Discriminant Analysis (PLS-DA) indicated that liver and serum metabolomes of Ts66Yah males and females are clearly distinguishable from their euploid controls (**Fig. 7A,B**). In general, there are more differential metabolites up- and down-regulated in Ts66Yah males than females (**Fig. 7C**). When comparing the differential metabolites found in liver and serum, there appeared to be limited overlap between the two compartments in both male and female Ts66Yah mice (**Fig. 7D**). We observed major sex differences in the differential metabolites found in liver and serum. There were 39 differential liver metabolites shared between Ts66Yah males and females, and 127 differential serum metabolites shared between the sexes (**Fig. 7E**). The majority of differential metabolites found in liver and serum, however, were not shared between the sexes.

**Figure 7.**
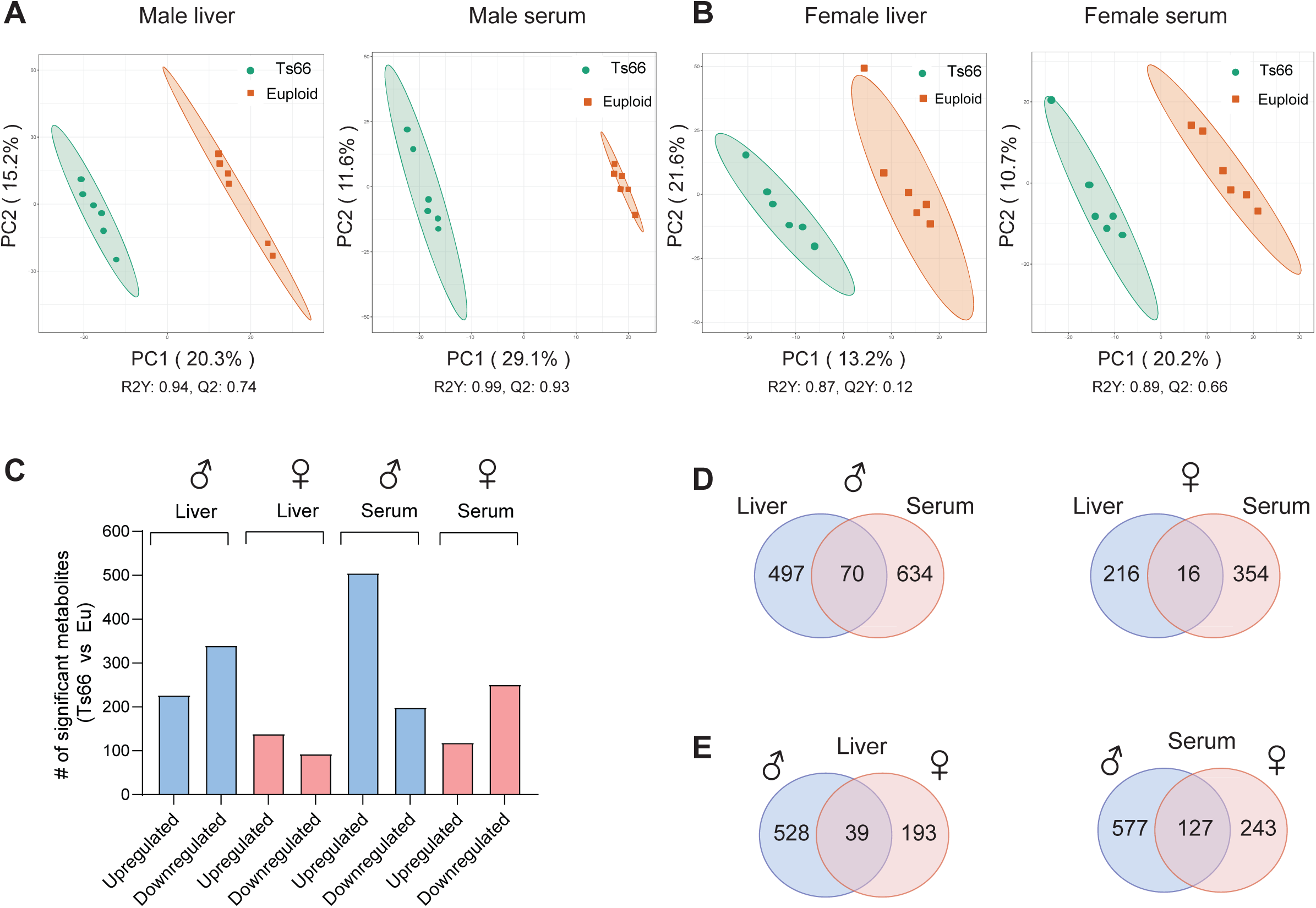
Altered liver and serum metabolome in Ts66Yah mice. **(A-B)** Partial least squares discrimination analysis (PLS-DA) of liver and serum metabolites of chow-fed male and female Ts66 mice and euploid controls. *N* = 6 samples per genotype per sex. **(C)** Total number of significant metabolites that are upregulated and downregulated in the liver and serum of male and female Ts66 mice. **(D)** Venn diagram of differential metabolites shared between liver and serum in Ts66 male and female mice. **(E)** Venn diagram of differential liver or serum metabolites shared between male and female Ts66 mice.

To provide greater details and insights, we highlighted some of the differential metabolites found in the liver and serum of Ts66Yah males and females (**Table 1-2**). Consistent with recent findings (60, 67), we also observed changes in hepatic bile acids content, with most of the bile acids showing reduced levels in Ts66Yah male mice. In contrast to the liver, serum bile acid levels are mostly elevated in Ts66Yah males and females. Bile acids are well known ligands for nuclear hormone receptors (e.g., FXR and TGR5), with pleiotropic roles in glucose and lipid metabolism (68, 69). Altered hepatic and circulating levels of bile acids are associated with, and may potentially contribute to, the systemic metabolic phenotypes in Ts66Yah mice. The opposing direction of change between liver (reduced) and serum (elevated) bile acid pools suggests a potential disruption in hepatic bile acid synthesis, recycling, or enterohepatic recirculation.

**Table 1.**
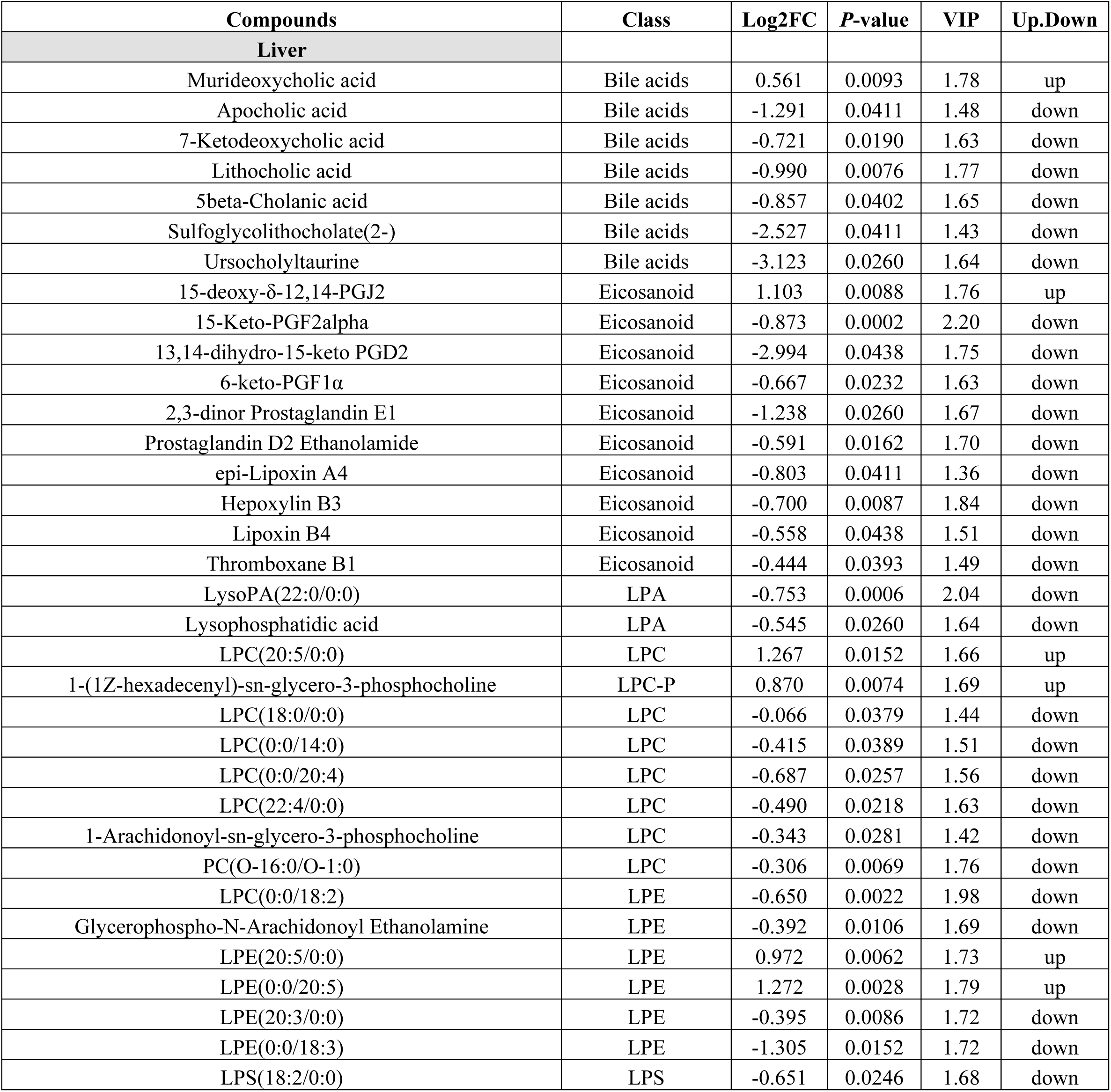
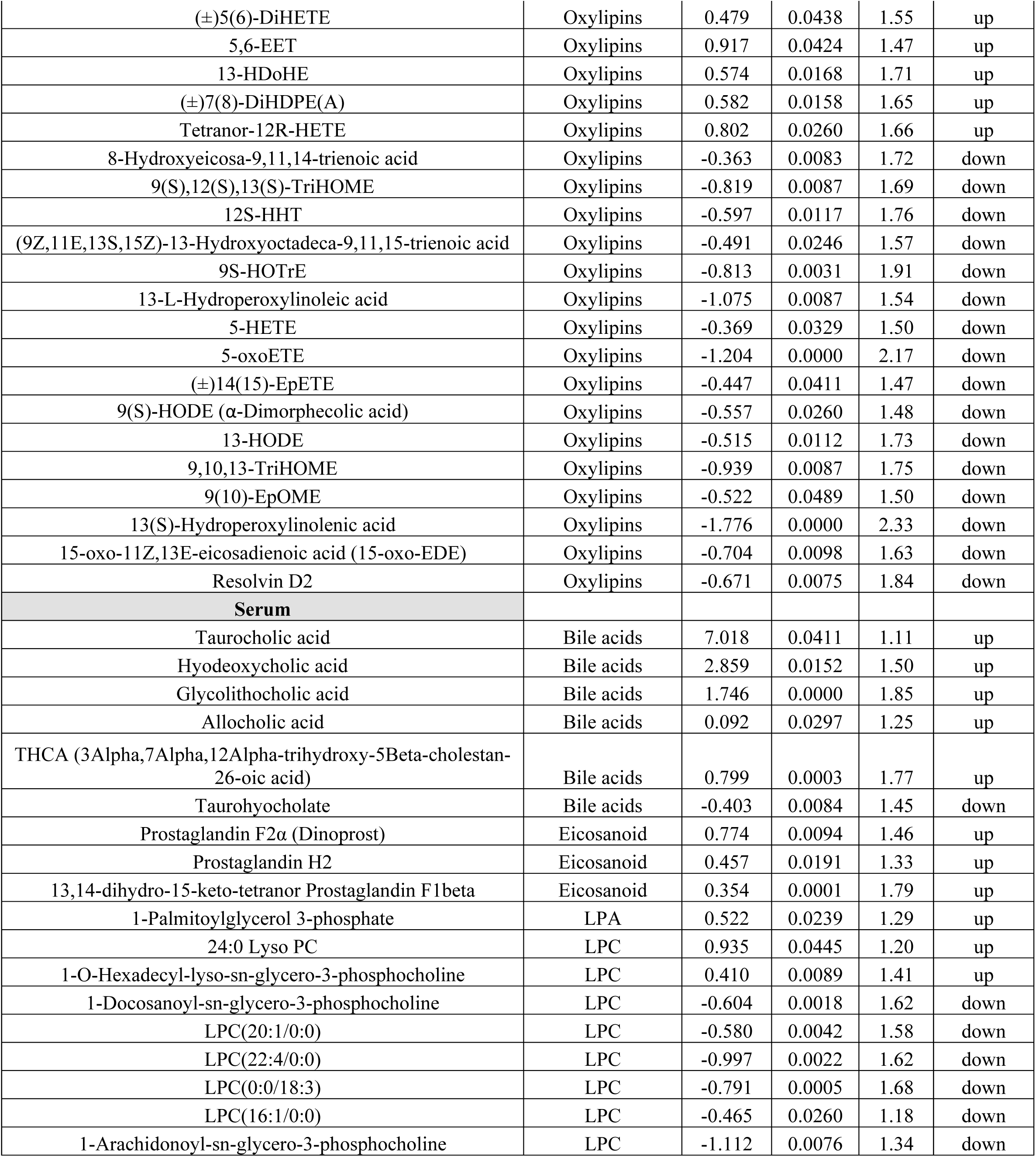
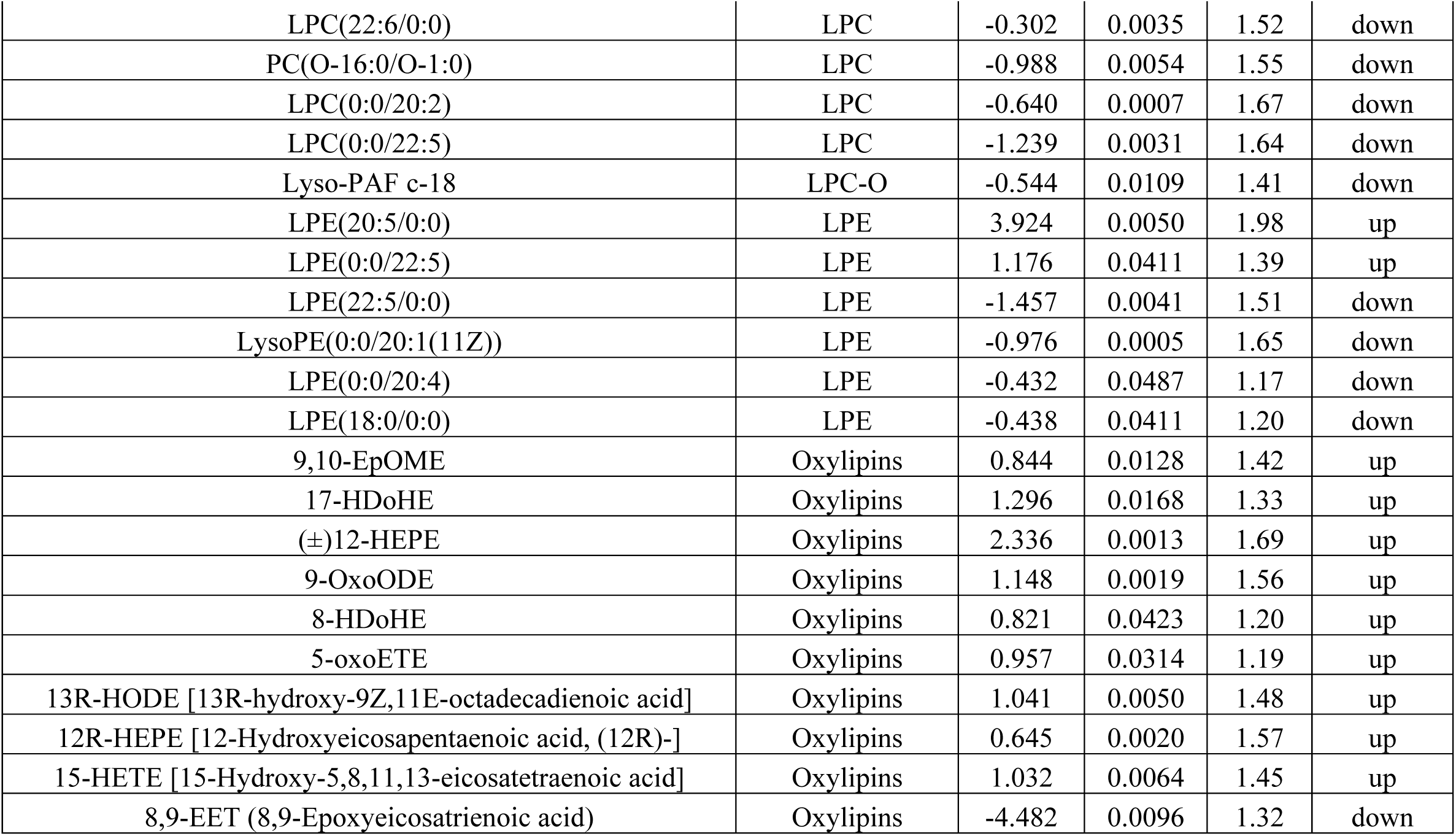
Selective differential metabolites in the liver and serum of Ts66Yah male mice. Metabolites are considered significantly different if fold change (FC) > 1.2 or < 0.833, *P*-value < 0.05, and the variable importance in projection (VIP) score is > 1. Sample size: Euploid control (*n* = 6) and Ts66Yah (*n* = 6). LPA, lyso-phosphatidic acid; LPE, lyso-phosphatidylethanolamine; LPC, lyso-phosphatidylcholine (PC); LPS, lyso-phosphatidylserine (PS); LPI, lyso-phosphatidylinositol.

**Table 2.** Selective differential metabolites in the liver and serum of Ts66Yah female mice. Metabolites are considered significantly different if fold change (FC) > 1.2 or < 0.833, *P*-value < 0.05, and the variable importance in projection (VIP) score is > 1. Sample size: Euploid control (*n* = 6) and Ts66Yah (*n* = 6). LPA, lyso-phosphatidic acid; LPE, lyso-phosphatidylethanolamine; LPC, lyso- phosphatidylcholine (PC); LPS, lyso-phosphatidylserine (PS); LPI, lyso-phosphatidylinositol.

| Compounds | Class | Log2FC | *P*-value | VIP | Up.Down |
| --- | --- | --- | --- | --- | --- |
| **Liver** |  |  |  |  |  |
| 6-keto-PGF1α | Eicosanoid | 0.554 | 0.0017 | 2.38 | up |
| 8-isoprostaglandin F1α | Eicosanoid | 0.739 | 0.0411 | 1.90 | up |
| Thromboxane B1 | Eicosanoid | 0.582 | 0.0309 | 2.02 | up |
| 11-Deoxyprostaglandin F1α | Eicosanoid | -0.750 | 0.0433 | 1.76 | down |
| 11-deoxy PGE1 | Eicosanoid | -0.100 | 0.0430 | 1.81 | down |
| 8-isoprostaglandin E2 | Eicosanoid | -0.222 | 0.0329 | 1.93 | down |
| LPC(20:5/0:0) | LPC | -0.854 | 0.0333 | 1.76 | down |
| 18:0 lyso-PC [1-Stearoylglycerophosphoserine] | LPC | -0.299 | 0.0276 | 1.94 | down |
| LPE(20:5/0:0) | LPE | -0.715 | 0.0496 | 1.79 | down |
| LysoPE(0:0/22:0) | LPE | -1.066 | 0.0043 | 2.40 | down |
| LPE(17:1/0:0) | LPE | -0.390 | 0.0244 | 1.98 | down |
| LPE(18:3/0:0) | LPE | -0.605 | 0.0412 | 1.84 | down |
| 13-HOTE | Oxylipins | 0.906 | 0.0103 | 2.26 | up |
| (±)9-HpODE | Oxylipins | 0.541 | 0.0468 | 1.89 | up |
| 5-F2t-IsoP (5-F2-isoprostane) | Oxylipins | 0.960 | 0.0411 | 1.71 | up |
| (±)14(15)-EpETE | Oxylipins | 0.613 | 0.0493 | 1.88 | up |
| 9-OxoOctadecanoic acid | Oxylipins | 0.560 | 0.0478 | 1.93 | up |
| 9S,15S-dihydroxy-5Z,13E-prostadienoic acid | Oxylipins | -0.273 | 0.0421 | 1.81 | down |
| **Serum** |  |  |  |  |  |
| Taurohyodeoxycholic acid | Bile acids | 3.124 | 0.0043 | 1.64 | up |
| Glycocholic acid | Bile acids | 0.165 | 0.0354 | 1.56 | up |
| Taurodeoxycholic acid (TDCA) | Bile acids | 4.103 | 0.0087 | 1.70 | up |
| Ursocholytaurine (TUDCA) | Bile acids | 6.553 | 0.0087 | 1.75 | up |
| 3b-Hydroxy-5-cholenoic acid | Bile acids | -0.832 | 0.0073 | 1.93 | down |
| 9(S)-HpODE | Oxylipins | 0.264 | 0.0040 | 1.94 | up |
| 8-HETrE | Oxylipins | -1.961 | 0.0087 | 2.00 | down |
| 9,10-EpOME | Oxylipins | -0.662 | 0.0388 | 1.45 | down |
| (±)4-HDHA | Oxylipins | -0.917 | 0.0150 | 1.72 | down |
| 17-HDoHE | Oxylipins | -1.979 | 0.0043 | 2.04 | down |
| (±)12-HEPE | Oxylipins | -1.566 | 0.0087 | 1.86 | down |
| 9,10-DHOME | Oxylipins | -0.750 | 0.0117 | 1.68 | down |
| 8-Hdohe | Oxylipins | -1.863 | 0.0034 | 2.03 | down |
| 5-oxoETE | Oxylipins | -1.011 | 0.0188 | 1.52 | down |
| 15(S)-Hydroxyecosatrienoic acid | Oxylipins | -1.174 | 0.0260 | 1.78 | down |
| 15-oxoETE | Oxylipins | -1.005 | 0.0152 | 1.68 | down |
| (±)18-HEPE | Oxylipins | -2.433 | 0.0260 | 1.77 | down |
| 9(S)-HODE | Oxylipins | -1.267 | 0.0245 | 1.43 | down |
| 16-Oxohexadecanoic acid | Oxylipins | -0.447 | 0.0449 | 1.45 | down |
| 9E-heptadecenoic acid | Oxylipins | -0.634 | 0.0477 | 1.47 | down |
| 15-HETE | Oxylipins | -0.830 | 0.0377 | 1.47 | down |
| 5,15-DiHETE | Oxylipins | -1.279 | 0.0411 | 1.44 | down |
| 6-trans Leukotriene B4 | Eicosanoid | -1.991 | 0.0152 | 1.78 | down |
| Leukotriene A4 | Eicosanoid | -1.773 | 0.0087 | 1.90 | down |
| S-(11-hydroxy-9-deoxy-Δ12-PGD2)-glutathione | Eicosanoid | -1.412 | 0.0022 | 2.11 | down |
| Prostaglandin F2α | Eicosanoid | -1.409 | 0.0102 | 1.89 | down |
| Prostaglandin H1 | Eicosanoid | -1.014 | 0.0260 | 1.57 | down |
| 8-iso-PGA1 | Eicosanoid | -0.405 | 0.0211 | 1.59 | down |
| 13,14-dihydro-15-keto-tetranor Prostaglandin E2 | Eicosanoid | -0.577 | 0.0157 | 1.59 | down |
| Glycerophospho-N-Arachidonoyl Ethanolamine | LPE | -0.419 | 0.0071 | 1.83 | down |
| LysoPC(18:3(6Z,9Z,12Z)) | LPC | -0.486 | 0.0260 | 1.50 | down |
| LPC P-18:0 [1-(1Z-octadecenyl)-sn-glycero-3-phosphocholine] | LPC | -0.618 | 0.0325 | 1.50 | down |
| LPE(20:3/0:0) | LPE | -0.442 | 0.0195 | 1.67 | down |
| LPI(18:1/0:0) | LPI | -1.299 | 0.0005 | 2.17 | down |

Consistent with previous findings (70), hepatic and serum kynurenine levels were also elevated in Ts66Yah mice, though only in females. In addition to bile acids, we also observed significant changes in hepatic and circulating acylcarnitine, free fatty acids, monoacylglycerols, diacylglycerides, and different classes of phospholipids—phosphatidic acid (PA), phosphatidylethanolamine (PE), phosphatidylcholine (PC), phosphatidylserine (PS) and phosphatidylinositol (PI) —in Ts66Yah males and females (**Fig. 7 - source data 1-4**), suggesting changes in lipid catabolism and remodeling. Of interest, the circulating levels of many Lyso-phospholipids (e.g., Lyso-PC, Lyso-PA, Lyso-PE), some with signaling roles (71), were also altered.

The levels of multiple oxylipins and eicosanoids—a class of lipids with pro- and anti-inflammatory roles (72), were also changed in Ts66Yah mice in a complex manner (**Table 1-2**). Some proinflammatory lipids were increased while others were reduced in a sex-dependent way, possibly reflecting ongoing systemic immune modulation. For example, several proinflammatory oxylipins and eicosanoids (e.g., 7(8)- DiHDPE, 5-oxoETE, 9,10-EpOME, 9-OxoODE, 15-HETE, Thromboxane B1,8-isoprostaglandin F1α, 9- HpODE, 5-F2t-IsoP, 9-Oxooctadecanoic acid) were elevated while some anti-inflammatory oxylipins and eicosanoids (e.g., lipoxin B4, epi-Lipoxin A4, 14(15)-EpETE, 9,10,13-TriHOME, resolving D2) were reduced. The immune profile of Ts66Yah mice, as reflected by the metabolite data, is broadly consistent with the multiplex cytokine profiling data (**Fig. 6**), as well as the transcriptomic data across tissues (gWAT, iWAT, BAT, liver, skeletal muscle) showing a heightened immune activation state (data are discussed further below). Taken together, our metabolomic analyses reveal major sex-dependent and independent remodeling of hepatic and serum metabolomes, highlighting parallel changes that are associated with, and likely also contributing directly or indirectly to, the systemic metabolic dysfunction in Ts66Yah mice.

### Transcriptomic signatures underpinning metabolic dysfunction in Ts66Yah mice

To gain mechanistic insights into molecular changes that underpin the metabolic phenotypes of Ts66Yah mice, we performed bulk RNA sequencing to assess global changes in the transcriptome and biological pathways across six metabolic tissues (**Fig. 8 – source data 1-24**). Except for the hypothalamus, Ts66Yah females have more differentially expressed genes (DEGs) across tissues compared to males (**Fig. 8A**). In Ts66Yah females, gWAT and skeletal muscle together accounted for the majority of DEGs, with the least number of DEGs seen in the hypothalamus (**Fig. 8A**). In Ts66Yah males, skeletal muscle has the highest number of DEGs, with the liver having the least DEGs. In general, there are more upregulated than downregulated DEGs across tissues, except in female skeletal muscle where there are more downregulated DEGs. Overlap analysis indicates that the majority of the DEGs are not shared between Ts66Yah males and females (**Fig. 8B**). Skeletal muscle has the highest number of shared DEGs across sex, with hypothalamus having the least shared DEGs. Among the upregulated DEGs, we observed that gWAT, iWAT, BAT, and liver have the highest number of DEGs that are female-specific, whereas skeletal muscle and hypothalamus have the highest number of DEGs that are male-specific. Among the downregulated DEGs, we noted that gWAT, BAT, liver, and skeletal muscle have the highest number of DEGs that are female-specific, whereas iWAT and hypothalamus have the highest number of DEGs that are male-specific.

**Figure 8.**
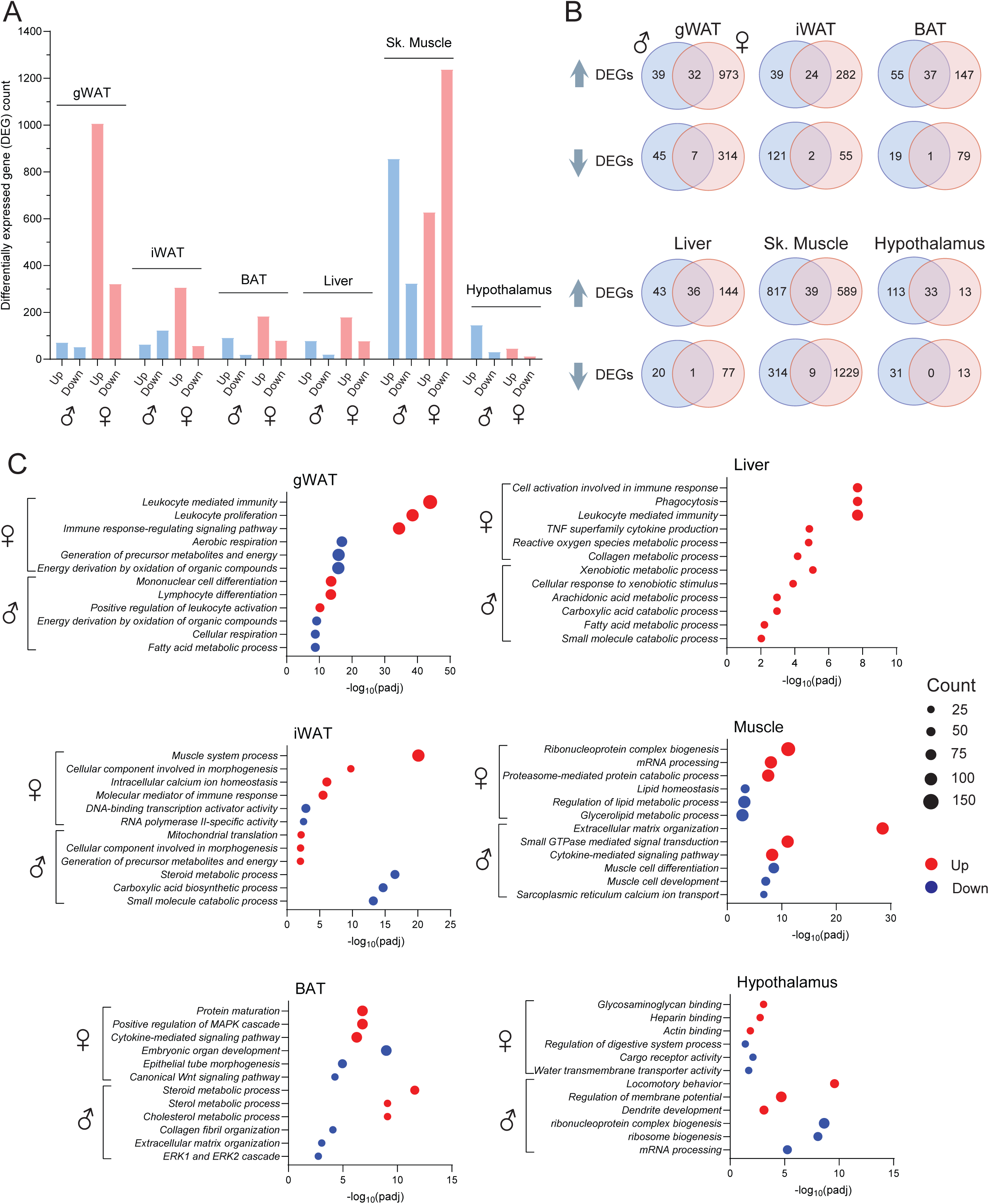
Altered pan-tissue transcriptomes and biological pathways in Ts66Yah mice. **(A)** Number of differentially expressed genes (DEGs) that are up or down regulated across six tissues in chow-fed male and female Ts66 mice. A DEG is defined as any gene with log2(FC) > 0.5 and padj (FDR) < 0.05. *N* = 6 per genotype per tissue. gWAT, gonadal white adipose tissue; iWAT, inguinal white adipose tissue; BAT, brown adipose tissue. **(B)** Overlap analysis showing DEGs that are shared between males and females, as well as those DEGs found in males or females only, across six tissues. **C)** Gene ontology highlighting some of the top biological pathways altered across six tissues in male and female Ts66 mice.

We performed gene ontology (GO) analysis to reveal which major biological pathways are enriched in Ts66Yah mice. Among the top biological pathways upregulated in female mice are those related to immune response and cytokine signaling (gWAT, iWAT, BAT, and liver), and the top downregulated pathways are those related to energy metabolism, cellular respiration, and lipid metabolism (gWAT and skeletal muscle) (**Fig. 8C**). In male mice, some of the top biological pathways upregulated include those related to immune response (gWAT), mitochondrial translation (iWAT), steroid and cholesterol metabolism (BAT), xenobiotic and arachidonic acid metabolism (liver), cytokine signaling (skeletal muscle), and extracellular matrix remodeling (skeletal muscle), and the down-regulated pathways include those related to cellular respiration, fatty acid metabolism, steroid, carboxylic acid, and small molecule catabolic processes (gWAT, iWAT, liver), mRNA processing (hypothalamus), MAPK signaling (BAT), and muscle cell function (skeletal muscle) (**Fig. 8C**). We highlighted some of the DEGs involved in immune activation, cellular respiration, fatty acid, lipid, and steroid metabolism, cytokine signaling, ECM remodeling, and regulation of reactive oxygen species in gWAT, iWAT, BAT, liver, and/or skeletal muscle (**Figure 8 – figure supplement 1-5**). These transcriptomic changes are broadly consistent with our physiological data and suggest that these concerted molecular changes across tissues likely contribute, at least in part, to the systemic metabolic dysfunction seen in Ts66Yah mice. Altogether, these data underscore major sex differences in the pan-tissue transcriptome, but also reveal common and shared biological pathways affected in Ts66Yah males and females that underpin their shared metabolic deficits.

### Metabolic response of Ts66Yah mice to an obesogenic diet

Individuals with DS are prone to developing obesity (10, 12). Given that diet is a major environmental factor contributing to obesity, we challenged the Ts66Yah mice with an obesogenic high-fat diet (HFD) to determine their ability to cope with metabolic stress associated with overnutrition. Weight gain over time in HFD-fed Ts66Yah males was largely indistinguishable from euploid controls, aligning with body composition where fat and lean mass were also not different between genotypes (**Fig. 9A,B**). Interestingly, while the trajectory of weight gain in Ts66Yah females was also indistinguishable from euploid controls, they gained greater lean mass on HFD (**Fig. 9C,D**). We performed indirect calorimetry analysis after the male and female mice were on HFD for 13 and 14 weeks, respectively. We observed no differences in food intake, physical activity, and energy expenditure across the circadian cycles and metabolic states (fed, fasted, and refed) in either Ts66Yah male or female mice (**Fig. 9E,F**). ANCOVA analyses (using lean mass as a covariate of energy expenditure) also indicated no differences in energy expenditure between genotypes of either sex across the circadian cycles and metabolic states (**Figure 9 – figure supplement 1**).

**Figure 9.**
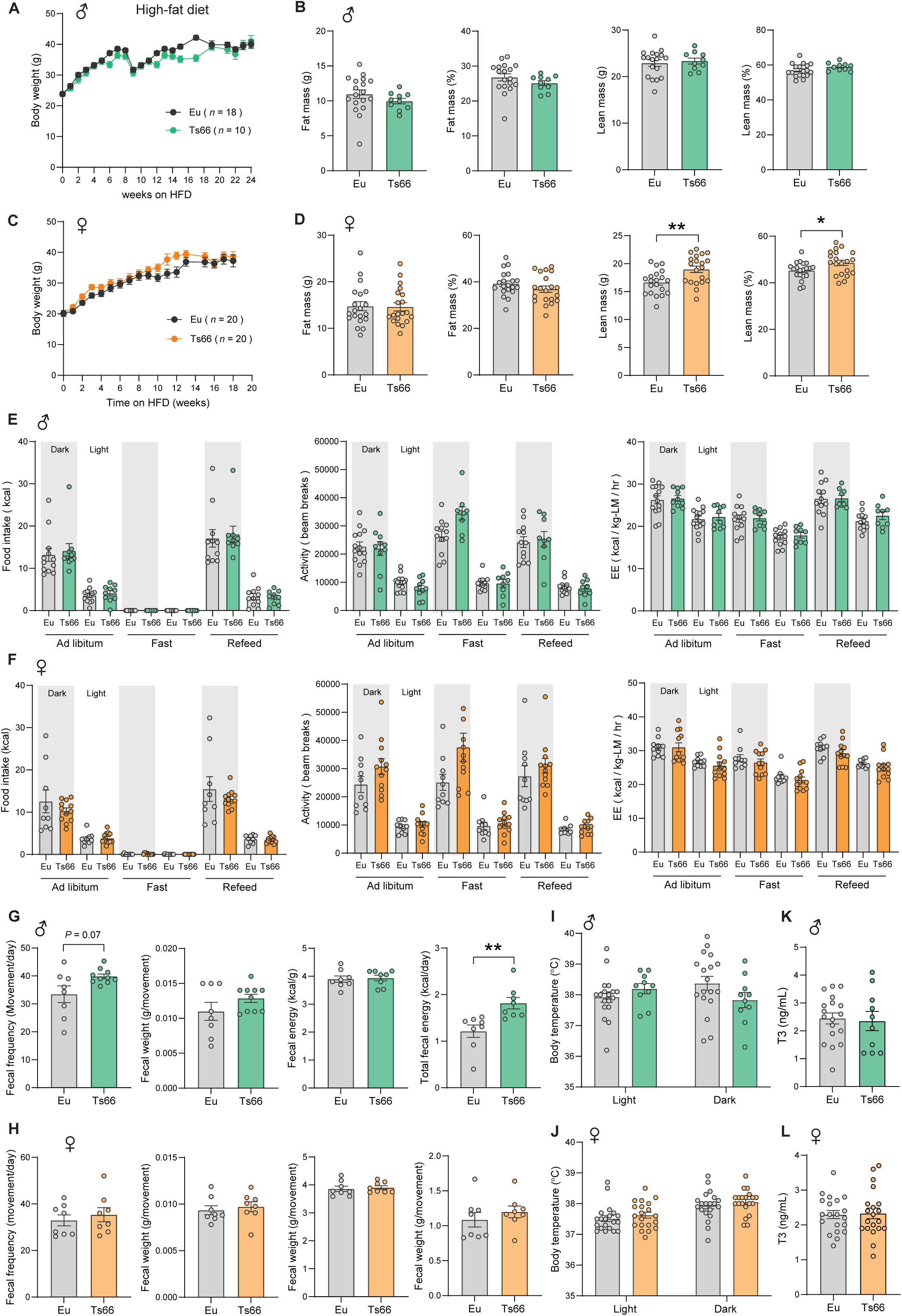
Impact of high-fat diet on metabolic parameters in Ts66Yah mice. **(A)** Body weight of HFD-fed male Ts66 mice and euploid (Eu) controls over time. **(B)** Absolute and relative (% of body weight) fat and lean mass in male mice (Eu = 18; Ts66 = 10). **(C)** Body weight of HFD-fed female Ts66 mice and euploid controls over time. **(D)** Absolute and relative (% of body weight) fat and lean mass in female mice (Eu = 20; Ts66 = 20). **(E-F)** Food intake, total physical activity level, and energy expenditure of male (E) and female (F) euploid and Ts66 mice across the circadian cycle (light and dark) and metabolic states (*ad libitum* fed, fast, refeed). Sample size for male (Eu = 9-14; Ts66 = 9-10) and female (Eu = 9-10; Ts66 = 12) mice. **(G-H)** Fecal frequency, average fecal weight, and fecal energy content (per gram and total) in male (G) and female (H) euploid and Ts66 mice. Sample size for male (Eu = 8; Ts66 = 10) and female (Eu = 8; Ts66 = 8) mice. **(I-J)** Body temperature in the light and dark cycle of male (I) and female (J) euploid and Ts66 mice. Sample size for male (Eu = 18; Ts66 = 10) and female (Eu = 20; Ts66 = 20) mice. **(K-L**) Serum triiodothyronine (T3) levels in male (K) and female (L) euploid and Ts66 mice. Sample size for male (Eu = 18; Ts66 = 10) and female (Eu = 20; Ts66 = 20) mice. All data are presented as mean ± SEM. * *P*<0.05; ** *P*<0.01. For body weight over time, data were analyzed by 2- way ANOVA with Sidek post hoc tests; other data were analyzed by two-tailed Student’s *t*-Test.

Since Ts66Yah mice fed standard chow have sex differences in fecal output, we again quantified fecal output and frequency, as well as fecal energy content in mice fed an HFD. Although none of the fecal parameters was different in Ts66Yah females, we observed an increase in fecal frequency (i.e., number of bowel movements per day) that contributed to higher total fecal energy content (**Fig. 9G-H**). This female-specific HFD effect stands in contrast to the male-specific fecal phenotype observed under chow, further underscoring how sex and diet interact to shape gastrointestinal function in Ts66Yah mice. Because we observed differences in the body temperature of chow-fed mice, we also measured body temperature across the circadian cycle, as well as serum T3 and sex hormone levels, in mice fed an HFD. None of these parameters were different between genotypes of either sex (**Fig. 9I-L and Fig 9 - figure supplement 2**).

At the termination of study, we assessed whether there were differences in tissue weight in Ts66Yah mice on HFD. At the time of tissue collection, male and female mice were 23 and 24 weeks old, and on HFD for a duration of 17 and 18 weeks, respectively. For Ts66Yah males, body weights and the absolute tissue weights of gWAT, iWAT, liver, heart, and kidney were not different from euploid controls; however, the relative (normalized to body mass) weights of liver and kidney were higher (**Figure 9 – figure supplement 3**). For Ts66Yah females, the absolute and relative weights of liver, and the absolute weights of heart, were higher; and the relative weights of iWAT were lower (**Figure 9 – figure supplement 3**). Taken together, these data indicate sex differences in the physiological response of Ts66Yah mice to an obesogenic diet, and highlighted the differential impact of HFD on organ size in these animals.

### High-fat diet exacerbates glucose intolerance and insulin resistance in Ts66Yah female mice

On standard chow, Ts66Yah mice developed pronounced glucose intolerance, insulin resistance, and dyslipidemia. We next asked if HFD would further exacerbate these phenotypes. In overnight (16 h) fasted Ts66Yah males, blood glucose, serum insulin, triglyceride, and cholesterol levels were not different from euploid controls, whereas NEFA and β-hydroxybutyrate (ketone) levels were significantly higher (**Fig. 10A**), suggesting greater adipose lipolysis and hepatic fat oxidation and ketogenesis in the fasted state. In the refed state, however, serum insulin, glucose, and lipid profile of Ts66Yah males were indistinguishable from euploid controls. In Ts66Yah females, serum parameters in the fasted and refed states were largely not different from euploid controls, except higher refed insulin levels (**Fig. 10B**). A similar blood glucose level but with elevated insulin in the refed state suggests diminished insulin action in Ts66Yah female mice.

**Figure 10.**
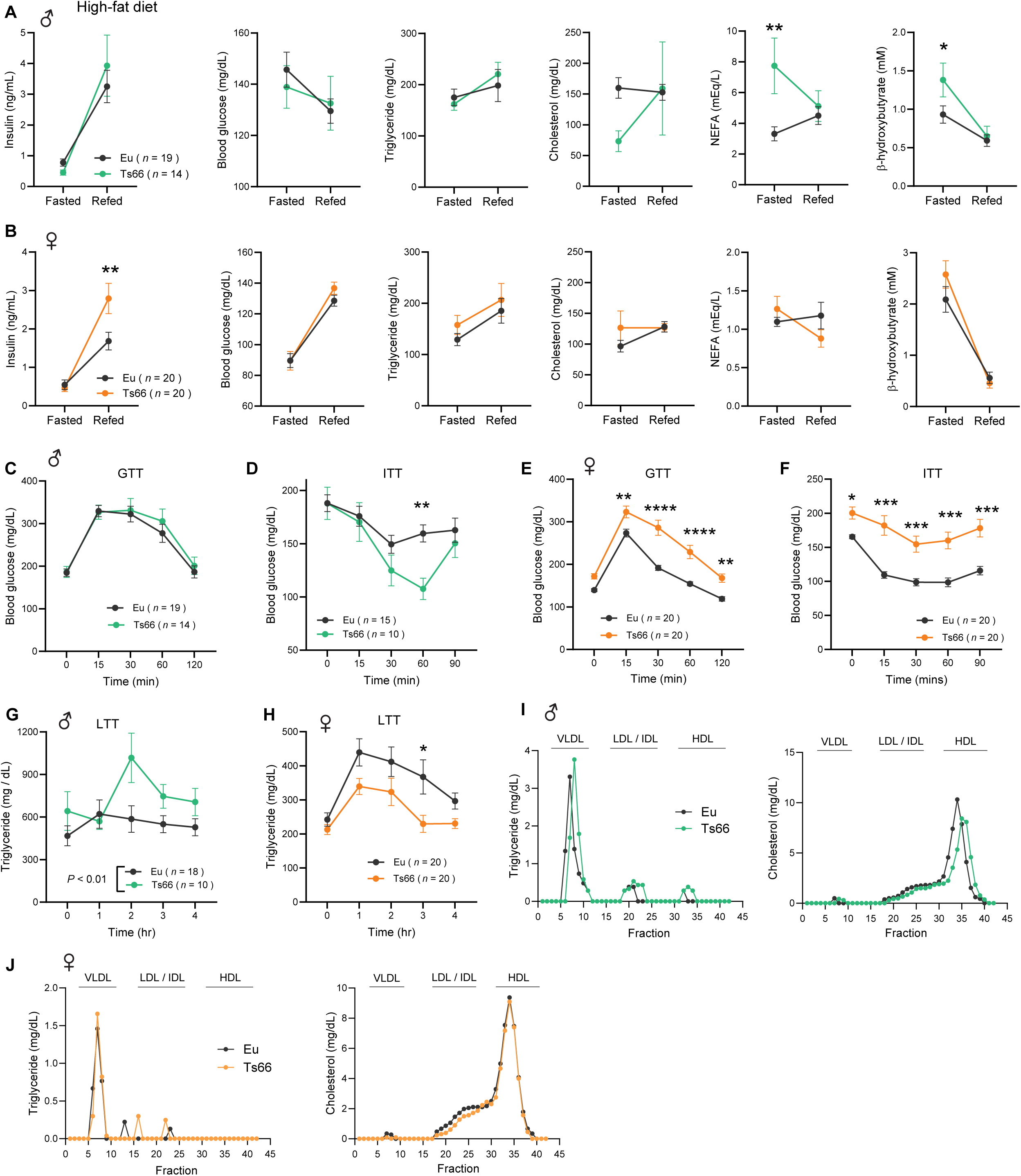
High-fat diet exacerbates glucose intolerance and insulin resistance in Ts66Yah mice. **(A-B)** Overnight fasting insulin, blood glucose, serum triglyceride, cholesterol, non-esterified free fatty acids (NEFA), and β-hydroxybutyrate (ketone) in male (A) and female (B) Ts66 mice and euploid (Eu) controls. Sample size for male mice (Eu = 19; Ts66 = 14) and female mice (Eu = 20; Ts66 = 20). **(C-F)** Glucose intolerance as determined by the glucose tolerance test (GTT) in male (C) and female (E) euploid and Ts66 mice. Exacerbated insulin resistance as determined by the insulin tolerance test (ITT) in male (D) and female (F) euploid and Ts66 mice. Sample size for male mice (WT = 15-19; Ts66 = 10-14) and female mice (Eu = 20; Ts66 = 20). **(G-H)** The rate of triglyceride clearance in response to lipid gavage as determined by the lipid tolerance test (LTT) in male (G) and female (H) euploid and Ts66 mice. Sample size for male mice (Eu = 18; Ts66 = 10) and female mice (Eu = 20; Ts66 = 20). **(I-J)** Pooled mouse sera from male (I) and female (J) euploid and Ts66 mice were fractionated by fast protein liquid chromatography (FPLC), and the triglyceride and cholesterol content of each fraction was quantified. Fractions corresponding to very-low density lipoprotein (VLDL), low-density lipoprotein (LDL), intermediate-density lipoprotein (IDL), and high-density lipoprotein (HDL) are indicated. All data are presented as mean ± SEM. * *P*<0.05; ** *P*<0.01; *** *P*<0.001. For all tolerance tests, data were analyzed by 2-way ANOVA with Sidek post hoc tests.

Next, we subjected Ts66Yah mice on HFD to a glucose tolerance test. Ts66Yah females, but not males, showed exacerbated glucose intolerance compared to euploid controls (**Fig. 10C and E**). Direct assessment of insulin sensitivity showed that Ts66Yah females have markedly reduced glucose clearance in response to insulin injection compared to euploid controls, whereas Ts66Yah males exhibited greater insulin sensitivity relative to euploid controls (**Fig. 10D and F**).

We performed lipid tolerance tests to determine whether Ts66Yah mice on HFD have impaired lipid handling capacity. We noted sex differences in lipid clearance after an acute lipid load. Whereas Ts66Yah males had diminished capacity to promote triglyceride clearance, Ts66Yah females had faster rate of triglyceride clearance compared to euploid controls (**Fig. 10G-H**). Analysis of lipoprotein profiles showed that both Ts66Yah males and females have a modestly higher VLDL-TG than euploid controls (**Fig. 10I-J**). Unlike the females, Ts66Yah males had lower HDL-cholesterol. Taken together, these data indicate that while there are important sex differences, HFD worsens the insulin resistance and dyslipidemia phenotypes in Ts66Yah mice despite similar weight gain and adiposity.

### High-fat diet impairs mitochondrial respiratory capacity in Ts66Yah mice

We next assessed the impact of HFD on mitochondrial function. In Ts66Yah males, neither mitochondrial respiration through CI, CII, or CIV, nor ATP synthase (CV) activity were significantly different from euploid controls (**Fig. 11**). In Ts66Yah females, however, we observed reduced mitochondrial respiratory capacity in BAT, skeletal muscle, and liver, with the effect most pronounced in BAT (**Fig. 11**). Notably, this pattern represents a striking reversal from the chow-fed state, in which skeletal muscle mitochondrial respiratory capacity was elevated in Ts66Yah females relative to euploid controls (**Fig. 4**), underscoring the negative impact of an obesogenic diet on mitochondrial function. As with the males, mitochondrial ATP synthase activity was also not different between genotype in females. These data indicate that HFD impairs mitochondrial function in a tissue- and sex-dependent manner in Ts66Yah mice.

**Figure 11.**
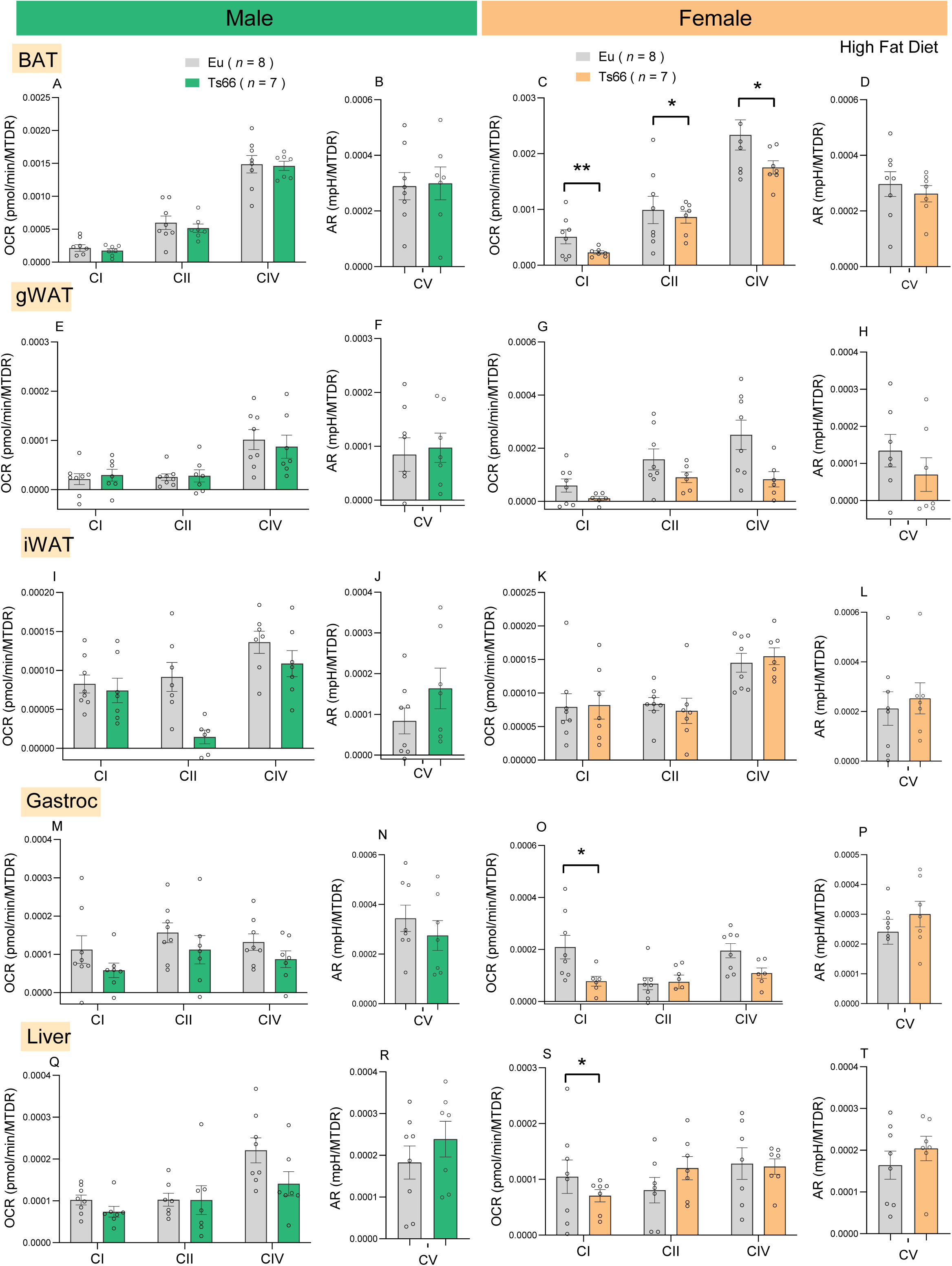
High-fat diet reduces mitochondrial respiratory capacity in Ts66Yah female mice. **(A-T)** Mitochondrial respiration through complex I (CI), CII, and CIV, as well as CV (ATP synthase) activity in BAT, gWAT, iWAT, gastroc, and liver of Tss66 male and female mice and euploid controls. All oxygen consumption rates (OCR) and Acidification Rates (AR) are normalized to mitochondrial content (based on MTDR). MTDR, MitoTracker Deep Red; BAT, brown adipose tissue; gWAT, gonadal white adipose tissue; iWAT, inguinal white adipose tissue; gastroc, gastrocnemius. Sample size for male (Eu = 8; Ts66 = 7) and female (Eu = 8; Ts66 = 7) mice. All data are presented as mean ± SEM. * *P*<0.05; ** *P*<0.01 (two-tailed Student’s *t*-Test)

### High-fat diet elevates hepatic injury, tissue fibrosis and oxidative stress in Ts66Yah mice

Hepatic injury, tissue fibrosis, and oxidative stress are frequent hallmarks of diet-induced obesity (73, 74). We therefore asked whether HFD worsens these conditions in Ts66Yah mice. In Ts66Yah males, but not females, we observed higher serum ALT levels, suggesting greater hepatic injury induced by an obesogenic diet (**Fig. 12A,D**). We noted that HFD increased the fibrotic marker (hydroxyproline) in gWAT, but not in iWAT and liver in Ts66Yah females; in contrast, Ts66Yah males had a lower level of the fibrotic marker in gWAT relative to euploid controls (**Fig. 12B,E**). While oxidative stress, as judged by malondialdehyde (MDA) level, was not different across tissues in Ts66Yah males, it was significantly higher in the gWAT and liver of Ts66Yah females relative to euploid controls (**Fig. 12C,F**). These data indicate that HFD affects hepatic injury, as well as tissue fibrosis and oxidative stress in a tissue- and sex-dependent manner in Ts66Yah mice.

**Figure 12.**
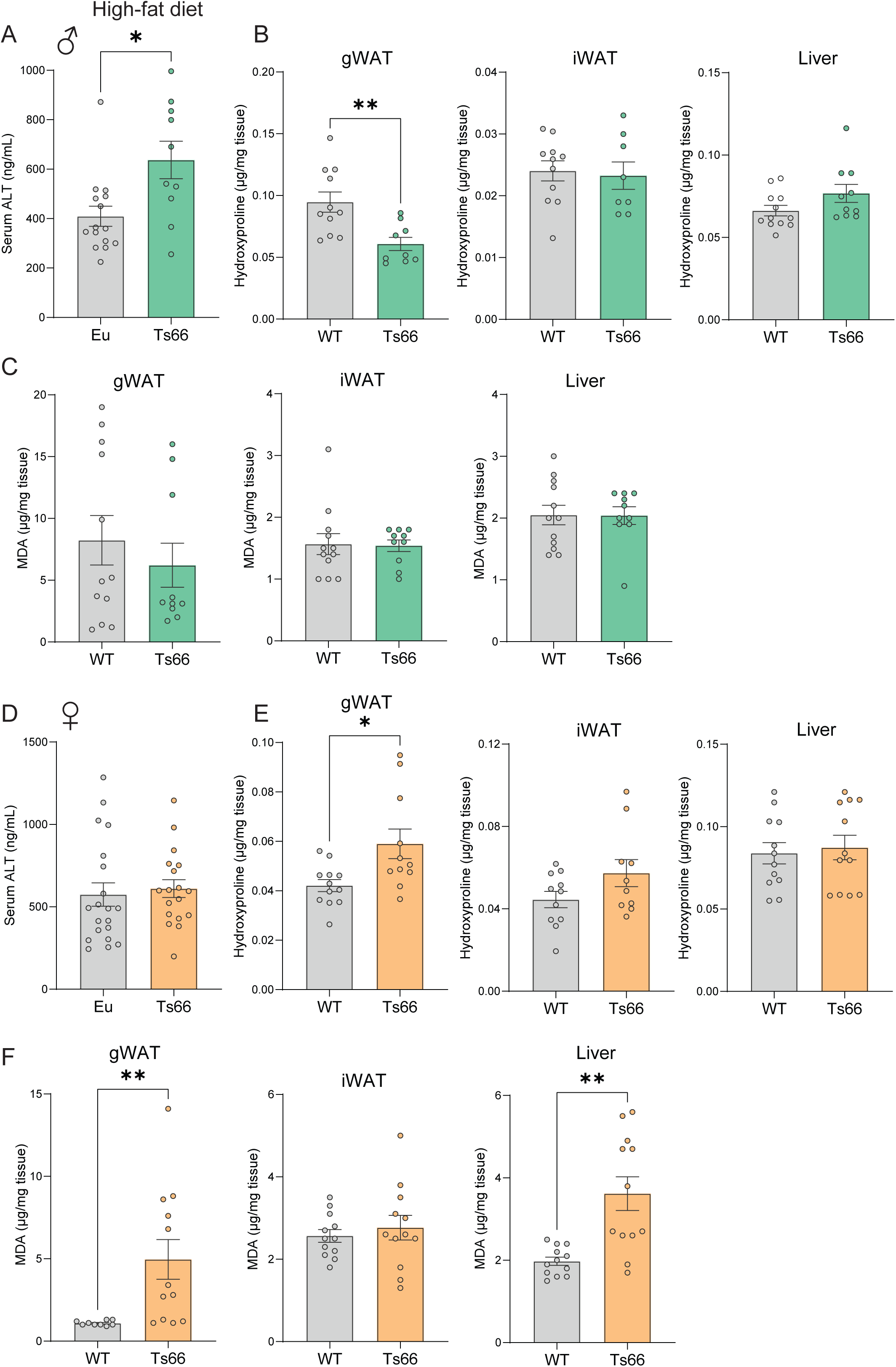
High-fat diet promotes liver injury and tissue fibrosis and oxidative stress in Ts66Yah mice. **(A-F)** Quantification of serum ALT (marker of liver injury), as well as hydroxyproline content (marker of fibrosis) and malondialdehyde (MDA; marker of oxidative stress) in gWAT, iWAT, and liver of male (A-C) and female (D-F) Ts66 mice and euploid (Eu) controls. gWAT, gonadal white adipose tissue; iWAT, inguinal white adipose tissue. Sample size: male Eu = 11-12 and Ts66 = 8-10; female Eu = 9-11 and Ts66 = 10-12. All data are presented as mean ± SEM. * *P*<0.05; ** *P*<0.01 (two-tailed Student’s *t*-Test)

### High-fat diet worsens hepatic steatosis in Ts66Yah mice

The higher serum ALT level in Ts66Yah male mice prompted us to examine liver histology for signs of injury and inflammation (**Fig. 13A**). Hepatic steatosis appeared to be significantly more pronounced in Ts66Yah males compared to euploid controls (Fig. 13B). We quantified markers of cell injury in histological sections; these include hepatocyte ballooning with or without Mallory-Denk bodies, apoptotic hepatocytes (acidophil body), confluent hepatocyte necrosis, and the presence of megamitochondria. We also quantified markers of inflammation; these include foci of lobular and portal inflammation, lipogranuloma, and the presence of giant cells and pigmented macrophages. Our quantifications revealed no significant differences in cell injury and inflammation between Ts66Yah males and euploid controls (**Fig. 13B**). In Ts66Yah females, none of the histological parameters was significantly different from euploid controls (**Fig. 13C**). Given a heightened proinflammatory serum profile seen in chow-fed Ts66Yah mice, we therefore also measured the circulating levels of IL-1β, IL-2, IL-4, IL-5, IL-6, IL-10, TNF-α, KC/GRO (CXCL1), and INF-γ. Relative to euploid controls, only the levels of KC/GRO and IL-10 were higher in Ts66Yah males and females, respectively (**Fig. 13 – figure supplement 1**). These data suggest that while HFD worsens liver steatosis in a sex-dependent manner, it did not further exacerbate hepatic inflammation and cell injury, nor markers of systemic inflammation.

**Figure 13.**
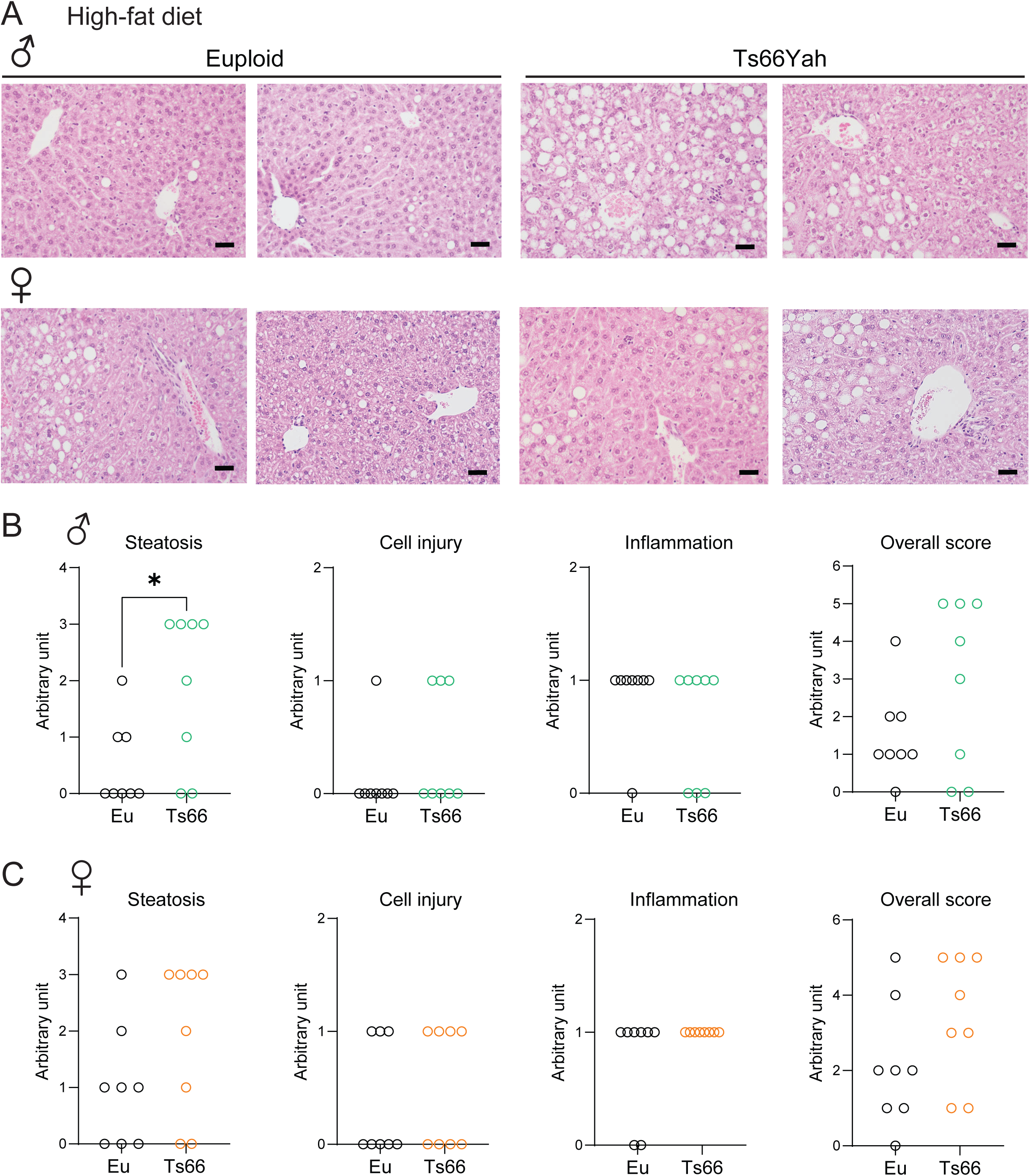
High-fat diet exacerbate liver steatosis in Ts66Yah mice. **(A)** two representative H&E- stained liver histology sections from male (top panel) and female (bottom panel) Ts66 mice and euploid (Eu) controls. Scale bar = 100 µM. **(B-C)** Histological assessment for steatosis, cell injury, inflammation, and overall score in male (B) and female (C) Ts66 mice and euploid controls. Sample size for male (Eu = 8; Ts66 = 8) and female (Eu = 8; Ts66 = 8) mice. * *P*<0.05 (two-tailed student *t*-Test)

### Phenotypic comparison between Ts66Yah and Ts65Dn mouse models

Ts66Yah model is derived from Ts65Dn, and represents an improved version whereby all the extra 46 triplicated protein-coding and non-coding genes unrelated to Hsa21 have been genetically removed (49), thus eliminating the potential confounding effects of the Mmu17 non-Hsa21 orthologous genes on the extra chromosome (58). Comparing the metabolic data of Ts66Yah with that of Ts65Dn (55) allows us to determine the effects and contributions of the triplicated non-Hsa21 gene orthologs to systemic metabolic health. In general, the behavioral, neurological, and bone phenotypes of Ts66Yah mice are milder than those of Ts65Dn mice (49, 58, 75–77). In contrast, we discovered that chow-fed Ts66Yah mice exhibit more severe metabolic phenotypes when compared to Ts65Dn mice (same genetic background), especially with regard to glucose intolerance and insulin resistance (**Table 3**). In the context of an obesogenic diet, however, we observed major sex differences. When challenged with an HFD, the Ts65Dn females exhibit clear glucose intolerance and pronounced insulin resistance whereas the Ts66Yah males have the opposite phenotype with improved insulin sensitivity relative to the euploid controls. In contrast, the HFD-fed Ts65Dn males exhibit glucose intolerance and insulin resistance whereas the Ts65Dn females appear indistinguishable from the euploid controls (55). Our data thus suggest that the removal of the confounding non-Hsa21 gene orthologs on the Mmu17 centromeric region reveals the more accurate impact of the triplicated Hsa21 gene orthologs on systemic metabolism under standard chow. It also reveals the confounding effects of the non-Hsa21 gene orthologs in dictating metabolic outcomes in the context of an obesogenic diet. Collectively, these findings highlight the differential impact of the triplicated Hsa21 gene orthologs on brain and bone versus peripheral metabolic tissues such as the adipose tissue, liver, and skeletal muscle.

**Table 3.**
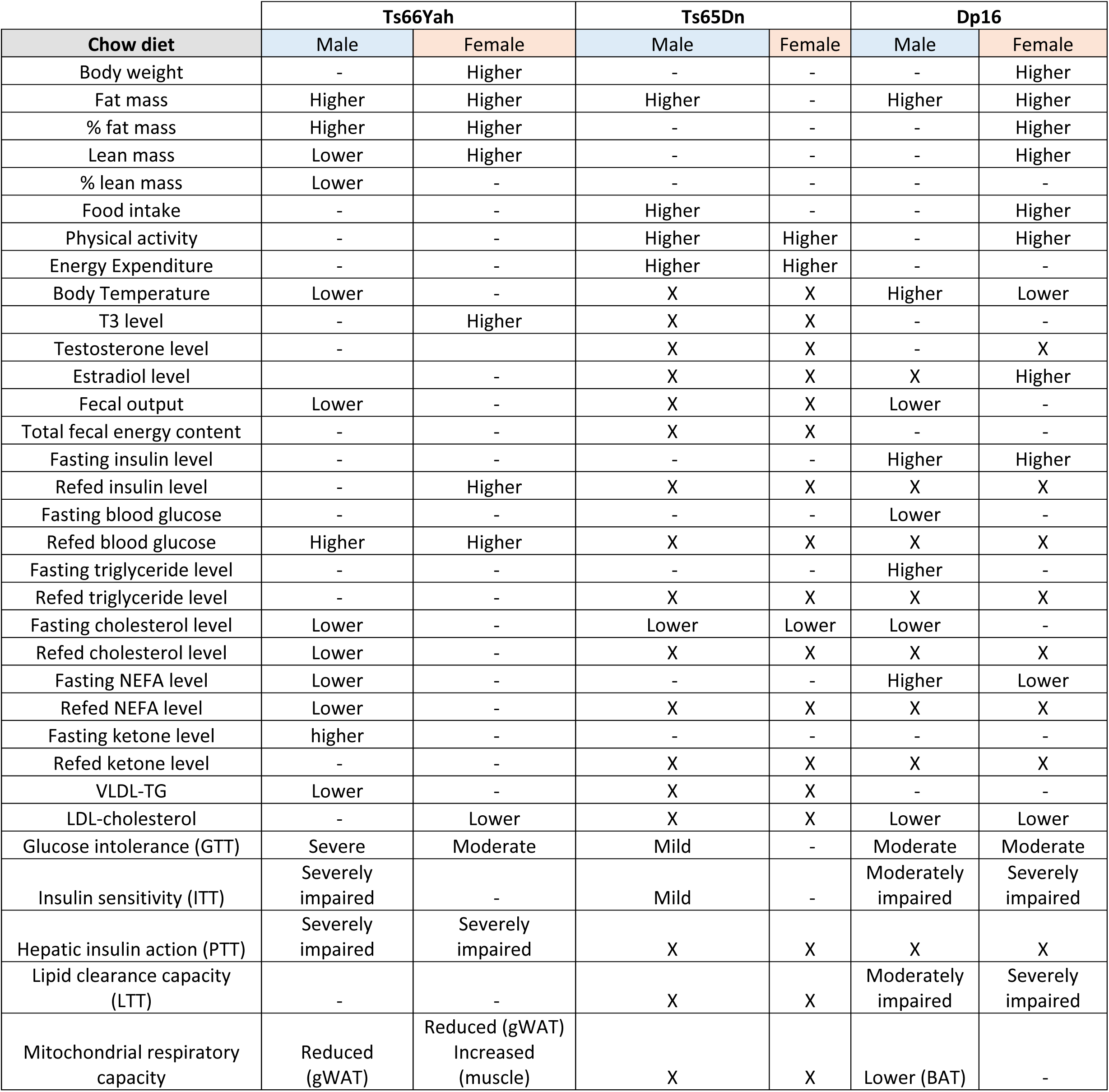
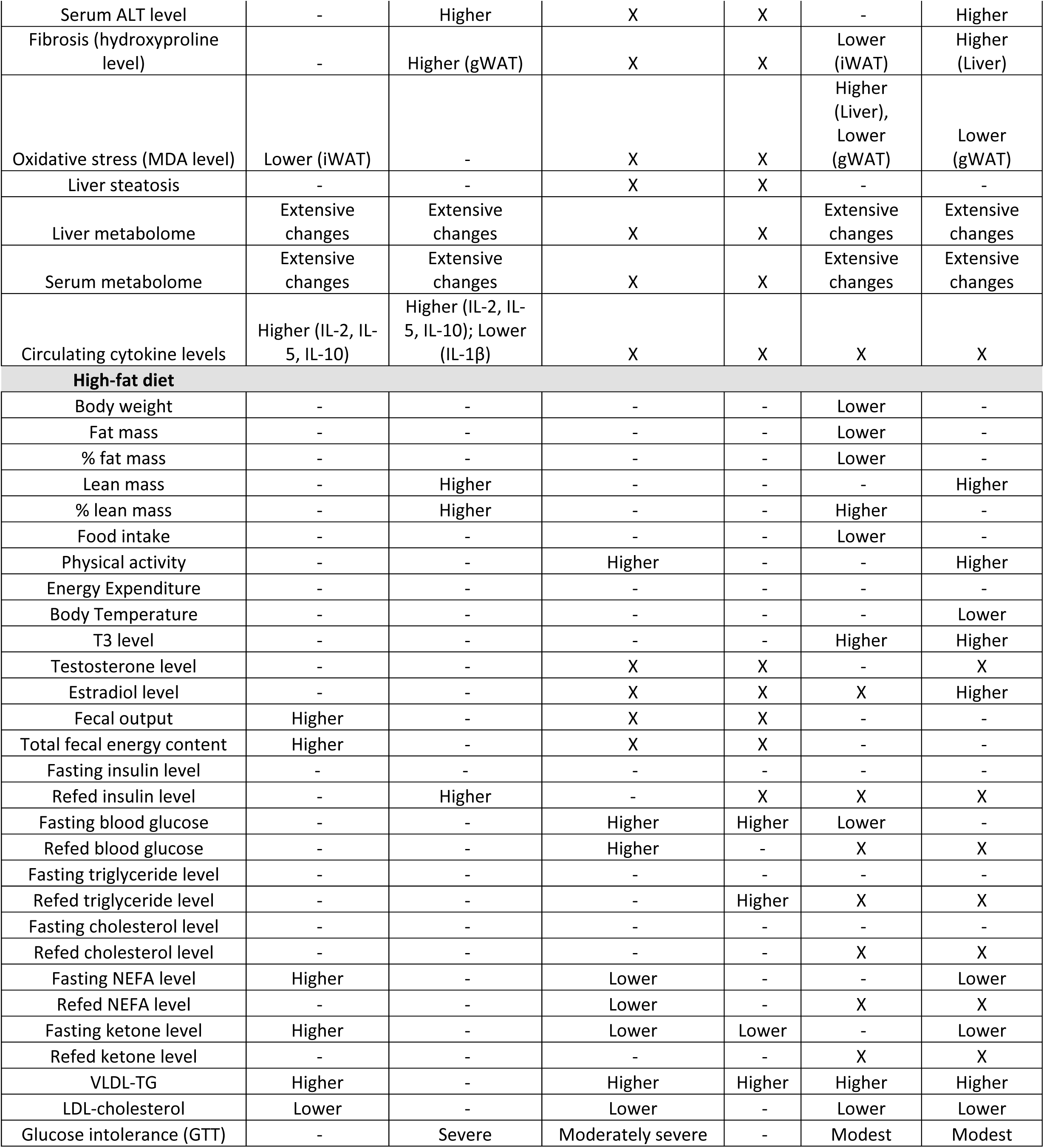
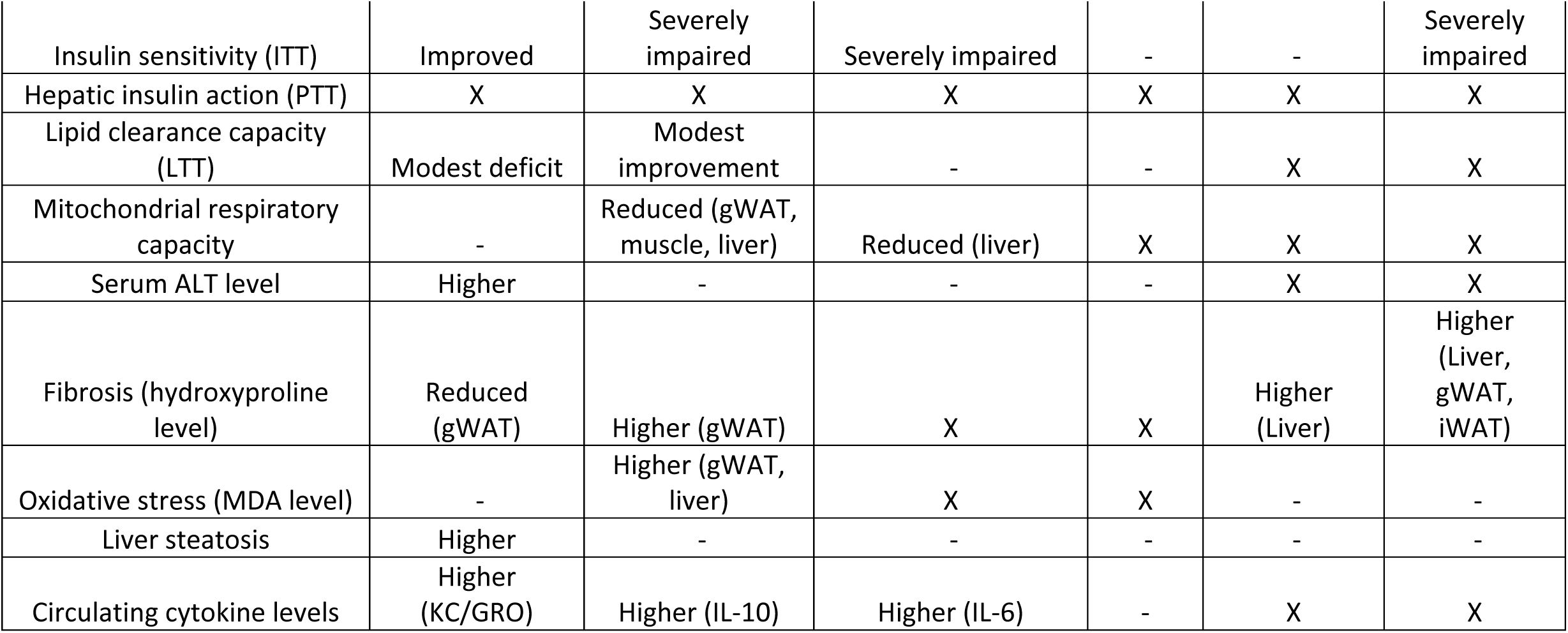
Comparison of the metabolic phenotypes of Ts66Yah mice with that of Ts65Dn (55) and Dp16 (60) mice. Phenotype that is not significantly different between genotypes is denoted with ‘-’. Assay or measurement that is not carried out is denoted with ‘X’. GTT, glucose tolerance test; ITT, insulin tolerance test; PTT, pyruvate tolerance test; LTT, lipid tolerance test; NEFA, non-esterified free fatty acid

### Phenotypic comparison between Ts66Yah and Dp16 mouse models

Recent studies have also suggested that the extra chromosome due to trisomy can contribute to phenotypic outcomes independent of the triplicated gene content (61), raising the hypothesis that some phenotypes are gene dosage-dependent whereas others are dependent on the presence of an extra chromosome. We therefore compared the metabolic phenotypes in Ts66Yah with that of the Dp16 mice (60). Dp16 mice carry a segmental duplication region of Mmu16 syntenic to Hsa21—generated by precise chromosomal engineering method—and carry no extra triplicated genes beyond the 115 triplicated Hsa21 gene orthologs (59). Ts66Yah and Dp16 mice share 103 identical triplicated genes (Mmu16: *Mir155*-*Zfp295*); thus, these two models have similar triplicated gene content, but only Ts66Yah mice carry an extra chromosome. There are a number of phenotypes shared between Ts66Yah and Dp16 mice. With respect to glucose intolerance and insulin resistance, Ts66Yah female mice exhibit a more severe phenotype when compared to Dp16 mice (**Table 3**). In contrast, histopathology scoring of the Dp16 mouse liver—where male and female data were combined—indicates elevated inflammation and fibrosis relative to wild-type contorls (67), which is absent in the Ts66Yah mice. We further compared the transcriptomic data from these two models. Surprisingly, only modest overlaps of up- or down-regulated DEGs are seen across gWAT, iWAT, BAT, liver, skeletal muscle, and hypothalamus in male and female mice (**Fig. 14**). In skeletal muscle and hypothalamus, there are more DEGs in Ts66Yah compared to Dp16 mice; in contrast, Dp16 mice have more DEGs in gWAT, iWAT, BAT, and liver than in Ts66Yah mice. Despite modest overlaps in DEGs across tissues, many of the affected biological pathways are similar between Ts66Yah and Dp16 mice; these include processes related to immune activation, glucose and lipid metabolism, mitochondrial function and cellular respiration, oxidative stress, and tissue fibrosis (60). We noted similar findings in our metabolomics data; similar classes of metabolites (e.g., bile acids, eicosanoids, oxylipins, lyso- phospholipids) are affected in both Ts66Yah and Dp16 mice (60). Thus, at the transcriptomic and metabolomic levels, a similar category of DEGs and metabolites—rather than identical gene sets and metabolites—appears to underpin the shared metabolic phenotypes between Ts66Yah and Dp16 mice. However, there are sufficient differences between Ts66Yah and Dp16 mice that suggest the possibility that trisomy may potentially affect phenotypic and transcriptomic outcomes independent of the triplicated gene content.

**Figure 14.**
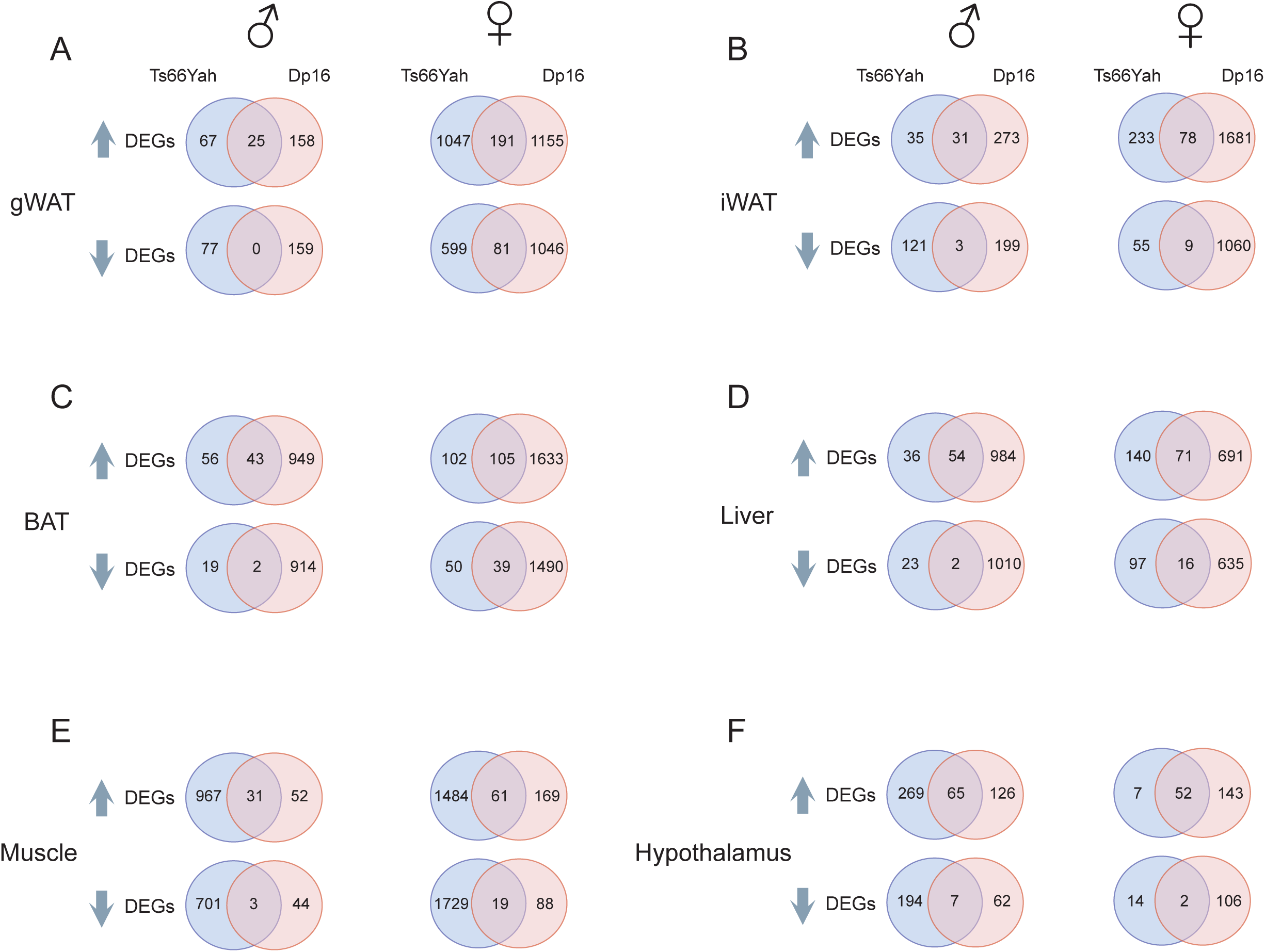
Overlap analyses of up- or down-regulated differentially expressed genes (DEGs) across tissues in Ts66Yah and Dp16 mice. DEGs up- or down-regulated in gWAT (A), iWAT (B), BAT (C), liver (D), skeletal muscle (E), and hypothalamus (F). DEG is defined as log2FC>0 and Padj (FDR) < 0.05.

## DISCUSSION

We set out to use the refined Ts66Yah trisomy model to directly address whether the reported metabolic phenotypes of Ts65Dn (55) are indeed attributed to the triplicated Hsa21 gene orthologs and not partially caused by the confounding effects of the extra triplicated non-Hsa21 gene orthologs located on the centromeric region of Mmu17 (49, 58, 75, 77). Based on comprehensive metabolic and multi-omics analyses, we showed that Ts66Yah mice exhibit extensive rewiring of pan-tissue transcriptomes, liver and serum metabolomes, and biological pathways that link to pronounced glucose intolerance, insulin resistance, and dyslipidemia. A heightened immune activation state, combined with impaired mitochondrial function and elevated tissue fibrosis and oxidative stress, likely contributed to the systems-wide metabolic dysregulation in Ts66Yah mice. The overall phenotype of Ts66Yah mice is broadly consistent with the susceptibility of individuals with DS to develop diabetes (9–12, 21), dyslipidemia (15, 17, 18, 21, 78), and reduced mitochondrial function (32). Some key phenotypes—glucose intolerance and insulin resistance— are more severe in the Ts66Yah than in the Ts65Dn mice. The collective findings validate Ts66Yah as a valuable genetic mouse model to understand DS-associated metabolic dysfunction. As the three remaining aneuploid models (Tc1, TcMAC21, and Ts65Dn) all have major limitations, our study helps establish Ts66Yah as a valuable trisomy model for comparative study to understand the contribution of gene dosage versus extra chromosome to DS phenotypes. Under the basal state when mice are fed a standard chow, Ts66Yah mice exhibit the most pronounced glucose intolerance and insulin resistance phenotypes when compared to the Ts65Dn and Dp16 mice. The Ts66Yah is thus a valuable model for genetic dissection of dosage-sensitive genes underpinning diabetes susceptibility in DS, and for testing therapeutic strategies to improve metabolic outcomes in DS.

In Ts66Yah mice, most of the triplicated Hsa21 gene orthologs are expressed at the expected ∼1.5-fold or higher across six metabolic tissues; their expression, however, is regulated in a sex- and tissue-dependent manner. This expression profile is consistent with the known variegated overexpression of Hsa21 genes across tissues in DS (79). Biological sex affects not only the expression of the triplicated genes, but also the disomic genes in adipose tissues, liver, skeletal muscle, and hypothalamus. The differential cascading effects of the triplicated genes on the rest of the transcriptome likely contributed, at least in part, to sex differences in phenotypic outcomes. For example, the Ts66Yah females have significantly more DEGs in adipose tissue and skeletal muscle than males, and these are associated with divergent weight trajectory; the Ts66Yah females are consistently heavier than their euploid littermates whereas the weight trajectory of Ts66Yah males is largely indistinguishable from or lower than their euploid controls. The numerous sex differences reported in the present study underscore and reinforce the known sex-dependent mechanisms that modulate tissue transcriptomes (80) and whole-body energy metabolism (81, 82). It is perhaps not surprising that sex is a biological determinant of metabolic risk in DS, as documented by sex differences in the susceptibility of individuals with DS to developing obesity and its associated co-morbidities (12, 23, 83–86).

Despite numerous sex differences noted (e.g., body weight, body composition, fecal output, and body temperature), both male and female Ts66Yah mice share a core phenotype that includes pronounced glucose and pyruvate intolerance, insulin resistance, and dyslipidemia. These core phenotypes are independent of divergent weight gain and adiposity, indicating that impaired glucose and lipid metabolism is due to the triplicated genes rather than simply a consequence of increased fat mass. The insulin resistance phenotype and fasting lipid and lipoprotein profiles are generally consistent with the clinical and epidemiological findings in DS (9–18). Metabolic deficits in the Ts66Yah mice are associated with reduced mitochondrial respiratory capacity, elevated tissue fibrosis and oxidative stress, chronic low-grade inflammation, and major rewiring of biological pathways due to altered pan-tissue transcriptomes and metabolomes; all of these changes are known to disrupt local and systemic metabolism (73, 87, 88). In light of the complex genetic perturbations in DS, and the integrative nature of metabolism due to inter-organ crosstalk (89), the metabolic phenotypes we observed in Ts66Yah mice are most likely due to the combined and concerted changes in transcriptome, metabolome, and biological pathways across major metabolic tissues rather than due to changes in one or two dominant pathways. Future studies are needed to provide greater insights into the molecular and cellular mechanisms underpinning metabolic dysfunction in Ts66Yah mice.

Similar to the Dp16 mice (60), our metabolomic analyses also uncovered extensive remodeling of the liver and serum metabolomes in Ts66Yah mice. The majority of differential metabolites are not shared between the sexes, a finding we have also seen in the Dp16 mice. This once again underscores the complex interactions of triplicated genes and biological sex in determining phenotypic outcome. The differential metabolites are clustered around pathways related to lipid, amino acid, and bile acid metabolism; several of these have been previously documented in DS (67, 70). Despite sex differences, several classes of metabolites with important signaling properties—bile acids, lyso-phospholipids, oxylipins, and eicosanoids—are shared by Ts66Yah males and females, a finding that is similar to the Dp16 mice. Aside from aiding digestion, primary and conjugated bile acids made in the liver, and secondary bile acids produced by gut bacteria, also play diverse signaling roles in modulating local and systemic metabolism (69, 90, 91). Altered bile acid pools in Ts66Yah mice—both primary (e.g., glycocholic acid, taurocholic acid, taurohyocholate) and secondary bile acids (e.g., lithocholic acid, taurodeoxycholic acid, 7- ketodeoxycholic acid, glycolithocholic acid, taurohyodeoxycholic acid, ursocholyltaurine, murideoxycholic acid, sulfoglycolithocholate, hyodeoxycholic acid)—suggest potential changes in the gut-liver axis and bile acid–regulated signaling that may contribute to metabolic dysfunction. The complex changes in multiple immuno-regulatory eicosanoids (72) and oxylipins (92, 93) appear to promote a pro-inflammatory state in Ts66Yah mice; the heightened immune activation phenotype is further corroborated by our pan-tissue transcriptomic and multiplex cytokine profiling data. Interestingly, we noted that multiple phospholipid species (e.g., Lyo-PC, Lyso-PA, Lyso-PA, Lyso-PE) are either elevated or reduced in the Ts66Yah16 mice. Some of these phospholipids are known to have complex signaling roles (71); their altered levels may potentially link to some aspects of metabolic deficits in Ts66Yah mice. It is likely that the remodeling of tissue and serum metabolomes, in combination with major changes in pan-tissue transcriptomes, contributed to the adverse systemic metabolic outcome in Ts66Yah mice.

Diet is an important environmental factor that can modify the metabolic phenotype in the context of trisomy. How a calorie-dense diet high in fat content impacts metabolic parameters in DS becomes relevant as individuals with DS are increasingly exposed to varied lifestyles and diets due to longer lifespan. When challenged with a high-fat diet, the trajectories of weight gain in Ts66Yah males and females were largely indistinguishable from the euploid controls. Despite no differences in body weight and adiposity, we observed striking sex differences in how the Ts66Yah males and females handle glucose and lipid load. When administered a bolus of glucose, the Ts66Yah females clearly showed impaired glucose clearance whereas the males showed no deficit relative to euploid controls. Direct assessment of insulin action showed that the Ts66Yah females are profoundly insulin resistant whereas the Ts66Yah males exhibit the opposite phenotype with improved insulin sensitivity. Lastly, when administered a lipid load, the Ts66Yah males showed impaired triglyceride clearance whereas the Ts66Yah females exhibited the opposite phenotype with greater rate of lipid clearance. In chow-fed animals, the core metabolic phenotypes (i.e., glucose intolerance, insulin resistance, and dyslipidemia) are shared by the Ts66Yah males and females. In contrast, we noted a sexually dimorphic response to HFD in the Ts66Yah mice, where sex clearly influences the metabolic outcome in the context of trisomy. These data strongly argue for the inclusion of both sexes in pre-clinical DS models to capture the complex interactions between triplicated genes, biological sex, and environmental factors such as diet.

Which among the triplicated genes are causal drivers of the metabolic phenotypes? The Ts66Yah mice carry 103 triplicated Hsa21 gene orthologs, several of which are known to affect metabolism, inflammation, tissue fibrosis and oxidative stress. These include *Dyrk1a* (94), *Dscr1*/*Rcan1* (95, 96), *AtpJ* (97), *Atp5o* (98), *Tiam1* (99, 100), *Prdm15* (101), *Ripk4* (102), *Znf295* (103), *Hmgn1* (104), *Cbr1* (105), *Bach1* (106), *Fam3b* (107), *Ets2* (108), *Adamts5* (109, 110), *Usp16* (111), *Runx1* (112), *Sim2* (113), and the interferon receptor gene locus (*Ifnar2*, *Il10rb*, *Ifnar1*, *Ifngr2*) (114, 115). Some of these triplicated genes show sexually dimorphic expression in gWAT (*Runx1*, *Cbr1*, *Hmgn1*, *Atp5o*, *Adamts5*), iWAT (*Il10rb*, *Adamts5*, *Ripk4*, *Bach1*, *Dyrk1a, Cbr1*, *Hmgn1*, *Atp5o*), BAT (*Adamts5*, *Tiam1*, *Ripk4*, *Bach1*, *Prdm15*), liver (*Atp5o*, *Ets2*, *Prdm15*), skeletal muscle (*Il10rb*, *Cbr1*, *Tiam1*, *Ifnar2*, *Atp5o*, *Sim2*, *Prdm15*, *Ifnar1*), and hypothalamus (*Il10rb*, *Hmgn1*, *Tiam1*, *Atp5o*). It is possible that the sex-biased expression of the triplicated genes contributes directly or indirectly to the sex differences in metabolic phenotypes in Ts66Yah mice. In the Dp16 mice, normalizing the expression dosage of the interferon receptor gene locus back to that of wild-type control does not reverse the hepatic lipid metabolism profile, thus ruling out the causal role of the interferon receptor gene locus in dictating hepatic lipid phenotype (67). However, it is presently unknown whether normalizing the expression dosage of the other triplicated genes—individually or in combination—could mitigate or reverse the metabolic outcomes in Ts66Yah mice. In recent years, several laboratories have successfully used an unbiased genetic dissection approach to pinpoint dosage-sensitive genes causally linked to DS phenotypes (116–119). Although laborious and costly, a similar approach is needed to establish causal relationships between the triplicated gene(s) and metabolic deficits in the Ts66Yah mice. Given the complex genetics and metabolic phenotypes of the Ts66Yah mice, it is most likely that multiple dosage-sensitive genes—acting additively or synergistically—are involved in dysregulating metabolism across tissues and in different dietary contexts.

An intriguing recent finding using two newly generated compound mouse models [Ts65Dn;Df(17)2Yey/+ and Dp(16)1Yey/Df(16)8Yey]—identical in triplicated gene content (a total of 103) but differing only in whether the mice carry a freely segregating extra chromosome—has suggested that trisomy itself can contribute to some DS phenotypes at the behavioral (T-maze) and transcriptomic (cerebral cortex) levels independent of the gene dosage effects (61). The Ts66Yah mice are genetically similar to the Ts65Dn;Df(17)2Yey/+ mice; in both models, the triplicated non-Hsa21 gene orthologs have either been genetically deleted (as in Ts66Yah) or normalized to two copies (as in Ts65Dn;Df(17)2Yey/+). The Ts66Yah mice share an identical 103 triplicated genes with the Dp16 mice. There are 12 additional triplicated genes carried by the Dp16, but not Ts66Yah, mice. Thus, these two models have similar triplicated gene content but only the Ts66Yah mice carry a freely segregating extra chromosome. The differences in phenotypes and pan-tissue transcriptomes between Ts66Yah and Dp16 mice provide additional support for the hypothesis that trisomy and gene dosage imbalance contribute differentially to DS phenotypes. This notion is consistent with recent findings suggesting that aneuploidy induces a general stress response (120), as well as disrupting the nuclear architecture and 3D genome organization in trisomic cells (121–123).

We wish to highlight several limitations of the study. First, no single mouse model fully recapitulates the entire spectrum of DS phenotypes (124, 125). Ts66Yah mice carry only 103 (∼54%) of the Hsa21 gene orthologs. It is known that the triplicated Hsa21 gene orthologs interact in complex ways to affect DS phenotypes and tissue transcriptomes (5, 126). While the Ts66Yah model clearly recapitulates some aspects of the metabolic deficits in DS, it remains to be determined whether mouse models carrying the full complement of the Hsa21 gene orthologs (127) would have similar or more severe metabolic dysfunction. Secondly, it is known that changes in mRNA abundance do not always correspond to reciprocal changes in protein level (128, 129). It would be informative to carry out parallel proteomic studies to determine the degree of concordance or discordance between transcriptomic and proteomic data, thus helping to prioritize candidate genes/proteins for further functional studies. Thirdly, the Dp16 mice are maintained on a C57BL/6J genetic background whereas Ts66Yah mice are on a B6EiC3Sn.BLiAF1/J background. Despite differences in severity, some of the key metabolic phenotypes—glucose intolerance and insulin resistance—are shared by the Ts66Yah and Dp16 mice, and are thus independent of mouse strains. However, whether some of the phenotypic and transcriptomic differences between the Ts66Yah and Dp16 mice are due to strain-dependent differences remain to be determined. Although the triplicated gene content between the Ts66Yah and Dp16 mice are similar, they are not identical because the Dp16 mice carry an additional 12 triplicated genes. A better comparison of metabolic phenotypes would be the use of the two compound models where the triplicated gene content is identical (61). Fourthly, although we strive to be comprehensive, our metabolic assays are not exhaustive. Specifically, we did not assess potential changes in pancreatic β-cell number, density, and secretory capacity in the Ts66Yah mice even though reduced pancreatic insulin content and defective glucose-induced insulin secretion have been previously observed in the Dp16 mice (130). In the Ts66Yah mice, we observed that physiologic fasting and refeeding insulin levels were either not different from euploid controls or were significantly elevated, thus suggesting that β- cell secretory capacity remains largely intact. Nevertheless, we could not rule out potential cytoarchitecture changes in the islet of Langerhans and subtle functional changes in the glucagon-producing α-cell, insulin-producing β-cell, and somatostatin-producing δ-cell. Lastly, we did not consider the potential contribution of gut microbiota to modulating host metabolic response to different diets (131).

In summary, our comprehensive molecular, biochemical, and physiological data have provided compelling evidence to support the recently generated Ts66Yah as a valid and valuable trisomy mouse model to study DS-associated metabolic dysfunction. The wealth of data generated provides an important foundation and physiological contexts to further dissect the mechanisms underlying metabolic dysregulation in DS, as well as the differential contribution of gene dosage and aneuploidy to DS phenotypes. Our data also indicate clear differences in phenotypes between the refined Ts66Yah model and the widely used Ts65Dn model (55), thus highlighting the confounding effects of the triplicated non-Hsa21 gene orthologs on systemic metabolism. Our current study on Ts66Yah mice, along with our previous studies on TcMAC21 (51), Ts65Dn (55), and Dp16 (60) mice, provided valuable information to inform the selection of appropriate DS mouse models to study DS-related metabolic phenotypes.

## MATERIALS AND METHODS

### Mouse model

Ts(17^16^)66Yah/J (abbreviated Ts66Yah) mice and euploid (Eu) littermate controls, maintained on B6EiC3Sn.BLiAF1/J genetic background, were obtained from the Jackson Laboratory (Strain # 036600). Mice were fed a standard chow (Envigo; 2018SX) or a high-fat diet (HFD; 60% kcal derived from fat, #D12492, Research Diets, New Brunswick, NJ). Mice were housed in polyethylene terephthalate (PET) cages on a 12h:12h light-dark photocycle (lights on at 6 am, lights off at 6 pm) with *ad libitum* access to water and food. For the HFD-fed group, HFD was provided beginning at 6 weeks old and continued for 18 weeks. At termination of the study, all mice were fasted for 2 h and euthanized. The age of mice at the time of tissue harvest: male and female mice fed a standard chow were 20 and 21weeks old, respectively; male mice fed an HFD were 23 weeks old (on HFD for 17 weeks); female mice fed an HFD were 24 weeks old (on HFD for 24 weeks). Tissues were collected, snap-frozen in liquid nitrogen, and kept at 80°C until analysis.

All mouse protocols were approved by the Institutional Animal Care and Use Committee of the Johns Hopkins University School of Medicine (animal protocol # MO25M273). All animal experiments were conducted in accordance with the National Institute of Health guidelines and followed the standards established by the Animal Welfare Acts.

### Body composition analysis

Body composition analyses for total fat mass, lean mass, and water content were determined using a quantitative magnetic resonance instrument (Echo-MRI-100, Echo Medical Systems, Waco, TX) at the Mouse Phenotyping Core facility at Johns Hopkins University School of Medicine. The chow-fed male and female group were analyzed at 11 and 12 weeks of age, respectively. The HFD-fed male mice were analyzed at 18 weeks of age (12 weeks on HFD), and HFD-fed female mice at 19 weeks of age (13 weeks on HFD).

### Indirect calorimetry

Chow- or HFD-fed Ts66Yah male and female mice and WT littermates were used for simultaneous assessments of daily body weight change, food intake (corrected for spillage), physical activity, and whole-body metabolic profile in an open flow indirect calorimeter (Comprehensive Laboratory Animal Monitoring System, CLAMS; Columbus Instruments, Columbus, OH) as previously described (132). In brief, data were collected for three days to confirm mice were acclimatized to the calorimetry chambers (indicated by stable body weights, food intakes, and diurnal metabolic patterns), then data were analyzed for the subsequent three days. Mice were observed with *ad libitum* access to food, throughout the fasting process, and in response to refeeding. Rates of oxygen consumption (*V̇*_O2_; mL·kg^-1^·h^-1^) and carbon dioxide production (*V̇*_CO2_; mL·kg^-1·^h^-1^) in each chamber were measured every 24 min. Respiratory exchange ratio (RER = *V̇*_CO2_/*V̇*_O2_) was calculated by CLAMS software (version 5.18) to estimate relative oxidation of carbohydrates (RER = 1.0) versus fats (RER = 0.7), not accounting for protein oxidation. Energy expenditure (EE) was calculated as EE= *V̇*_O2_× [3.815 + (1.232 × RER)] and normalized to lean mass. We also performed ANCOVA analysis on EE using body weight as a covariate (63). Physical activity (total and ambulatory) was measured by infrared beam breaks in the metabolic chamber. Average metabolic values and summed intake and activity values were calculated per subject and averaged across subjects for statistical analysis by Student’s t-test. The chow-fed male and female group were analyzed at 12 and 13 weeks of age, respectively. The HFD-fed male mice were analyzed at 19 weeks of age (13 weeks on HFD), and HFD-fed female mice at 20 weeks of age (14 weeks on HFD).

### Body Temperature

Deep colonic temperature was measured by inserting a lubricated (Medline, water-soluble lubricating jelly, MDS032280) probe (Physitemp, BAT-12 Microprobe Thermometer) into the anus of mice at a depth of 2 cm. Stable numbers were recorded in both the dark and light cycle for each mouse. Deep colon temperature measurements on the chow-fed male and female group were performed at 17 weeks of age. The HFD-fed male mice were measured at 17 weeks of age (11 weeks on HFD), and HFD-fed female mice at 18 weeks of age (12 weeks on HFD).

### Fecal bomb calorimetry and assessment of fecal parameters

Fecal pellet frequency and average fecal pellet weight were monitored by housing each mouse singly in clean cages and counting the number of fecal pellets and recording their weight at the end of a 24 h period. Fecal pellets were shipped to the University of Michigan Animal Phenotyping Core for fecal bomb calorimetry. Briefly, fecal samples were dried overnight at 50°C prior to weighing and grinding them to powder. Each sample was mixed with wheat flour (90% wheat flour, 10% sample) and formed into 1.0 g pellet, which was then secured into the firing platform and surrounded by 100% oxygen. The bomb was lowered into a water reservoir and ignited to release heat into the surrounding water. Together these data were used to calculate fecal pellet frequency (bowel movements/day), average fecal pellet weight (g/bowel movement), fecal energy (cal/g feces), and total fecal energy (kcal/day). Fecal bomb calorimetry and assessment of fecal parameters for the chow-fed male and female group were performed at 17 weeks of age. The HFD-fed male mice were assessed at 17 weeks of age (11 weeks on HFD), and female HFD-fed mice at 18 weeks of age (12 weeks on HFD).

### Glucose, insulin, and lipid tolerance tests

All tolerance tests were conducted as previously described (133–135). For glucose tolerance tests (GTTs), mice were fasted for 6 h before glucose injection. Glucose (Sigma, St. Louis, MO) was reconstituted in saline (0.9 g NaCl/L) to a final concentration of 1 g/10 mL (for the chow-fed mice) or 2g/10 mL (for the HFD-fed mice), sterile-filtered, and injected intraperitoneally (i.p.) at 1 mg/g body weight (i.e., 10 μL/g body weight for chow-fed mice or 5 μL/g body weight for HFD-fed mice). Blood glucose was measured at 0, 15, 30, 60, and 120 min after glucose injection using a glucometer (Contour next, Parsippany, NJ). GTT for the male chow-fed group was performed at 11 weeks of age, female chow-fed group at 12 weeks of age, male HFD-fed group at 15 weeks of age (9 weeks on HFD), and female HFD-fed mice at 16 weeks of age (10 weeks on HFD). For insulin tolerance tests (ITTs), food was removed 2 h before insulin injection. 6.5 μL of insulin stock (4 mg/mL; Gibco) was diluted in 10 mL of saline, sterile-filtered, and injected i.p. at 0.75 U/kg body weight (i.e., 10 uL/g body weight) for chow-fed mice and 1 U/kg (i.e., 13.3 µL/g body weight) for high-fat diet (HFD) mice. Blood glucose was measured at 0, 15, 30, 60, and 90 min after insulin injection using a glucometer (Contour next). ITT for the male chow-fed group was performed at 11 weeks of age, female chow-fed group at 12 weeks of age, male HFD-fed group at 16 weeks of age (10 weeks on HFD), and female HFD-fed group at 17 weeks of age (11 weeks on HFD). For lipid tolerance tests (LTTs), mice were fasted for 12 h and then injected i.p. with 20% emulsified Intralipid (soybean oil; Sigma; 10 μL/g of body weight). Sera were collected via tail bleed using a Microvette® CB 300 (Sarstedt) at 0, 1, 2, 3, and 4 h post-injection. Serum triglyceride levels were quantified using kits from Infinity Triglycerides (Thermo Scientific). LTT for the male chow-fed group was performed at 19 weeks of age, female chow-fed group at 18 weeks of age, the male HFD-fed group at 21 weeks of age (15 weeks on HFD), and female HFD-fed mice at 22 weeks of age (16 weeks on HFD).

### Fasting and refeeding glucose, insulin, and lipid profile

Mice were fasted overnight (∼16 h), beginning at 1 h before the dark cycle (around 5 pm). Clean cages were provided before food withdrawal. Overnight fasting blood glucose levels from tail bleed were measured using a glucometer. Serum was collected at the 16 h fast (around 10 am in the morning), and at 2 h after food was reintroduced (refeeding). Fasting and refeeding insulin, triglyceride, cholesterol, non-esterified free fatty acids (NEFA), and β-hydroxybutyrate levels were quantified. Age of the mice at the time of assessment: male chow-fed group at 10 weeks of age, female chow-fed group at 11 weeks of age, male HFD-fed group at 13 weeks of age (7 weeks on HFD), and female HFD-fed group 14 weeks of age (8 weeks on HFD).

### Blood and tissue chemistry analysis

Tail vein blood samples were allowed to clot on ice and then centrifuged for 10 min at 10,000 x *g*. Serum samples were stored at -80°C until analyzed. Serum triglycerides (TG) and cholesterol were measured according to manufacturer’s instructions using an Infinity kit (Thermo Fisher Scientific, Middletown, VA). Non-esterified free fatty acids (NEFA) were measured using a Wako kit (Wako Chemicals, Richmond, VA). Serum β-hydroxybutyrate (ketone) concentrations were measured with a StanBio Liquicolor kit (StanBio Laboratory, Boerne, TX). Serum insulin (Crystal Chem, 90080), T3 (Calbiotech, T3043T-100), testosterone (Cayman, 582701), estradiol (Cayman, 501890), and alanine aminotransferase (ALT; Abcam, ab282882) levels were measured using commercial kits according to manufacturer’s instructions.

Hydroxyproline assay (Sigma Aldrich, MAK569) was used to quantify total collagen content in liver and adipose tissues (gWAT and iWAT) according to the manufacturer’s instructions, with specific optimizations for each tissue type. Tissues were homogenized in deionized water using a bead mill homogenizer. Liver, iWAT, and gWAT were homogenized to a concentration of 0.1mg tissue/µL. All samples were hydrolyzed with an equal volume of HCl at 120°C for 3 hours. The volume of hydrolyzed tissue samples plated for the dehydration step was optimized for each tissue type and experimental group to ensure all measurements fell within the linear range of the standard curve. Hydroxyproline content was calculated based on a standard curve and normalized to tissue weight.

Lipid peroxidation levels (marker of oxidative stress) in liver and adipose tissues (gWAT and iWAT) were assessed by the quantification of malondialdehyde (MDA) levels via the Thiobarbituric Acid Reactive Substances (TBARS) assay (Cayman Chemical, 700870) according to the manufacturer’s instructions.

### Multiplex profiling of serum proinflammatory cytokine level

Multiplex profiling of serum IL-1β, IL-2, IL-4, IL-5, IL-6, IL-10, TNF-α, KC/GRO (also known as CXCL1), and IFN-γ levels were performed on MESO QuickPlex SQ120 instrument (Mesoscale Discovery, Rockville, Maryland) using the S-PLEX mouse proinflammatory panel 1 kit (Mesoscale Discovery; cat # K15744S) according to manufacturer’s instruction. Data analyses were performed on Mesoscale Discovery (MSD) workbench.

### Serum lipoprotein-triglyceride and cholesterol analysis by FPLC

Food was removed for ∼2 h (in the light cycle) prior to blood collection. Sera collected from mice were pooled (*n* = 10-14/group) and sent to the Mouse Metabolism Core at Baylor College of Medicine for fast protein liquid chromatography (FPLC) separation. A total of 45 fractions were collected, and TG and cholesterol in each fraction were quantified.

### Extraction of hepatic lipids

Lipid extraction from frozen liver samples was performed using a modified Folch method (136). Briefly, approximately 25 mg of frozen liver tissue was weighed and homogenized in 400 µL of cold sucrose buffer (250 mM sucrose, 10 mM Tris, 1 mM EDTA) using a bead beater (FastPrep-24, MP Biomedical). The samples underwent three rounds of 20-second bead beating cycles. Lipids were then extracted from the liver homogenate by adding 1.5 mL of a chloroform:methanol (2:1, v/v) solution to the mixture. The sample was vortexed thoroughly and centrifuged at 1700 rpm for 5 minutes at 4℃ to separate the phases. The lower chloroform phase, containing lipids, was carefully transferred to a new tube. Each tube was then dried to dehydrate samples. The dried lipid extracts were reconstituted in 50 µL of tert-butanol:methanol:TritonX- 100 solution (3:1:1, v/v/v) for triglyceride and cholesterol quantification. Reconstituted samples were either used immediately or stored at -80°C until analysis.

### Hepatic cholesterol quantification

Following tissue extraction, total cholesterol content was quantified using the Infinity Cholesterol Reagent kit (Thermo Fisher Scientific, Middletown, VA) according to the manufacturer’s instructions. Quantified cholesterol values were normalized to tissue weight.

### Untargeted serum and liver metabolomic analyses

Serum and liver metabolites were extracted and subjected to LC-MS/MS detection on a TripleTOF 6600 mass spectrometer system at Novogene (Sacramento, CA) (*n* = 6 mice per genotype per sex). Age of the mice when serum samples and liver tissues were collected: male chow-fed group at 20 weeks of age, female chow-fed group at 21 weeks of age, male HFD-fed group at 23 weeks of age (17 weeks on HFD), and female HFD-fed group at 24 weeks of age (18 weeks on HFD). Data were processed using Novogene in-house analysis pipeline. In brief, the raw mass spectrometry data were first converted to mzXML format using ProteoWizard (137). Peak extraction, alignment, and retention time correction were then performed with XCMS software (138). The peaks with missing rate >50% in each group of samples were filtered. The blank values were filled with KNN and 1/5 minimum (1/5 minimum filled for blank values >50%, KNN filled for blank values <50%), and the peak area was corrected by SVR method. The metabolites were annotated by searching the Novogene’s in-house database, integrated public database, prediction database and metDNA. Finally, substances with a comprehensive identification score above 0.5 and a CV value of QC samples less than 0.3 were extracted, and then positive and negative mode were combined (substances with the highest qualitative grade and the lowest CV value were retained) to obtain the all_sample_data.xlsx file. A total of 1873 differential metabolite were identified from the 48 samples. Multivariate statistical analysis was conducted on the metabolites, including Principal Component Analysis (PCA) and Partial Least Squares Discriminant Analysis (PLS-DA). PLS-DA is a supervised discriminant analysis statistical method. This method uses partial least squares regression (139) to establish the relationship model between the relative quantitative value of metabolites and the sample category to realize the prediction of the sample category. The PLS-DA model of each comparison group was established, and the model evaluation parameters (R2, Q2) obtained by 7-cycle cross-validation. Differential metabolites were screened according to the criteria: VIP > 1.0 fold change (FC) > 1.2 or FC < 0.833 and *P*-value < 0.05 (Student’s t test were used when the data follow a normal distribution, otherwise Wilcoxon rank-sum test). VIP refers to the variable importance in the projection of the first principal component of the PLS-DA model, and the VIP value represents the contribution of the metabolites to the grouping. KEGG enrichment (FDR correction by Benjamini and Hochberg method) and GSEA analysis was performed on KEGG entries based on the changes in quantitative values of metabolites. All metabolomics data, raw spectral files, and details of experimental protocol and data analyses have been deposited in a public repository, the Metabolomics Workbench (140).

### Liver histology and pathology scoring

Scoring was based on murine system (141) adapted from the scoring system established by the Nonalcoholic Steatohepatitis Clinical Research Network working group (142). Formalin-fixed, paraffin embedded pieces of liver sectioned at 5 microns and stained with hematoxylin and eosin (H&E) were examined by a hepatobiliary pathologist (Robert A. Anders) using a Nikon Eclipse 50i compound light microscope with 4 and 10x objectives for scanning the entire surface area of the resection. A 20x objective was used for deeper resolution of abnormalities if needed. Steatosis scored as a percentage of the surface area of the liver tissue, not including portal tracts or large normal anatomic structures. Steatosis is defined as macrovesicular steatosis which includes both large and small droplet fat. Large lipid droplets are easily identified on H&E at scanning 4-10X magnification and is larger than an adjacent non-steatotic hepatocyte. Small droplet fat is easily visible at 10X as discrete lipid vacuoles in the hepatocyte cytoplasm. The percentage of steatosis was binned into 3rds. 0-5% = none (score of 0), 5-33% = mild (score of 1), 34-66% = moderate (score of 2), and 67 to 100% = severe (score of 3). Hepatocyte injury was scored as absent (score of 0) or present (score of 1). The types of injury include hepatocyte ballooning with or without Mallory-Denk bodies, apoptotic hepatocytes/acidophil body, confluent hepatocyte necrosis, megamitochondria. Inflammation was score as follow: 0 = absent, scored 1 = few/scattered with three or fewer separate foci, scored 2 = frequent with four or more separate foci. The overall score is the sum of all the score for steatosis, cell injury, and inflammation.

### Mitochondrial respirometry

Respirometry was conducted on frozen tissue samples to assay for mitochondrial activity as described previously (143, 144). Briefly, brown adipose tissue (BAT), gonadal white adipose tissue (gWAT), inguinal white adipose tissue (iWAT), gastrocnemius, (Gastroc) and liver were dissected from mice fed standard chow and HFD, snapped frozen in liquid nitrogen, and stored at -80°C for later analysis. Samples were thawed in MAS buffer (70mM sucrose, 220 mM mannitol, 5 mM KH_2_PO_4_, 5 mM MgCl_2_, 1 mM EGTA, 2 mM HEPES pH 7.4), finely minced with scissors, and then homogenized with a glass Dounce homogenizer. The resulting homogenate was spun at 1000 *g* for 10 min at 4°C. The supernatant was collected and immediately used for protein quantification by BCA assay (Thermo Scientific, 23225). Each well of the Seahorse microplate was loaded with 4 µg BAT, 15 µg iWAT 15 µg gWAT, 8 µg gastroc, and 8 µg liver homogenate protein. Each biological replicate consists of three technical replicates. Samples from all tissues were treated separately with NADH (1 mM) as a complex I substrate or Succinate (a complex II substrate, 5 mM) in the presence of rotenone (a complex I inhibitor, 2 µM), then with the inhibitors rotenone (2 µM) and antimycin A (4 µM), followed by TMPD (0.45 mM) and Ascorbate (1 mM) to activate complex IV, and finally treated with azide (40 mM) to assess non-mitochondrial respiration. Additionally, to isolate CV (ATP synthase)-dependent activity, three baseline time points were first recorded. All samples were then treated with 4 μM rotenone (Complex I inhibitor) and 4 μM antimycin A (Complex III inhibitor), followed by three additional measurement time points. Complex V was subsequently activated by the addition of 20 mM ATP (Sigma A26209) together with 7.87 mM carbonyl cyanide p-trifluoromethoxyphenylhydrazone (FCCP; Enzo BML-CM120), an uncoupler of oxidative phosphorylation, and three more time points were collected. Finally, 10 μM oligomycin (Sigma 495455) was added to inhibit Complex V in order to distinguish non-mitochondrial acidification. All mitochondrial respiration data were normalized to mitochondrial content, quantified using MitoTracker Deep Red (MTDR, ThermoFisher, M22426) as described (143, 144). Briefly, lysates were incubated with MTDR (1 µM) for 10 min at 37°C, then centrifuged at 2000 *g* for 5 min at 4°C. The supernatant was carefully removed and replaced with 1x MAS solution and fluorescence was read with excitation and emission wavelengths of 625 and 670 nm, respectively. To minimize non-specific background signal contribution, control wells were loaded with MTDR and 1x MAS and subtracted from all sample values.

### RNA-sequencing and bioinformatics analysis

Bulk RNA sequencing of Ts66Yah (*n* = 6) and euploid littermate control (*n* = 6) mouse liver, gWAT, iWAT, brown adipose tissue (BAT), skeletal muscle (gastrocnemius), and hypothalamus were performed by Novogene (Sacramento, California, USA) on a NovaSeq X Plus platform and pair-end reads (2 x 150 bp) were generated, with 6 G raw data per sample. Tissues were collected in male and female chow-fed group at 20 and 21 weeks of age, respectively. Sequencing data was analyzed using the standard Novogene Analysis Pipeline. Sequencing reads were aligned to Mus musculus reference genome (GRCm39/mm39). Data analysis was performed using a combination of programs, including Fastp, Hisat2, and FeatureCounts.

Differential expressions were determined through DESeq2. The resulting *P*-values were adjusted using the Benjamini and Hochberg’s approach for controlling the false discovery rate (FDR). Genes with an adjusted *P*-value (FDR) < 0.05 and log2(FC) > 0.5 were assigned as differentially expressed. Gene ontology (GO), Kyoto Encyclopedia of Genes and Genomes (KEGG), and Reactome (http://www.reactome.org) enrichment were implemented by ClusterProfiler. All volcano plots and heat maps were generated in Graphpad Prism 10 software. All statistics were performed on log transformed data. All heat maps were generated from column z-score transformed data. The z-score of each column was determined by taking the column average, subtracting each sample’s individual expression value by said average then dividing that difference by the column standard deviation. Z-score = (value – column average)/column standard deviation. High-throughput sequencing data from this study have been submitted to the NCBI Sequence Read Archive (SRA) under accession number # PRJNA1456105.

## Statistical analyses

All results are expressed as mean ± standard error of the mean (SEM). Statistical analysis and outlier tests were performed with GraphPad Prism 10 software (GraphPad Software, San Diego, CA). Data were analyzed with two-tailed Student’s *t*-tests or by repeated measures ANOVA. For two-way ANOVA, we performed Sidek or Bonferroni post hoc tests. *P* < 0.05 was considered statistically significant.

## AUTHOR CONTRIBUTIONS

GWW: Conceptualization; MS, FC, DCS, SA, GWW: Formal analysis; GWW: Funding acquisition; MS, FC, DKM, LT, NW, SA, GWW: Investigation; DCS, SA: Methodology; GWW: Project administration; GWW: Supervision; MS, FC, DCS, GWW: Visualization; GWW: Roles/Writing - original draft; ; MS, FC, DKM, LT, NW, SA: Writing - review & editing.

## ACKNOWLEDGEMENTS

The work was funded, in part, by grants from the National Institute of Health (DK084171 to GWW). MS and DCS were supported by an NIH T32 training grant (HL007534). The FPLC/serum analyses were conducted by the Mouse metabolic Phenotyping Center (MMPC) at Baylor College of medicine, funded by NIH grants DK114356 and UM1HG006348. The fecal bomb calorimetry analysis was performed at the University of Michigan Animal Phenotyping Core, supported by center grants 1U2CDK135066-01 (Mi-MPMOD) and DK020572 (MDRC). The Metabolomics Workbench/National Metabolomics Data Repository (NMDR) is supported by the NIH (U2C-DK119886), Common Fund Data Ecosystem (CFDE) (3OT2OD030544) and Metabolomics Consortium Coordinating Center (M3C) (1U2C-DK119889).

Research reported in this publication was supported by the National Institutes of Health. The content is solely the responsibility of the authors and does not necessarily represent the official views of the National Institutes of Health

## DATA AVAILABILITY

All RNA-seq data have been deposited in NCBI Sequence Read Archive (SRA), with the BioProject # PRJNA1456105. The metabolomics data is available at the NIH Common Fund’s National Metabolomics Data Repository (NMDR) website, the Metabolomics Workbench (https://www.metabolomicsworkbench.org) where it has been assigned Study ID (ST005076, ST005082).

The data can be accessed directly via it’s Project DOI: https://doi.org/http://dx.doi.org/10.21228/M8HV9K

## CONCLICT OF INTEREST

We declared that none of the authors has conflict of interest.

## SUPPLEMENTAL FIGURE FILES LEGENDS AND SOURCE DATA

**Figure 1 – figure supplement 1 source data 1.** Relative expression (fold change) and adjusted p-value (FDR) for all 41 protein-coding genes on the Mmu17 centromeric region across six tissues in male and female Ts66Yah mice.

**Figure 7 – Source data 1.** Differential metabolites in Ts66Yah male liver vs euploid controls. Differential metabolites criteria: VIP > 1.0, fold change (FC) > 1.2 or FC < 0.833 and *P*-value < 0.05. Sample name notation: male euploid (wild-type) liver (M_WT_L), male euploid serum (M_WT_S), male Ts66 liver (M_Ts66_L), male Ts66 serum (M_Ts66_S), female euploid liver (F_WT_L), female euploid serum (F_WT_L), female Ts66 liver (F_Ts66_L), female Ts66 serum (F_Ts66_S).

**Figure 7 – Source data 2.** Differential metabolites in Ts66Yah female mouse liver vs euploid controls. Differential metabolites criteria: VIP > 1.0, fold change (FC) > 1.2 or FC < 0.833 and *P*-value < 0.05. Sample name notation: male euploid (wild-type) liver (M_WT_L), male euploid serum (M_WT_S), male Ts66 liver (M_Ts66_L), male Ts66 serum (M_Ts66_S), female euploid liver (F_WT_L), female euploid serum (F_WT_L), female Ts66 liver (F_Ts66_L), female Ts66 serum (F_Ts66_S).

**Figure 7 – Source data 3.** Differential metabolites in Ta66Yah male mouse serum vs euploid controls. Differential metabolites criteria: VIP > 1.0, fold change (FC) > 1.2 or FC < 0.833 and *P*-value < 0.05. Sample name notation: male euploid (wild-type) liver (M_WT_L), male euploid serum (M_WT_S), male Ts66 liver (M_Ts66_L), male Ts66 serum (M_Ts66_S), female euploid liver (F_WT_L), female euploid serum (F_WT_L), female Ts66 liver (F_Ts66_L), female Ts66 serum (F_Ts66_S).

**Figure 7 – Source data 4.** Differentially expressed metabolites in Ts66Yah female mouse serum vs euploid controls. Differential metabolites criteria: VIP > 1.0, fold change (FC) > 1.2 or FC < 0.833 and *P*-value < 0.05. Sample name notation: male euploid (wild-type) liver (M_WT_L), male euploid serum (M_WT_S), male Ts66 liver (M_Ts66_L), male Ts66 serum (M_Ts66_S), female euploid liver (F_WT_L), female euploid serum (F_WT_L), female Ts66 liver (F_Ts66_L), female Ts66 serum (F_Ts66_S).

**Figure 8 – Source data 1.** Differentially expressed genes (DEGs) upregulated in the gonadal white adipose tissue (gWAT) of chow-fed Ts66 male mice relative to euploid controls.

**Figure 8 – Source data 2.** Differentially expressed genes (DEGs) down-regulated in the gonadal white adipose tissue (gWAT) of chow-fed Ts66 male mice relative to euploid controls.

**Figure 8 – Source data 3.** Differentially expressed genes (DEGs) upregulated in the inguinal white adipose tissue (iWAT) of chow-fed Ts66 male mice relative to euploid controls.

**Figure 8 – Source data 4.** Differentially expressed genes (DEGs) down-regulated in the inguinal white adipose tissue (iWAT) of chow-fed Ts66 male mice relative to euploid controls.

**Figure 8 – Source data 5.** Differentially expressed genes (DEGs) upregulated in the brown adipose tissue (BAT) of chow-fed Ts66 male mice relative to euploid controls.

**Figure 8 – Source data 6.** Differentially expressed genes (DEGs) down-regulated in the brown adipose tissue (BAT) of chow-fed Ts66 male mice relative to euploid controls.

**Figure 8 – Source data 7.** Differentially expressed genes (DEGs) upregulated in the liver of chow-fed Ts66 male mice relative to euploid controls.

**Figure 8 – Source data 8.** Differentially expressed genes (DEGs) down-regulated in the liver of chow-fed Ts66 male mice relative to euploid controls.

**Figure 8 – Source data 9.** Differentially expressed genes (DEGs) upregulated in the skeletal muscle (gastrocnemius) of chow-fed Ts66 male mice relative to euploid controls.

**Figure 8 – Source data 10.** Differentially expressed genes (DEGs) down-regulated in the skeletal muscle (gastrocnemius) of chow-fed Ts66 male mice relative to euploid controls.

**Figure 8 – Source data 11.** Differentially expressed genes (DEGs) upregulated in the hypothalamus of chow-fed Ts66 male mice relative to euploid controls.

**Figure 8 – Source data 12.** Differentially expressed genes (DEGs) down-regulated in the hypothalamus of chow-fed Ts66 male mice relative to euploid controls.

**Figure 8 – Source data 13.** Differentially expressed genes (DEGs) upregulated in the gonadal white adipose tissue (gWAT) of chow-fed Ts66 male mice relative to euploid controls.

**Figure 8 – Source data 14.** Differentially expressed genes (DEGs) down-regulated in the gonadal white adipose tissue (gWAT) of chow-fed Ts66 male mice relative to euploid controls.

**Figure 8 – Source data 15.** Differentially expressed genes (DEGs) upregulated in the inguinal white adipose tissue (iWAT) of chow-fed Ts66 female mice relative to euploid controls.

**Figure 8 – Source data 16.** Differentially expressed genes (DEGs) down-regulated in the inguinal white adipose tissue (iWAT) of chow-fed Ts66 female mice relative to euploid controls.

**Figure 8 – Source data 17.** Differentially expressed genes (DEGs) upregulated in the brown adipose tissue (BAT) of chow-fed Ts66 female mice relative to euploid controls.

**Figure 8 – Source data 18.** Differentially expressed genes (DEGs) down-regulated in the brown adipose tissue (BAT) of chow-fed Ts66 female mice relative to euploid controls.

**Figure 8 – Source data 19.** Differentially expressed genes (DEGs) upregulated in the liver of chow-fed Ts66 female mice relative to euploid controls.

**Figure 8 – Source data 20.** Differentially expressed genes (DEGs) down-regulated in the liver of chow-fed Ts66 female mice relative to euploid controls.

**Figure 8 – Source data 21.** Differentially expressed genes (DEGs) upregulated in the skeletal muscle (gastrocnemius) of chow-fed Ts66 female mice relative to euploid controls.

**Figure 8 – Source data 22.** Differentially expressed genes (DEGs) down-regulated in the skeletal muscle (gastrocnemius) of chow-fed Ts66 female mice relative to euploid controls.

**Figure 8 – Source data 23.** Differentially expressed genes (DEGs) upregulated in the hypothalamus of chow-fed Ts66 female mice relative to euploid controls.

**Figure 8 – Source data 24.** Differentially expressed genes (DEGs) down-regulated in the hypothalamus of chow-fed Ts66 female mice relative to euploid controls.

---

**Figure 1 – figure supplement 1.**
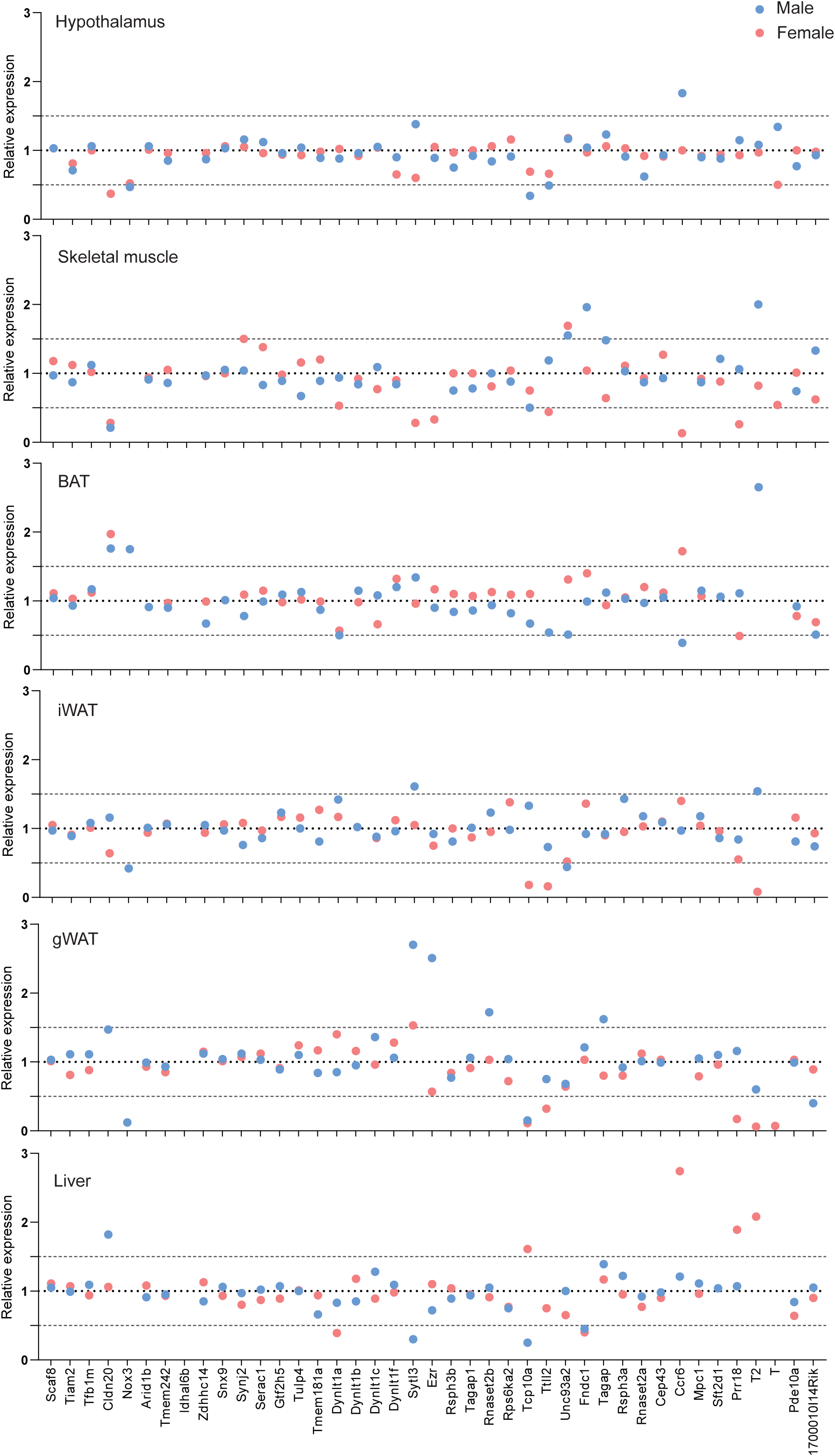
Normalization of expression dosage of triplicated genes on the centromeric region of Mmu17 in Ts66Yah mice. The relative expression level of the 41 protein-coding genes on the Mmu17 centromeric region was shown for liver, gonadal white adipose tissue (gWAT), inguinal white adipose tissue (iWAT), brown adipose tissue (BAT), skeletal muscle, and hypothalamus. Male data are indicated in blue and female data are indicated in pink.

**Figure 2 – figure supplement 1.**
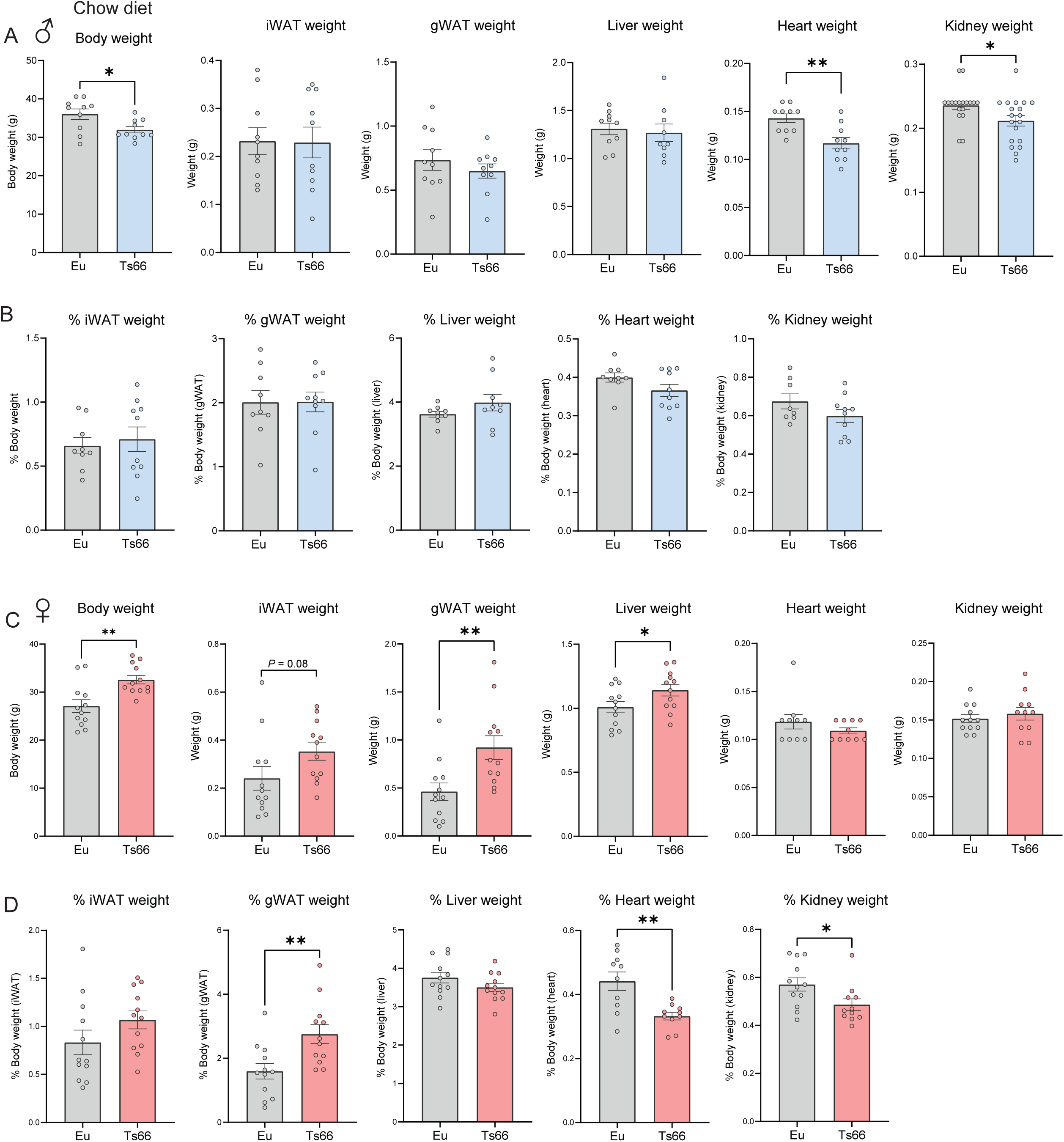
Body and tissue weights of chow-fed mice at termination of study. Tissues were collected from chow-fed male and female mice at 20 and 21 weeks of age, respectively. Body weights and the absolute (A and C) and relative (B and D; % of body weight) weights of iWAT, gWAT, liver, heart, and kidney in euploid (Eu) and Ts66 male (A-B) and female (C-D) mice. gWAT, gonadal white adipose tissue; iWAT, inguinal white adipose tissue. Sample size for male mice: Eu = 10; Ts66 = 10. Sample size for female mice: Eu = 12; Ts66 = 12. All data are presented as mean ± SEM. * *P*<0.05; ** *P*<0.01 (two-tailed Student’s *t*-TEST)

**Figure 2 – figure supplement 2.**
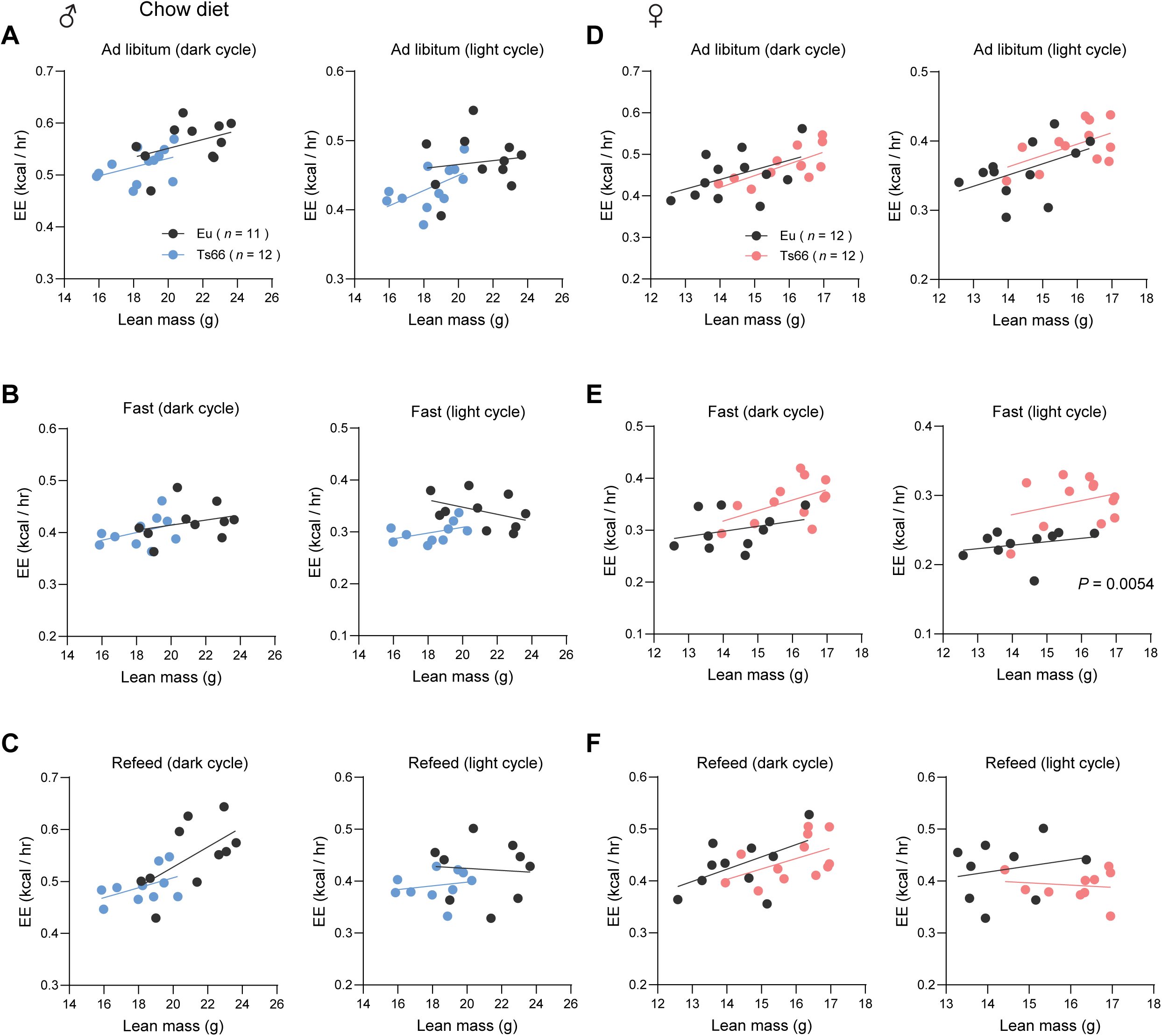
ANCOVA analysis of energy expenditure in chow-fed mice where lean mass is used as a covariate. ANCOVA analysis of male euploid (Eu) and Ts66 mice across the circadian cycle (dark and light) in *ad libitum* fed (A), fasted (B), and refed (C) states. ANCOVA analysis of female euploid and Ts66 mice across the circadian cycle (dark and light) in *ad libitum* fed (D), fasted (E), and refed (F) states. Male Sample size: Eu = 11; Ts66 = 12. Female sample size: Eu = 12; Ts66 = 12.

**Figure 2 – figure supplement 3.**
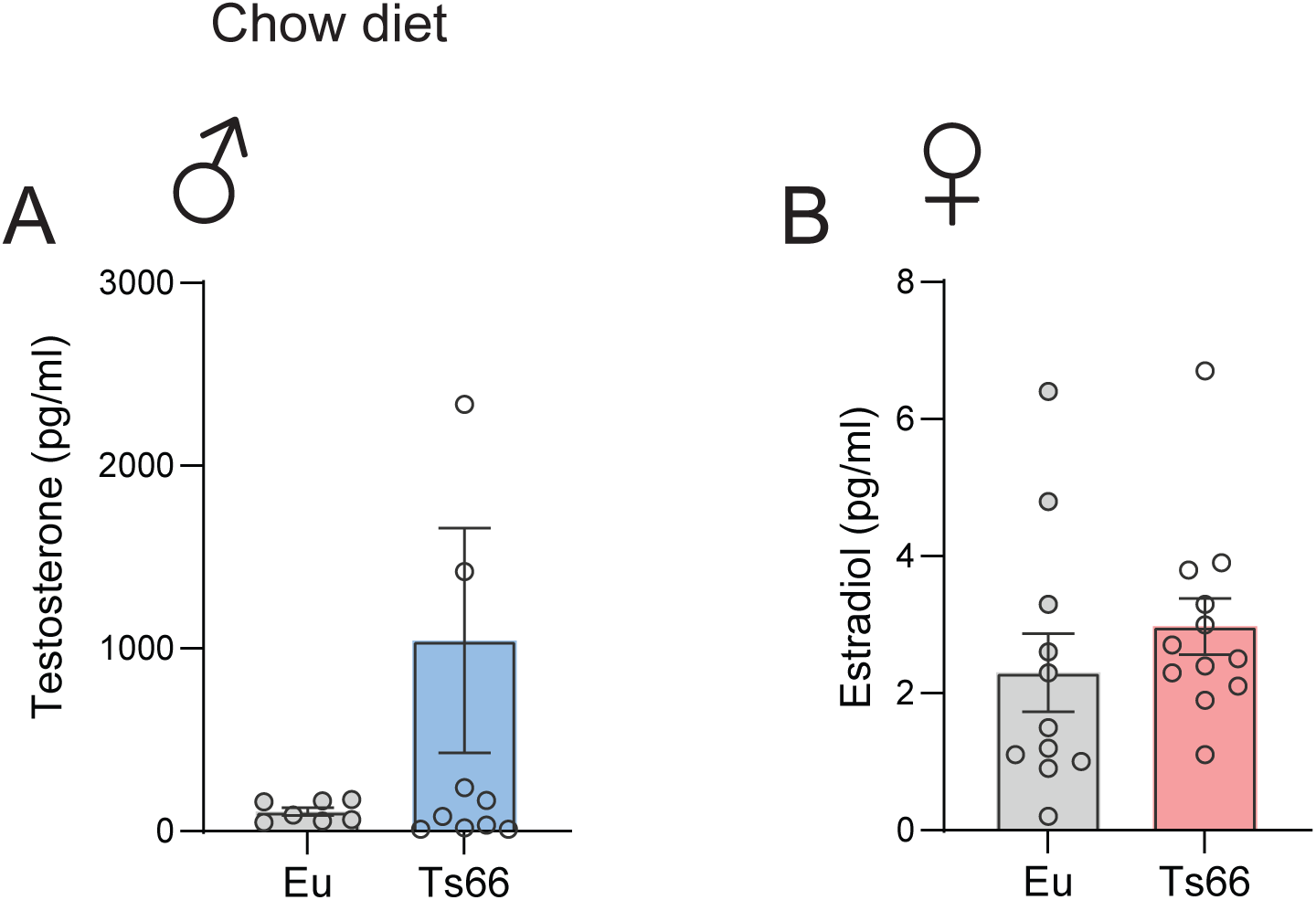
Sex hormone levels in chow-fed mice. (A) Serum testosterone male euploid and Ts66 mice. (B) Serum estradiol in female euploid and Ts66 mice. Male Sample size: Eu = 7; Ts66 = 10. Female sample size: Eu = 11; Ts66 = 12.

**Fig 5 – figure supplement 1.**
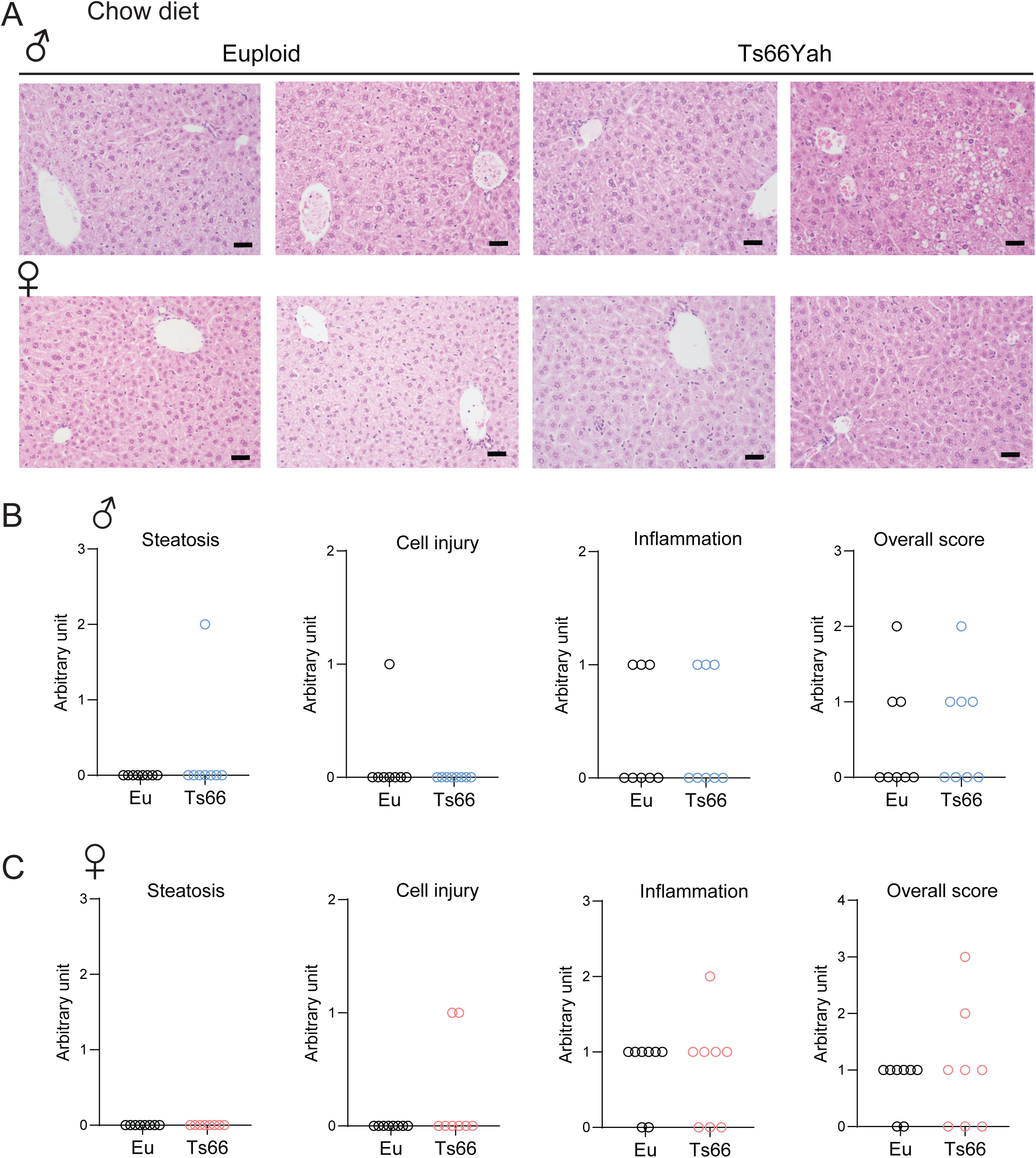
Liver histology and pathology scoring in Ts66Yah mice. **(A)** Two representative H&E-stained liver histology sections from male (top panel) and female (bottom panel) Ts66 mice and euploid (Eu) controls. Scale bar = 100 µM. **(B-C)** Histological assessment for steatosis, cell injury, inflammation, and overall score in male (B) and female (C) Ts66 mice and euploid controls. Sample size for chow-fed male (Eu = 8; Ts66 = 8) and female (Eu = 8; Ts66 = 8) mice.

**Figure 7 – figure supplement 1.**
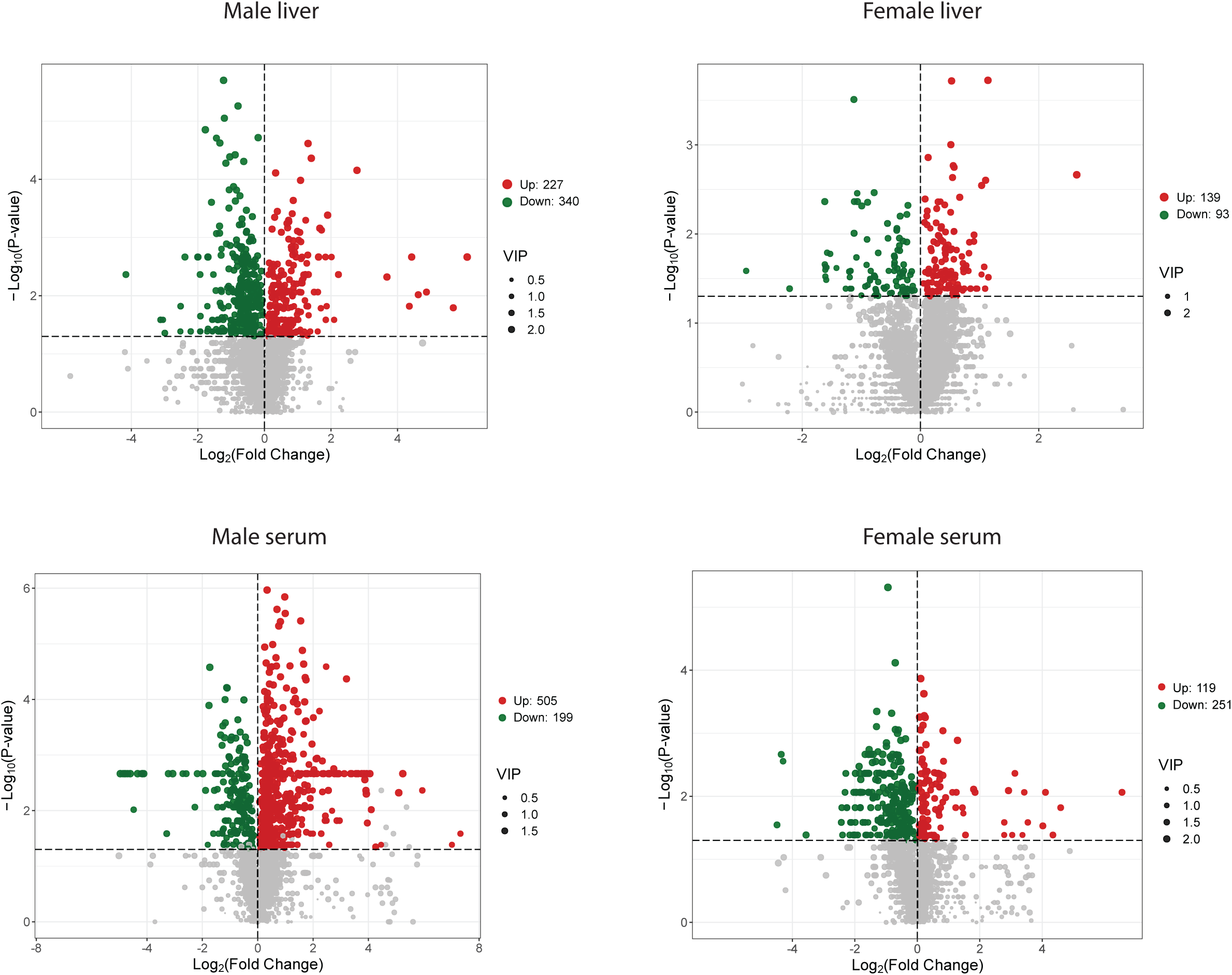
Differential metabolites found in the liver and serum of Ts66Yah male and female mice. Volcano plots showing differential metabolites up- and down-regulated in Ts66 male liver, female liver, male serum, and female serum. *n* = 6 per genotype. VIP, Variable Importance in Projection. VIP scores provide a quantitative measure of a metabolite’s discriminatory power between different groups. Metabolites with a VIP score of 1.0 or greater are considered significant.

**Figure 8 – figure supplement 1.**
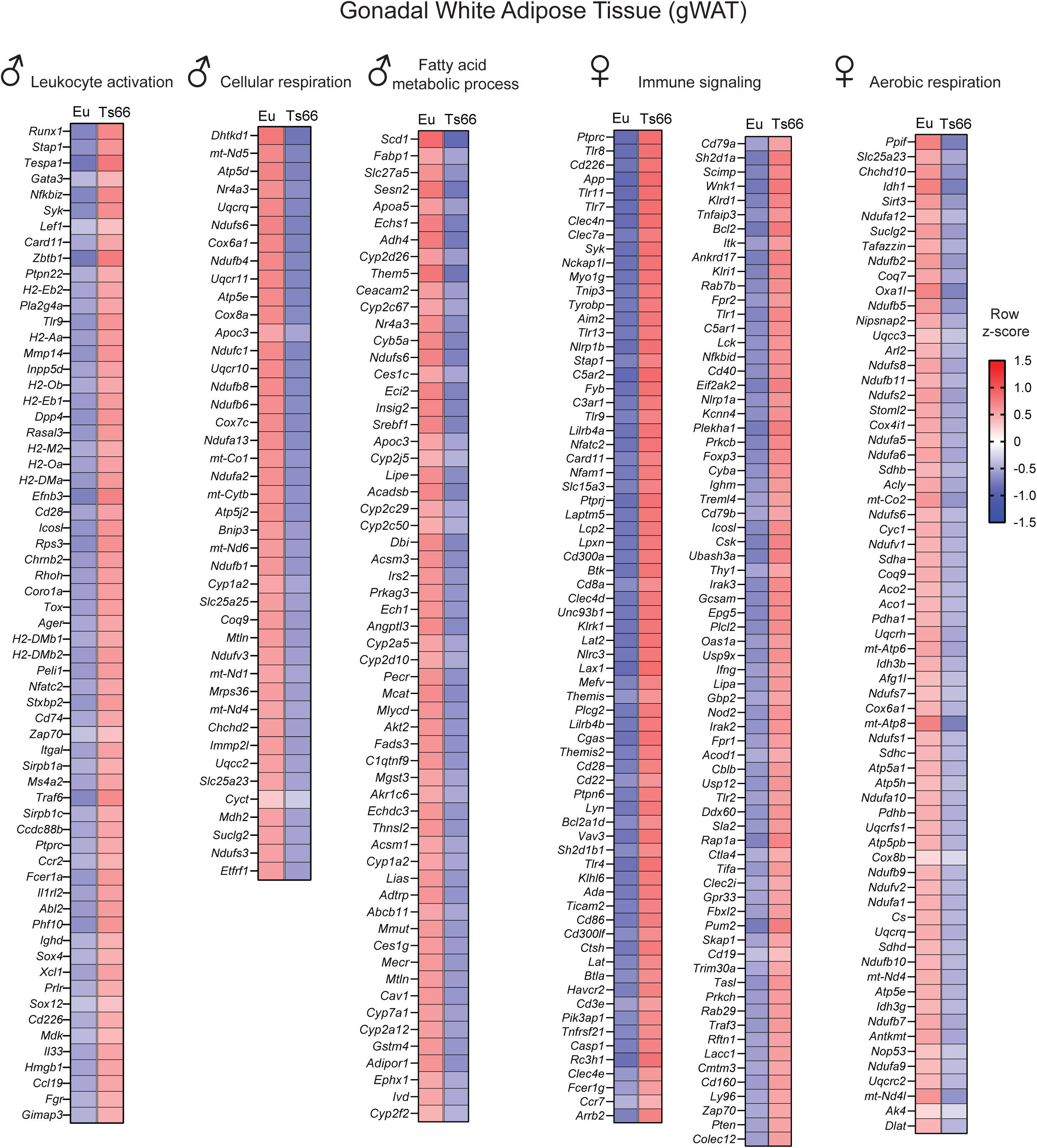
Differentially expressed genes (DEGs) involved in leukocyte activation, cellular respiration, fatty acid metabolic process, immune signaling, and aerobic respiration that are up- or down-regulated in the gonadal (visceral) white adipose tissue (gWAT) of Ts66Yah male and female mice.

**Figure 8 – figure supplement 2.**
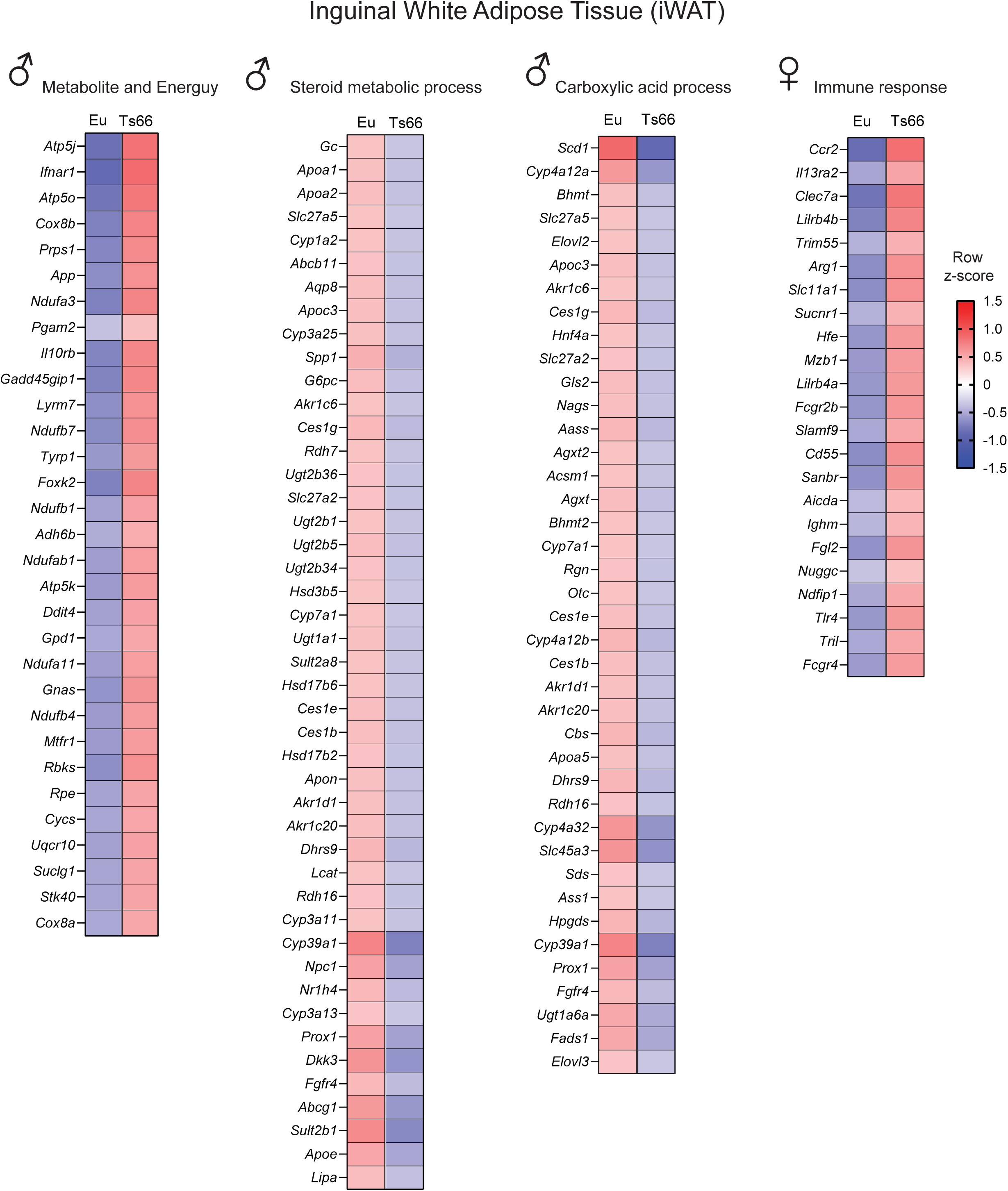
Differentially expressed genes (DEGs) involved in metabolite and energy, steroid metabolic process, carboxylic acid process, and immune response that are up- or down- regulated in the inguinal (subcutaneous) white adipose tissue (iWAT) of Ts66Yah male and female mice.

**Figure 8 – figure supplement 3.**
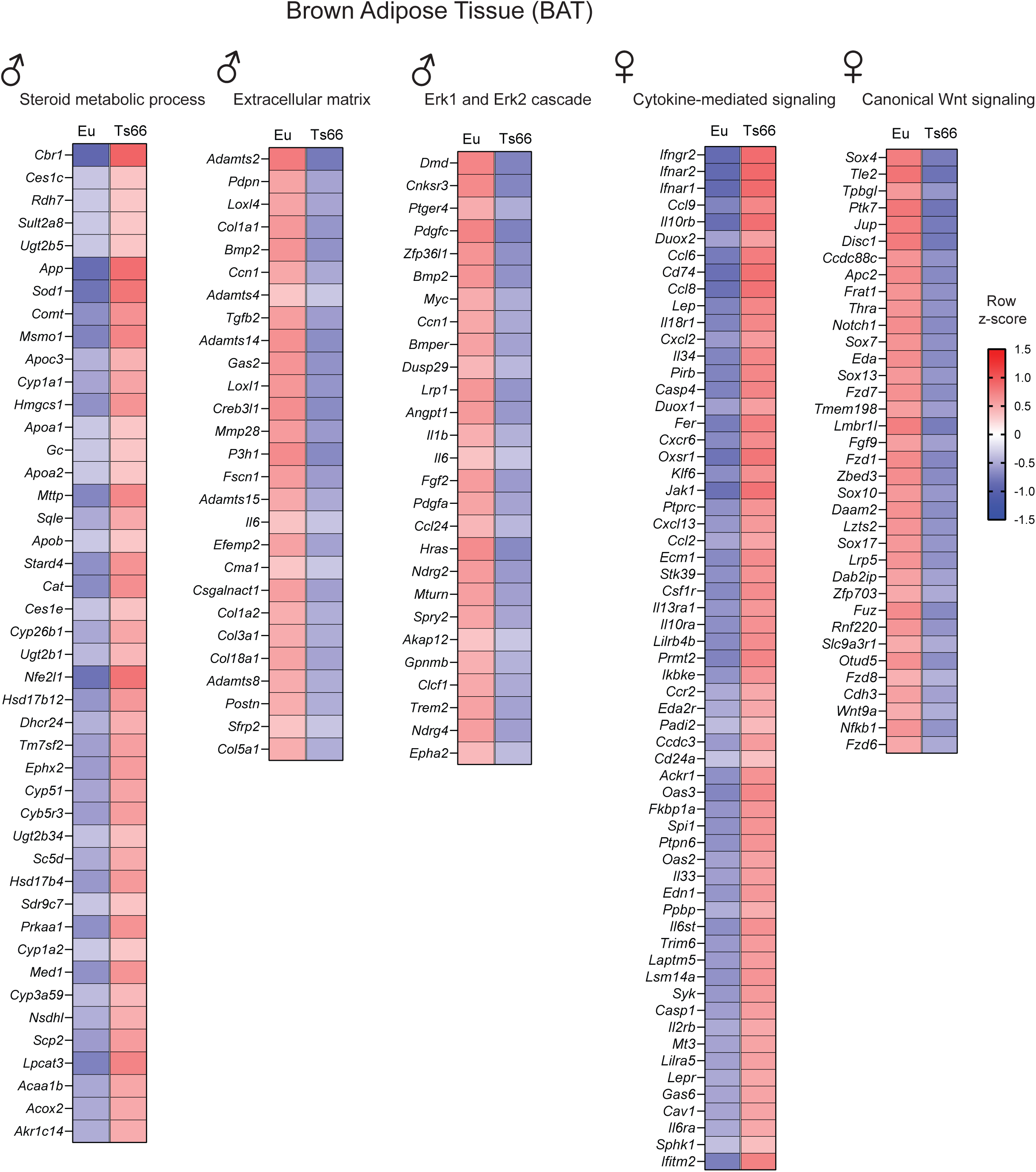
Differentially expressed genes (DEGs) involved in steroid metabolic process, extracellular matrix, Erk1 and Erk2 cascade, cytokine-mediated signaling, and canonical Wnt signaling that are up- or down-regulated in the brown adipose tissue (BAT) of Ts66Yah male and female mice.

**Figure 8 – figure supplement 4.**
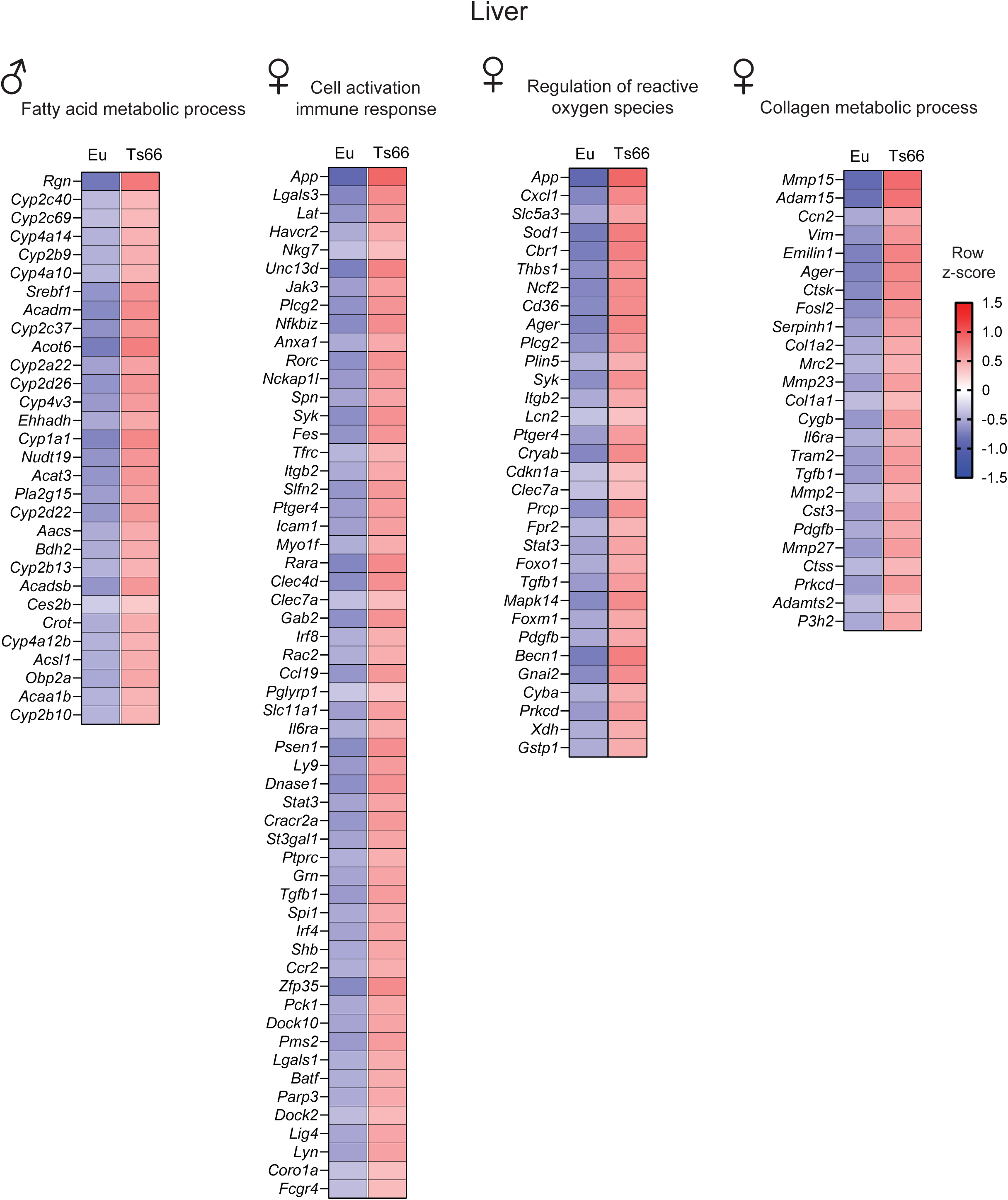
Differentially expressed genes (DEGs) involved in fatty acid metabolic process, cell activation and immune response, regulation of reactive oxygen species, and collagen metabolic process that are up- or down-regulated in the liver of Ts66Yah male and female mice.

**Figure 8 – figure supplement 5.**
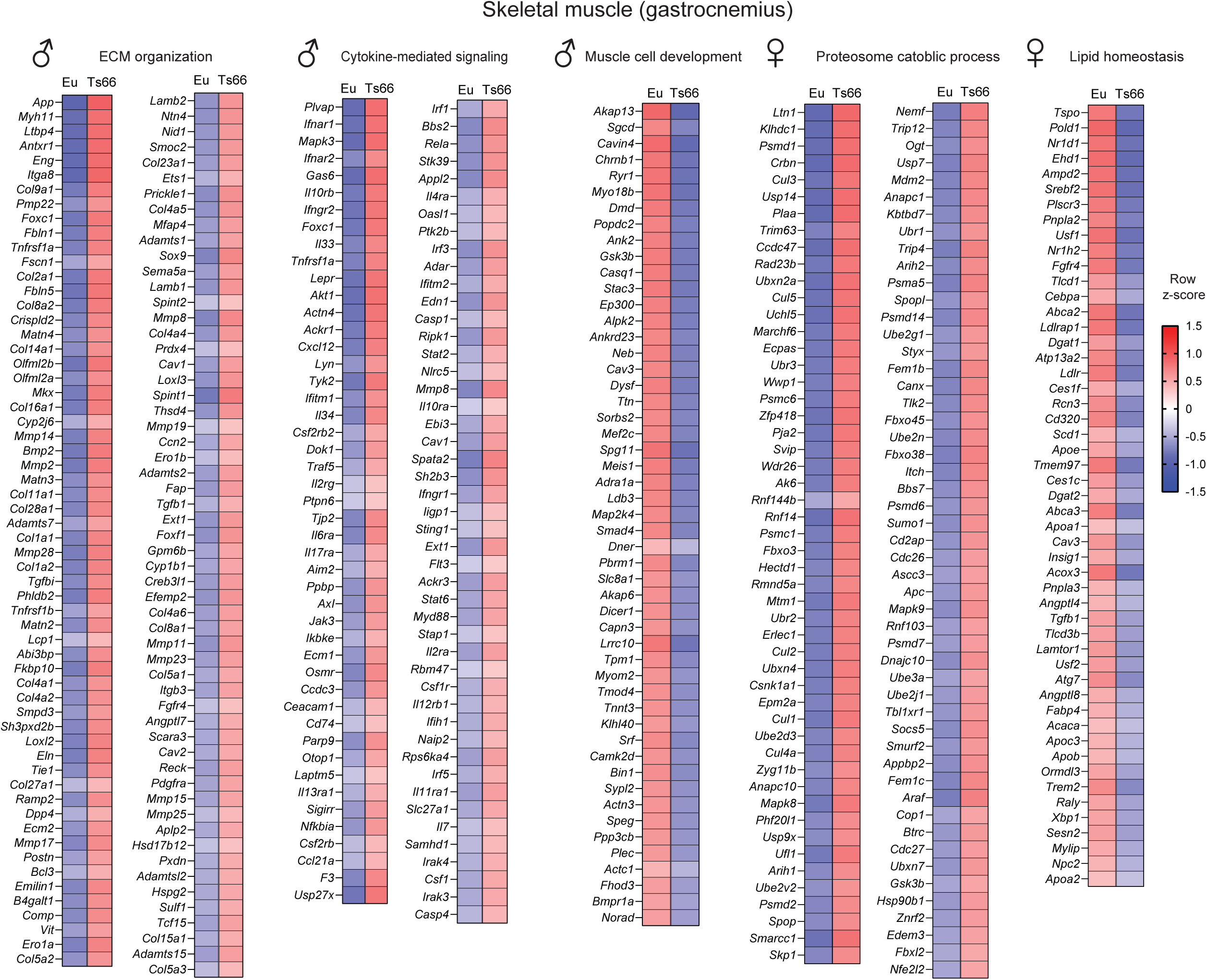
Differentially expressed genes (DEGs) involved in extracellular matrix (ECM) organization, cytokine-mediated signaling, muscle cell development, proteosome catabolic process, and lipid homeostasis that are up- or down-regulated in the skeletal muscle (gastrocnemius) of Ts66Yah male and female mice.

**Figure 9 – figure supplement 1.**
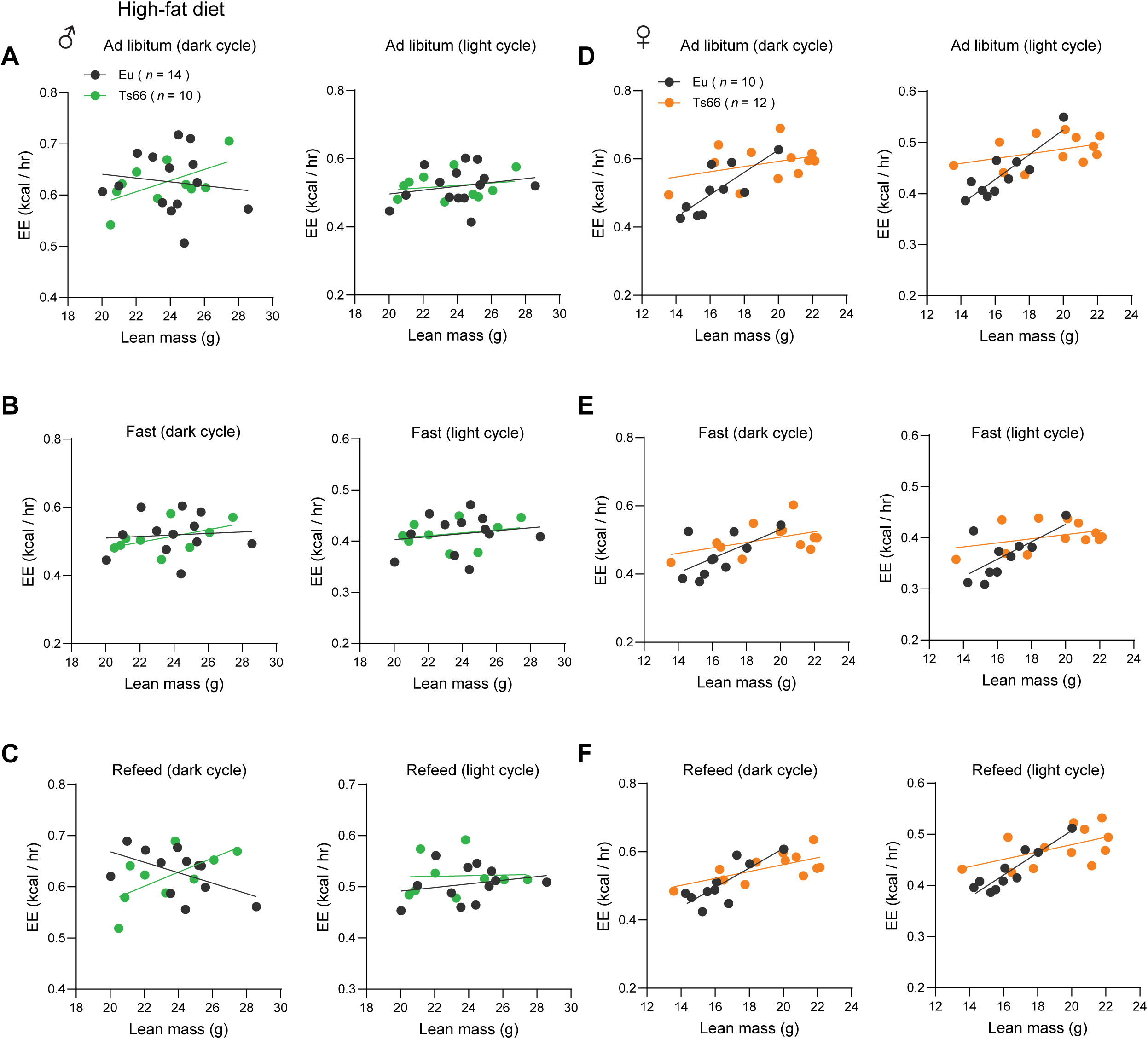
ANCOVA analysis of energy expenditure in HFD-fed mice where lean mass is used as a covariate. ANCOVA analysis of male euploid (Eu) and Ts66 mice across the circadian cycle (dark and light) in *ad libitum* fed (A), fasted (B), and refed (C) states. ANCOVA analysis of female euploid and Ts66 mice across the circadian cycle (dark and light) in *ad libitum* fed (D), fasted (E), and refed (F) states. Male Sample size: Eu = 14; Ts66 = 10. Female sample size: Eu = 10; Ts66 = 12.

**Figure 9 – figure supplement 2.**
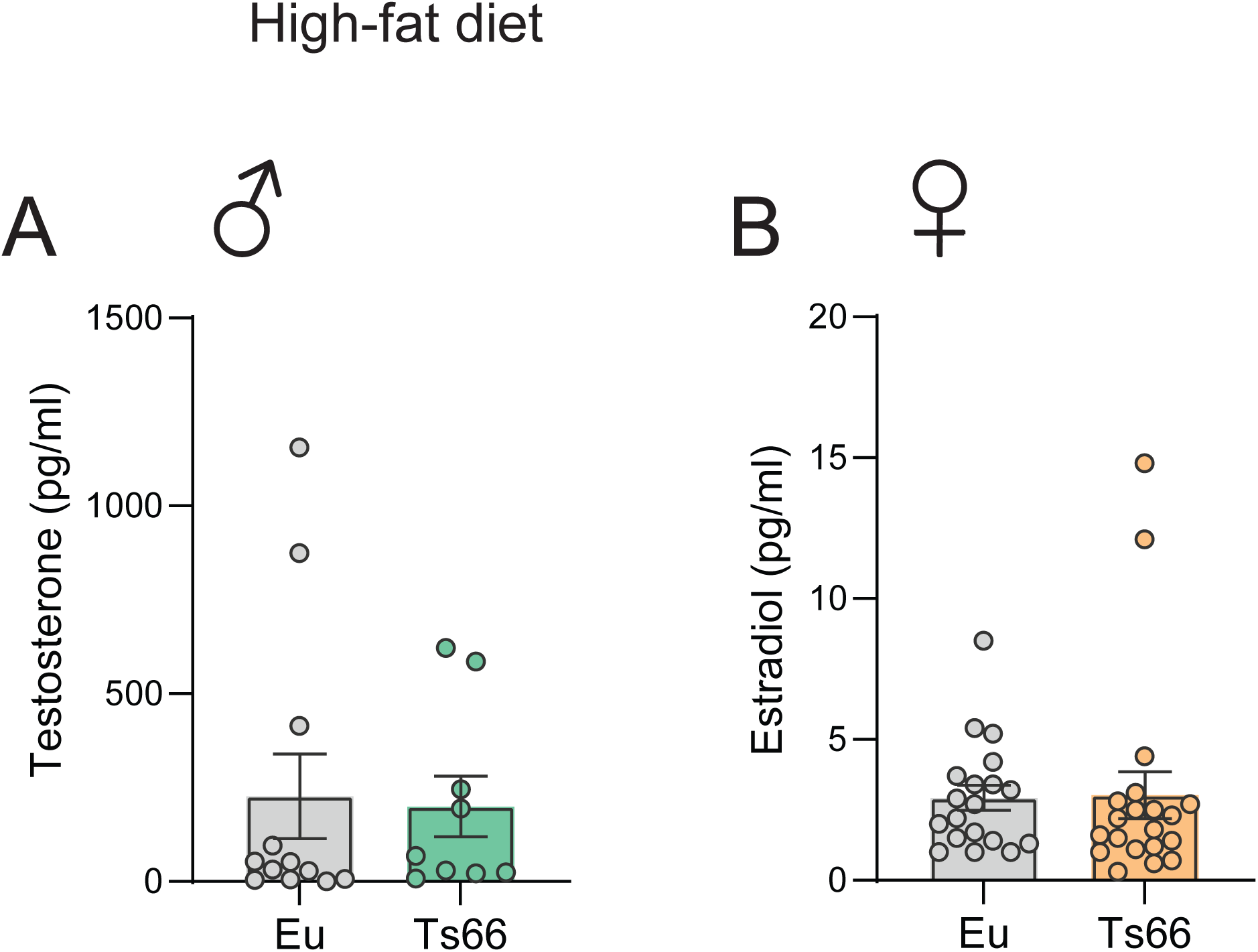
Sex hormone levels in HFD-fed mice. (A) Serum testosterone male euploid and Ts66 mice. (B) Serum estradiol in female euploid and Ts66 mice. Male Sample size: Eu = 12; Ts66 = 11. Female sample size: Eu = 19; Ts66 = 20.

**Figure 9 – figure supplement 3.**
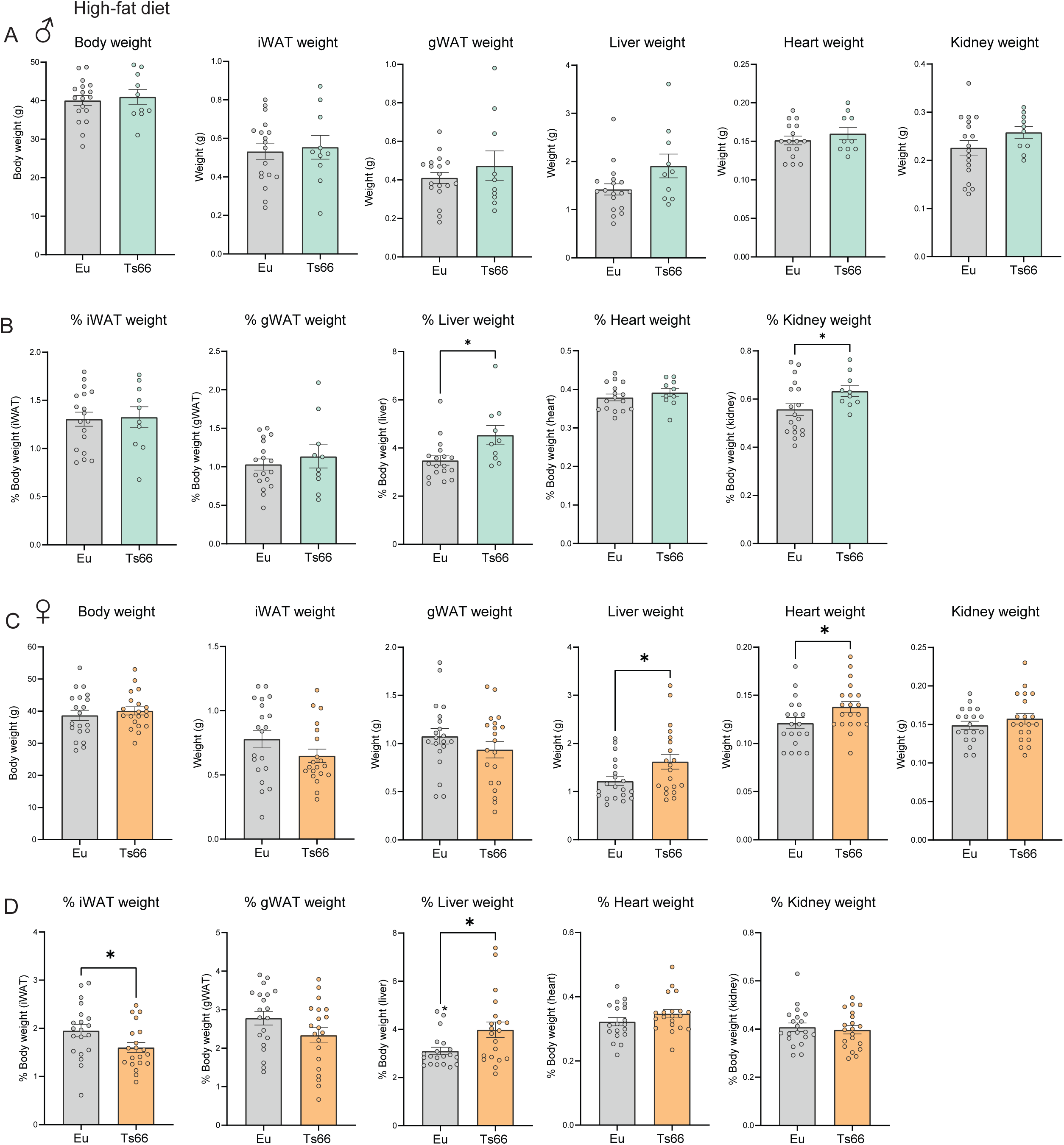
Body and tissue weights of HFD-fed mice at termination of study. Tissues were collected from HFD-fed male and female mice at 23 and 24 weeks of age, respectively. Body weights and the absolute (A and C) and relative (B and D; % of body weight) weights of iWAT, gWAT, liver, heart, and kidney in euploid (Eu) and Ts66 male (A-B) and female (C-D) mice. gWAT, gonadal white adipose tissue; iWAT, inguinal white adipose tissue. Sample size for male mice: Eu = 18; Ts66 male = 10. Sample size for female mice: Eu = 20; Ts66 = 20. All data are presented as mean ± SEM. * *P*<0.05 (two-tailed student *t*-TEST)

**Figure 13 – figure supplement 1.**
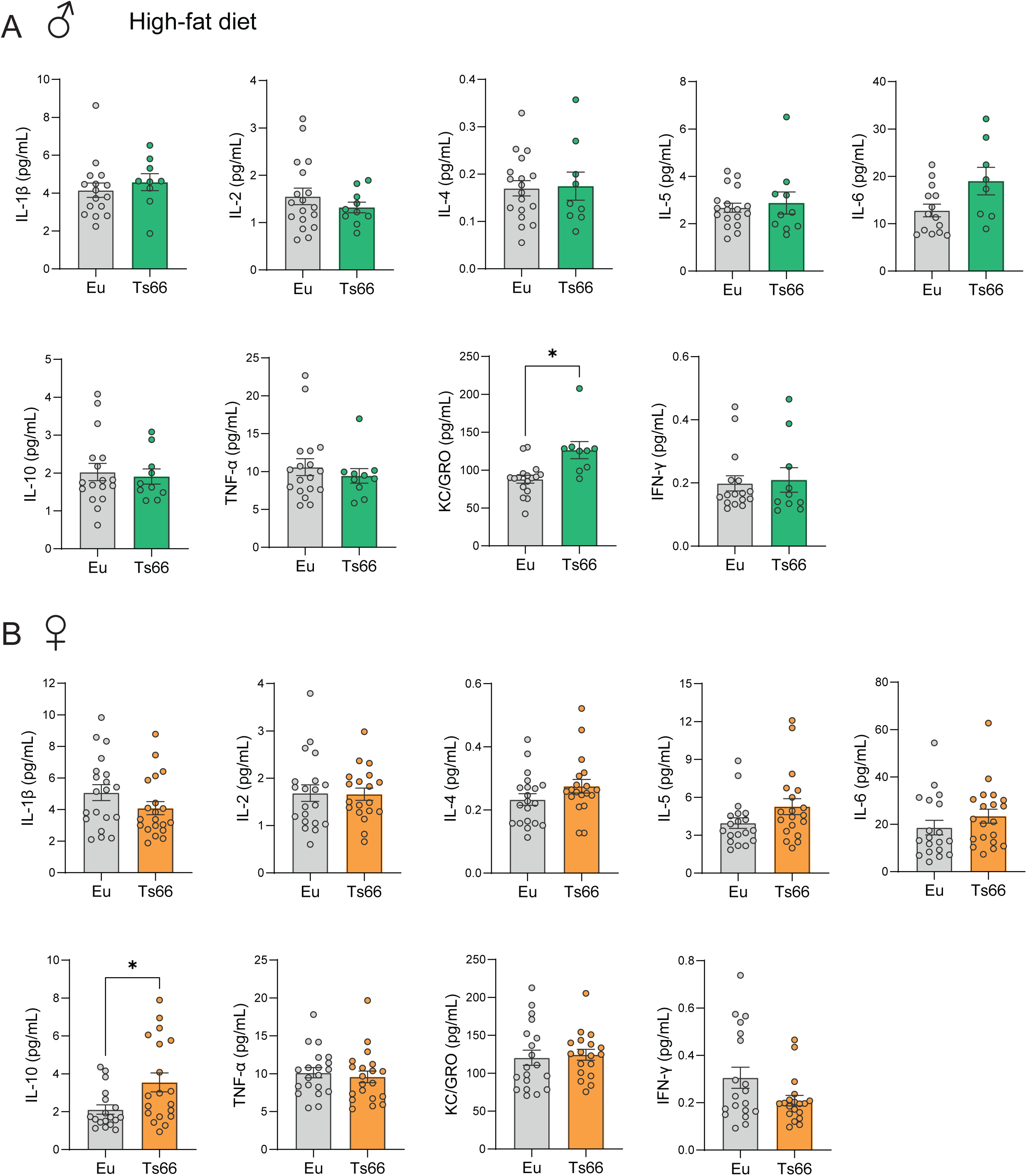
Impact of high-fat diet on serum proinflammatory cytokine profile in Ts66Yah mice. **(A-B)** Multiplex profiling of serum IL-1β, IL-2, IL-4, IL-5, IL-6, IL-10, TNF-α, KC/GRO (also known as CXCL1), and INF-γ in male (A) and female (B) Ts66 mice and euploid (Eu) controls. Sample size for male (Eu = 14-18; Ts66 = 9-10) and female (Eu = 19-20; Ts66 = 19-20) mice.

## Notes

### Competing Interest Statement

The authors have declared no competing interest.

